# Atypical phosphatases drive dissolved organic phosphorus utilization by phosphorus-stressed phytoplankton in the California Current Ecosystem

**DOI:** 10.1101/2025.04.09.648040

**Authors:** Viktoria Steck, Robert H. Lampe, Simran Bhakta, Kimberly Camarena Marrufo, Jamee C. Adams, Eesha Sachdev, Kiefer O. Forsch, Katherine A. Barbeau, Andrew E. Allen, Julia M. Diaz

**Affiliations:** Geosciences Research Division, Scripps Institution of Oceanography, University of California San Diego, La Jolla, CA 92093, United States; Current address: Marine Biogeochemistry Division, GEOMAR Helmholtz Centre for Ocean Research Kiel, D-24148 Kiel, Germany; Integrative Oceanography Division, Scripps Institution of Oceanography, University of California San Diego, La Jolla, CA 92093, United States; Microbial and Environmental Genomics, J. Craig Venter Institute, La Jolla, CA 92037, United States

## Abstract

In the ocean, dissolved organic phosphorus (DOP) supports the health and productivity of marine phytoplankton, a phenomenon most often investigated under inorganic phosphate (Pi) scarcity. However, microbial DOP acquisition occurring in Pi replete ocean environments remains poorly understood. Here, we conducted a combination of nutrient addition experiments, alkaline phosphatase (AP) rate measurements, and metatranscriptomics analyses along an onshore-to-offshore gradient in the California Current Ecosystem (CCE), a relatively Pi-rich upwelling region. We found that AP activity (APA) and eukaryotic transcripts for DOP utilization were present throughout the CCE. In bottle incubations, APA was upregulated in response to iron (Fe) and nitrogen (N) additions. Major contributors to these trends included atypical alkaline phosphatases (AP_aty_) of diatoms in the coastal upwelling area, and unclassified non-cytoplasmic P-diesterases (PDE_nc_) of multiple eukaryotic taxa in the offshore regime. APA and gene expression dynamics were not coupled to phytoplankton growth, suggesting that these organisms experience underlying P stress, a state of cellular metabolism caused by Pi scarcity, even in regions primarily growth-limited by other elements. While AP_aty_ and PDE_nc_ were highly abundant among the microbial community phosphatase pool, these genes were missing from a widely used annotation database, highlighting the importance of manual curation for the detection of these atypical and unclassified proteins. Altogether, these results emphasize the functional diversity of phosphatases sustaining microbial community health in diverse and productive marine habitats.

## Introduction

Phosphorus (P) is an essential nutrient for marine phytoplankton, which generate roughly half of the global net primary production [1]. Dissolved organic phosphorus (DOP) constitutes the majority of available P in the world’s ocean gyres [2]. Previously thought to be utilized only during inorganic phosphate (Pi or PO_4_ ^3-^) scarcity, DOP is now recognized to support marine microbial nutrition and primary productivity across various ocean environments [2], including nutrient-rich coastal areas [3–5]. However, the mechanisms governing microbial DOP utilization across different trophic regimes are poorly understood.

Marine microorganisms produce hydrolytic enzymes to access DOP, which often require metal cofactors to function and thus link the biogeochemical P cycle with trace metals [2]. One widespread enzyme family are alkaline phosphatases (APs), containing zinc (Zn) in the isoform PhoA [6] and iron (Fe) in PhoX and PhoD [7,8], plus unusual metals such as Co [9]. Indeed, microbial AP activity (APA) was found to be limited by environmental Fe or Zn availability in North Atlantic plankton communities [10,11]. DOP also supports primary productivity in Pacific subtropical gyres [12,13], albeit with overall lower APA than in the Atlantic [10], potentially due to less trace metal inputs through dust fluxes [14]. While one study in the North Pacific detected that Fe-limited *Trichodesmium* colonies produce Zn-dependent *phoA* [15], the relationship between P nutrition and trace metals in the Pacific remains unclear.

Ambient DOP consists of P-esters (P-O-C bonds, 80-85% of bulk DOP; e.g. P-sugars, P-lipids, nucleotides), P-anhydrides (P-O-P, 8-13%; e.g. nucleoside di-and triphosphates, pyro-P, poly-P), and phosphonates (C-P, 5-10%; e.g. phosphonolipids, small molecule metabolites) [16]. In order to access this chemically diverse nutrient reservoir, phytoplankton need to employ a large functional variety of different phosphatase enzymes [2]. Besides APs, which hydrolyze P-monoesters [10], several classes of P-diesterases (PDE) and-triesterases (PTER) are utilized, many of which are responsive to Pi availability [17,18]. More recently, P-anhydrides and phosphonates were found to be bioavailable to marine plankton [19–24]. The P-anhydride degrading proteins nucleoside triphosphatase (5’-NT) and inorganic pyrophosphatase (*ppa*) are implicated in the growth of diatoms under low Pi [21], and some nucleoside diphosphate (NUDIX) hydrolases upregulated under Pi scarcity [18]. Together with polyphosphatases (*ppx*), 5’-NTs are also used by heterotrophs to acquire P and/or carbon-containing byproducts [25,26].

Here, we provide physiological and transcriptional insights into DOP utilization within natural mixed microbial assemblages experiencing multifold nutrient stressors between the North Pacific Subtropical Gyre (NPSG) and the California Current Ecosystem (CCE). The CCE is an eastern boundary current upwelling system with a high variability of ecological zones and strong diatom-driven productivity [27]. Large parts of the central CCE are perpetually nitrogen (N)-limited, but experience seasonal and intermittent Fe-limitation especially in upwelling and transition zones [28–31]. The ability to metabolize DOP buffers phytoplankton against nutrient stress [32], and further deciphering how DOP supports marine microorganism communities in the face of environmental changes is vital for improving primary production and climate predictions. This knowledge is especially important in light of key disturbances in Pacific Ocean ecosystems, including altered nutrient stoichiometry [33], marine heatwaves [27], water column hypoxia, and ocean acidification [34].

## Methods

### Field sampling

Sampling was conducted on two expeditions in the California Current: June 2020 aboard the R/V Sally Ride (SR2003), and July–August 2021 aboard the R/V Roger Revelle (P2107). P2107 was part of the CCE Long-Term Ecological Research (LTER) program and used drifters for quasi-Lagrangian tracking of discrete water parcels [35]. Each series of measurements during continuous tracking of one water mass was denoted a Lagrangian cycle. For comparison to other studies from P2107, station 1 corresponds to cycle 1, station 2 to cycle 2, station 3 to the California Current Transect, and station 4 to cycle 3 (**Fig. 1**). SR2003 served one station at 32.51304 °N, 118.21386 °W. On P2107, seawater for dissolved inorganic nutrients was collected using a SBE911+ CTD system mounted in a 24-place 10 L Niskin rosette with an SBE32 carousel (Sea-Bird Electronics). Trace metal clean seawater was collected using single acid-cleaned, Teflon-coated 12 L GO-Flo bottles (General Oceanics) mounted on non-metallic hydroline (Amsteel) triggered with acid-cleaned Teflon messengers designed by Ken Bruland (SR2003), or 5 L X-Niskin bottles (Ocean Test Equipment) mounted on a powder-coated rosette deployed on kevlar-coated hydrowire (SpaceLay) (P2107). Subsurface sampling was conducted at the depth below the top of the nitracline, determined by the nitrate-sensors ISUS (SR2003) or SUNA (P2107) mounted on the CTD rosette, which did not always correspond to the depth of maximum Chl-*a* (**Fig. S1**). This region was previously identified to feature higher Fe stress due to mismatched stoichiometric supply of N and Fe from diapycnal mixing [30].

**Figure 1.**
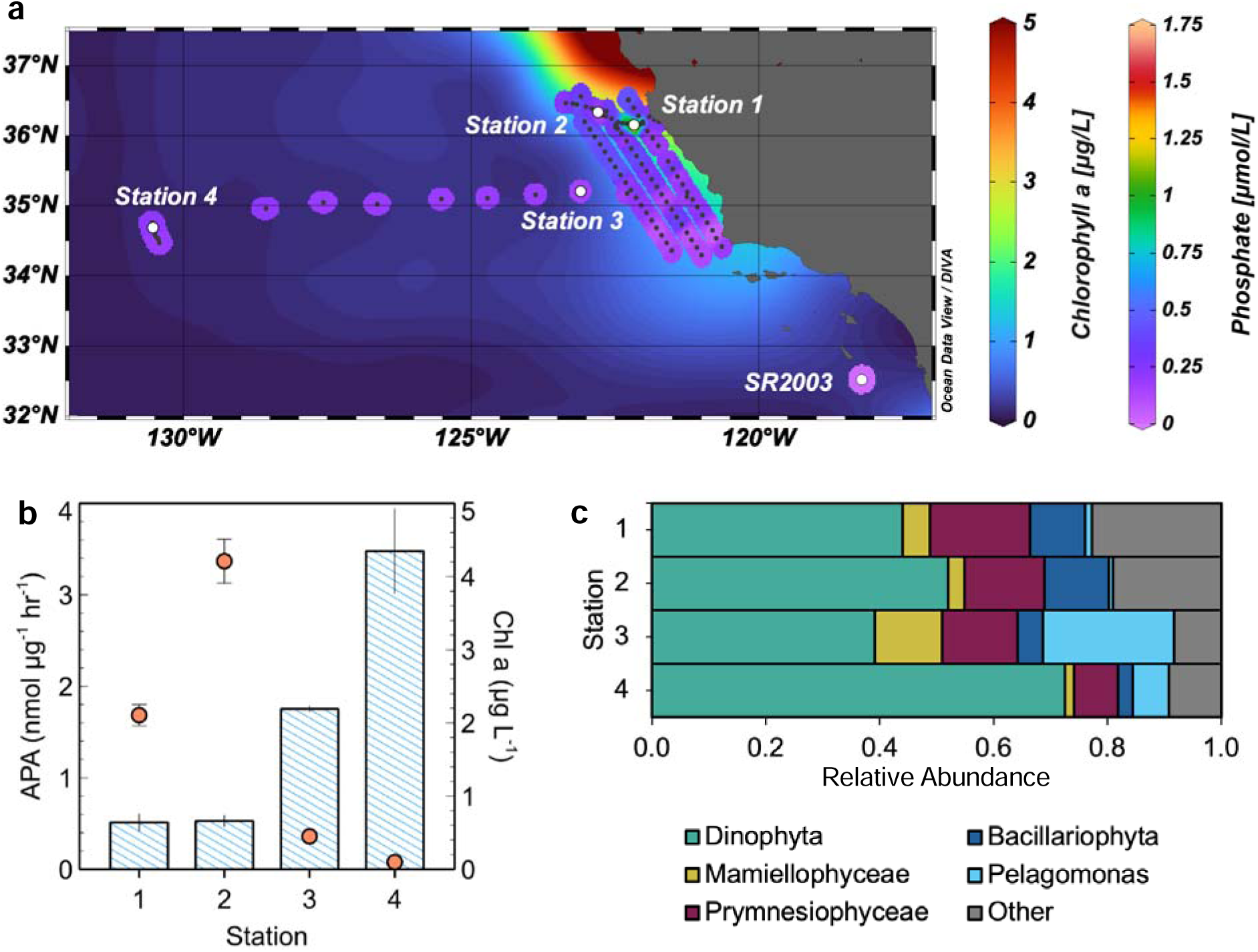
Cruise track, ambient biogeochemical parameters, and eukaryotic community composition in the California Current Ecosystem. **a)** Surface concentrations of inorganic phosphate (PO_4_ ^3-^; gridded projection) during the survey transect (black dots) and at stations 1-4 (white circles) on the backdrop of satellite-derived surface chlorophyll-*a* (Chl-*a*; contoured background). **b)** *In situ* alkaline phosphatase activity (APA; blue bars) and chlorophyll-*a* (Chl-*a*; orange circles) at stations 1 (15 m), 2 (20 m), 3 (40 m), and 4 (140 m). Error bars represent one standard error of the mean, *n* = 3. **c)** Average relative abundances of major eukaryotic groups at ambient conditions inferred from poly(A)-selected mRNA (metatranscriptome), obtained at stations 1 (15 m), 2 (20 m), 3 (40 m), and 4 (140 m). The average abundances were derived from the relative abundances of eukaryotic-assigned transcripts per million normalized reads (TPM).

### Field incubations

Bottle incubations at stations 1-4 (**Fig. 1**) were set up under positive pressure with HEPA-filtered air in a trace metal clean plastic bubble (SR2003) or UNOLS trace metal clean van (P2107). Metal stocks were prepared as 1.5 μM (CoCl_2_·6H_2_O) or 15 μM (FeCl_3_·6H_2_O, ZnSO_4_·7H_2_O, MnSO_4_·6H_2_O) in 2% HCl. Nutrient stocks (NaNO_3_, NH_4_Cl) were prepared as 15 mM in H_2_O, treated with Chelex for 24 hours, and 0.2 µm-filtered prior to use. Polycarbonate bottles, previously cleaned with 1% Citranox followed by 10% HCl (optima grade, Fischer Scientific) for at least two weeks each, were rinsed three times, filled with seawater, and spiked with combinations of the trace metals (final concentrations): Fe (5 nM), Zn (5 nM), Co (0.5 nM), and nitrogen (+N) as both NO_3_ ^2-^ (5 mM) and NH_4_^+^ (5 mM). At P2107 stations 1-4, 1 L polycarbonate bottles (n = 12) were treated in triplicate with +Fe, +N, +Fe+N, including an unamended control. At SR2003, 60 mL polycarbonate bottles (n = 24) were treated in triplicate with +Fe, +Zn, +Co, +N, +Fe+N, +Zn+N, +Co+N, plus an unamended control. Bottles were incubated on-deck in circulating seawater from the ship’s underway and covered with neutral density screening to mimic in situ temperature and light, as confirmed with a PAR sensor, and sampled after 48h.

### Environmental measurements

Samples were analyzed for chlorophyll-*a* [36], cell counts [20], dissolved inorganic nutrients [37], soluble reactive phosphorus [38], dissolved organic phosphorus [39], dissolved iron [40,41], APA [20] and metatranscriptomics [42] using previously established protocols (see *Supplementary Material, Standard Procedures*).

### Statistical analyses

Statistical analyses of chlorophyll-*a*, cell counts, and APA were performed in JMP Pro (15.2.0) using repeated-measures ANOVA followed by Tukey’s Honest significant difference (HSD) test. Statistical analyses of mRNA (fold-changes) were performed in R using the DESeq2 v1.42.1 Wald test adjusted for multiple testing via the Benjamini and Hochberg method [43]. *P*-values and adjusted *p*-values <0.05 were considered significant.

## Results

### Chlorophyll-*a* and phytoplankton abundance

The P1207 cruise track followed an onshore-to-offshore gradient in productivity from coastal upwelling in the CCE toward the NPSG (**Fig. 1a**, **Table 1**). Inshore, station 1 featured high chlorophyll-*a* (Chl-*a*) and phytoplankton cell concentrations, which indicated the onset of a bloom. Quasi-Lagrangian tracking of this discrete water parcel showed that Chl-*a* doubled while phytoplankton cell concentration remained similar at station 2. Biomass declined at Station 3, in the transition zone, and further at station 4, located at the border of the oligotrophic NPSG. Similarly, the chlorophyll maximum deepened going from onshore to offshore (**Fig. S1**). During the SR2003 cruise, located south of P2107 in a non-upwelling area, ambient phytoplankton cell concentrations averaged 4,920 cells mL^-1^, comparable to stations 3-4.

**Table 1:**
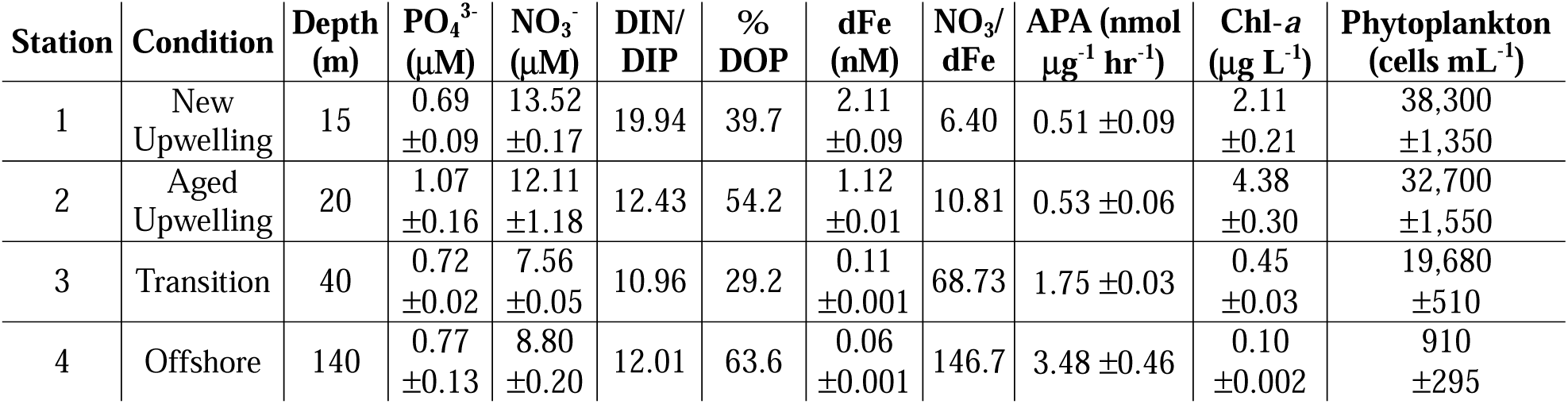
***In situ* environmental conditions at stations 1-4.** Error values represent one standard error of the mean, *n* = 3. % DOP is presented as fraction of TDP. DIN = nitrate + nitrite + ammonia.

### Dissolved nutrients

At the surface (∼2 m), the upwelling event at station 1 injected more than 1 μM phosphate into the euphotic zone, which was quickly removed to 0.29-0.52 μM (**Fig. 1a**). Phosphate measured at depth in waters used for incubation experiments (15-140 m) reached 0.69-1.07 μM, and was highest at station 2 (**Table 1**; **Figs. S2-3**). At the surface, nitrate concentrations generally followed the trend of phosphate, except for depletion below the detection limit at offshore station 4 (**Fig. S2**). Nitrate was considered replete (>7 μM) at the incubation sampling depths of stations 1-4, but decreased slightly from onshore to offshore (**Table 1**). Dissolved inorganic N:P ratios fell below the Redfield ratio [44] at stations 2-4 (N:P = 10.96-12.43), and above Redfield at station 1 (N:P = 19.94; **Table 1**). DOP made up a maximum of 98% of the ambient total dissolved phosphorus (TDP) pool at the surface of the cruise transect, and between 29-64% at depths of stations 1-4 (**Table 1**; **Fig. S3**). Dissolved iron (dFe) in the upwelled water mass declined from 2.11 nM at station 1 (15 m) to 1.12 nM at station 2 (20 m), then dropped to sub-nanomolar levels at remaining stations (**Table 1**). The ratio of dissolved nitrate to dFe (NO_3_:dFe), a proxy for Fe limitation in the diatom-dominated CCE, surpassed the threshold of ∼8 μmol/nmol [29] at stations 2-4.

### Alkaline phosphatase activity (APA)

Despite the prevailing Pi-replete environment (>0.7 μM at incubation sampling depths), APA was measurable at all stations (**Fig. 1b**; **Table 1**). Ambient volumetric APA at these depths rose from 26.0 nM day^-1^ at station 1 to 39.9 nM day^-1^ at station 2, then declined to 13.7 nM day^-1^ and 8.2 nM day^-1^ at stations 3-4, respectively (**Fig. S4**). When normalized to Chl-*a*, specific APA increased 4-fold along the on-to offshore continuum, from ∼0.5 nmol μg^-1^ hr^-1^ in costal regimes (stations 1-2) to 1.75 nmol μg^-1^ hr^-1^ at station 3 and 3.48 nmol μg^-1^ hr^-1^ at station 4 (**Table 1**). This trend was even more pronounced for APA normalized to phytoplankton cell abundance, which enhanced 14-fold from station 1 to station 4 (**Fig. S4**). No correlation was found between APA, phosphate concentrations, or N:P ratios (**Fig. S5**).

### Eukaryotic community composition

Dinoflagellates (Dinophyta) constituted the main eukaryotic phytoplankton class at stations 1-4 (**Fig. 1c**) and mostly contained the harmful algal bloomers *Alexandrinum*, *Ceratium* and *Kareniaceae* (**Fig. S6a**). Prymnesiophyceae, especially *Emiliania* and *Phaeocystis*, were the second most abundant eukaryotes, followed by diatoms (Bacillariophyta), the green algae class Mamiellophyceae, and *Pelagomonas*, a small picoeukaryote that dominates deep chlorophyll maxima in this region [45]. After dinoflagellates and prymnesiophytes, diatoms were most prevalent at stations 1-2, mamiellophytes and *Pelagomonas* at station 3, and *Pelagomonas* at station 4 (**Fig. 1c**). Coastal diatoms were mostly centric (65-73%) and comprised chain-forming *Chaetoceros, Thalassiosira,* and *Pseudo-nitzschia*, while the percentage of pennate lineages increased in transition and offshore zones (42-47%; **Fig. S7a**). Mamiellophyceae consisted predominantly of *Bathycoccus*, *Ostreococcus*, and *Micromonas* (**Fig. S6a**). Other eukaryotes included ciliates (3.4-6.1%), cryptophytes (3.2-4.6%), other Stramenopiles (8.5-10.5%) and rhizaria (0.6-0.7%) (**Table S6**). Altogether, eukaryotic community composition differed substantially among the coast (stations 1-2) and offshore (stations 3-4) (**Fig. S8**).

### Functional gene expression

To assess the nutritional status of ambient plankton assemblages, relative transcript abundances of established nutrient stress biomarkers were tracked from onshore to offshore. The N stress genes for nitrate assimilation (NRT2, NR, *nirA*) [46] and alternative N acquisition such as urea uptake or amino acid catabolism (DUR3, *glnA*) [47,48] were elevated at stations 1 and 4 (**Fig. 2a**). The iron starvation induced proteins ISIP1-3 [49–51] and flavodoxin *fldA* [52] peaked at stations 2-3 (**Fig. 2b**). Since diatoms possess two clades of flavodoxin, of which only one is Fe-sensitive [53], their stress markers were plotted separately from the rest of the eukaryotic community (**Fig. S9**). Diatom *fldA* followed the trends of diatom ISIP1-3 being highest at station 2, while Fe stress markers of remaining eukaryotes were highest at station 3. Within the whole community, enriched transcripts for the Zn/Co responsive proteins ZCRPA/B as well as potential Zn/Co transporters of the ZIP family were found at station 2 (**Fig. 2d**) [46,54,55]. Molecular markers for phosphate stress, i.e. the high-affinity Pi transporters *pstS* [46,56] and *SLC34A* [57], the low-Pi-induced AP isoform *phoA* [10], and *SQD1*, which converts phospholipids to sulfolipids under Pi scarcity [58], show a general decrease from station 1 to 4, with the exception of *phoA* at station 1 (**Fig. 2c**). Moreover, all four P stress markers were overrepresented in diatoms at station 2, a trend that was not shared by the remainder of the community (**Fig. S9**).

**Figure 2.**
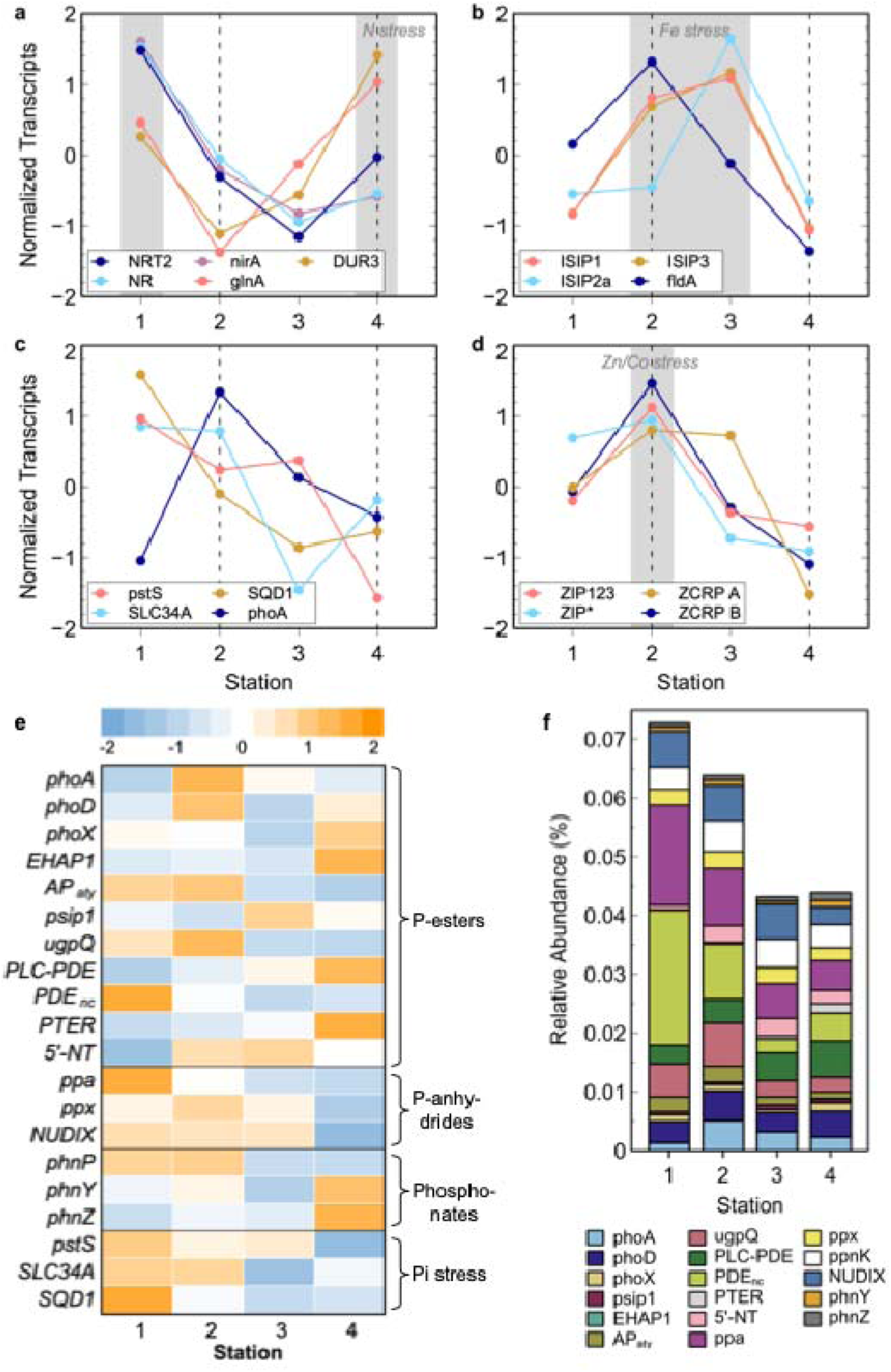
*In situ* transcriptomic signatures of phytoplankton community nutrient stress at stations 1-4. Established stress biomarkers for nitrogen (N; **a**), iron (Fe; **b**), phosphorus (P; **c**) and zinc/cobalt (Zn/Co; **d**) as z-score normalized relative abundances (%) of transcripts per million. Dotted lines indicate stations selected for experimental bioassays with transcriptomics; shaded areas denote inferred nutrient stress. (**e**) Z-score normalized percentages of transcripts involved in P metabolism. (**f**) Relative abundances of DOP hydrolase transcripts. NRT2, nitrate/nitrite transporter; NR, NAD(P)H-nitrate reductase; *nirA*, ferredoxin-nitrite reductase; *glnA*, glutamine synthetase; DUR3, urea-proton symporter; ISIP1-3, iron stress responsive proteins 1-3; *fldA*, flavodoxin A; *pstS*, high-affinity phosphate transport system substrate-binding protein; SLC34A, sodium-dependent phosphate cotransporter; SQD1, UDP-sulfoquinovose synthase; *phoA*, alkaline phosphatase PhoA; ZIP123 and ZIP8, solute carrier family 39 (zinc transporter); ZCRP A/B, zinc/cobalt responsive proteins A/B; *phoD*, alkaline phosphatase PhoD; *phoX*, alkaline phosphatase PhoX; EHAP1, *Emiliania huxleyi* AP1; AP_aty_, atypical alkaline phosphatase; *psip1*, alkaline phosphatase Psip1; *ugpQ*, glycerophosphoryl diester phosphodiesterase; PLC-PDE, PLC-like phosphodiesterase; PDE_nc_, unclassified phosphodiesterase; PTER, phosphotriesterases; 5NT, 5’-nucleotidases; *ppa*, inorganic pyrophosphatase; *ppx*, exopolyphosphatase; *nudH*, (di)nucleoside polyphosphate hydrolase; NUDIX, 8-oxo-dGTP diphosphatase; *phnP*, phosphonate regulator; *phnY*, 2-aminoethylphosphonate dioxygenase; *phnZ*, 2-amino-1-hydroxyethylphosphonate dioxygenase.

A multitude of DOP utilization signatures was present throughout the CCE (**Fig. 2e-f**), including genes involved in the utilization of all major DOP bond classes: P-esters, P-anhydrides, and phosphonates. However, the Kyoto Encyclopedia of Genes and Genomes (KEGG) Orthologs (KOs) commonly used for functional metatranscriptome annotations [59] did not capture several marine microbial APs. These genes fell into two categories (**Tables S3-4**): (1) those that could be assigned to an existing KO based on literature references or BLASTP search, and (2) those that could not. In particular, the latter case included atypical APs (AP_aty_), e.g. atypical PhoA_aty_ of *Emiliania huxleyi* [60,61]; and diverse PDEs. Hence, we compiled AP_aty_ with 12 published and manually annotated homologs of atypical AP sequences stemming from Stramenopiles, Prymnesiophyceae, and green algae (**Table S3**). Two categories of uncharacterized PDEs, containing 20 genes from diatoms, hacrobia, and green algae, were created in a similar manner (**Table S4**): phospholipase C (PLC)-like PDE (3 genes) is involved in the degradation of phosphoinositides, while non-cytoplasmic PDE_nc_ (17 genes) represent secreted or membrane-bound extracellular, dimetal-containing P-diesterases.

PDE_nc_ constituted the largest fraction of DOP-targeting genes observed in the study area (up to 33%), followed by *ppa*, NUDIX, PLC-PDEs, *phoD* and *ugqQ* (**Fig. 2f**). AP_aty_ were within 4-5% of DOP-targeting genes comparable to *phoA* (2-9%) and *phoX* (1-4%). All phosphatases had broad taxonomic representation; most PDEs were attributed to prymnesiophytes, *ppa* to dinoflagellates, and *ugpQ* to diatoms (**Fig. S10**). Spatially, AP_aty_ was enriched at stations 1-2, *phoA* and *phoD* at station 2, *phoX* at station 4, and the high-affinity phosphatase *psip1* [62] at station 3 (**Fig. 2e,f**). PDE_nc_ was most prevalent at station 1, *ugpQ* at station 2, and PLC-PDEs and P-triesterases (PTER) at station 4. Genes involved in P-anhydride metabolism, i.e. *ppa*, *ppx*, and NUDIX, were abundant throughout all stations, whereas the phosphonate-degrading oxygenases *phnYZ* increased offshore.

### Nutrient-addition incubations

To investigate microbial health and P nutrition under different nutrient stress regimes, on-deck incubations were performed at stations 1-4 with addition of Fe and N. Eukaryotic community metatranscriptomics were conducted from incubations at stations 2 and 4.

After a 48h incubation, phytoplankton growth by Chl-*a* increased at coastal stations 1-2 following +Fe additions (Fe/FeN; **Fig. 3a,b**). In contrast, at the offshore station 4, Chl-*a* was stimulated following +N addition (N; **Fig. 3c**). This growth response to +N was mirrored by a significant increase in phytoplankton cell abundance, while stations 1-2 did not show significant changes in cell counts (**Fig. 3d-f; Fig. S11**). No significant biomass responses to Fe or N additions were detected at station 3 (**Fig. 3g**). For SR2003 in the southern CCE, phytoplankton abundances were significantly elevated within Zn or Fe-amendments, with or without N (**Fig. S12**).

**Figure 3.**
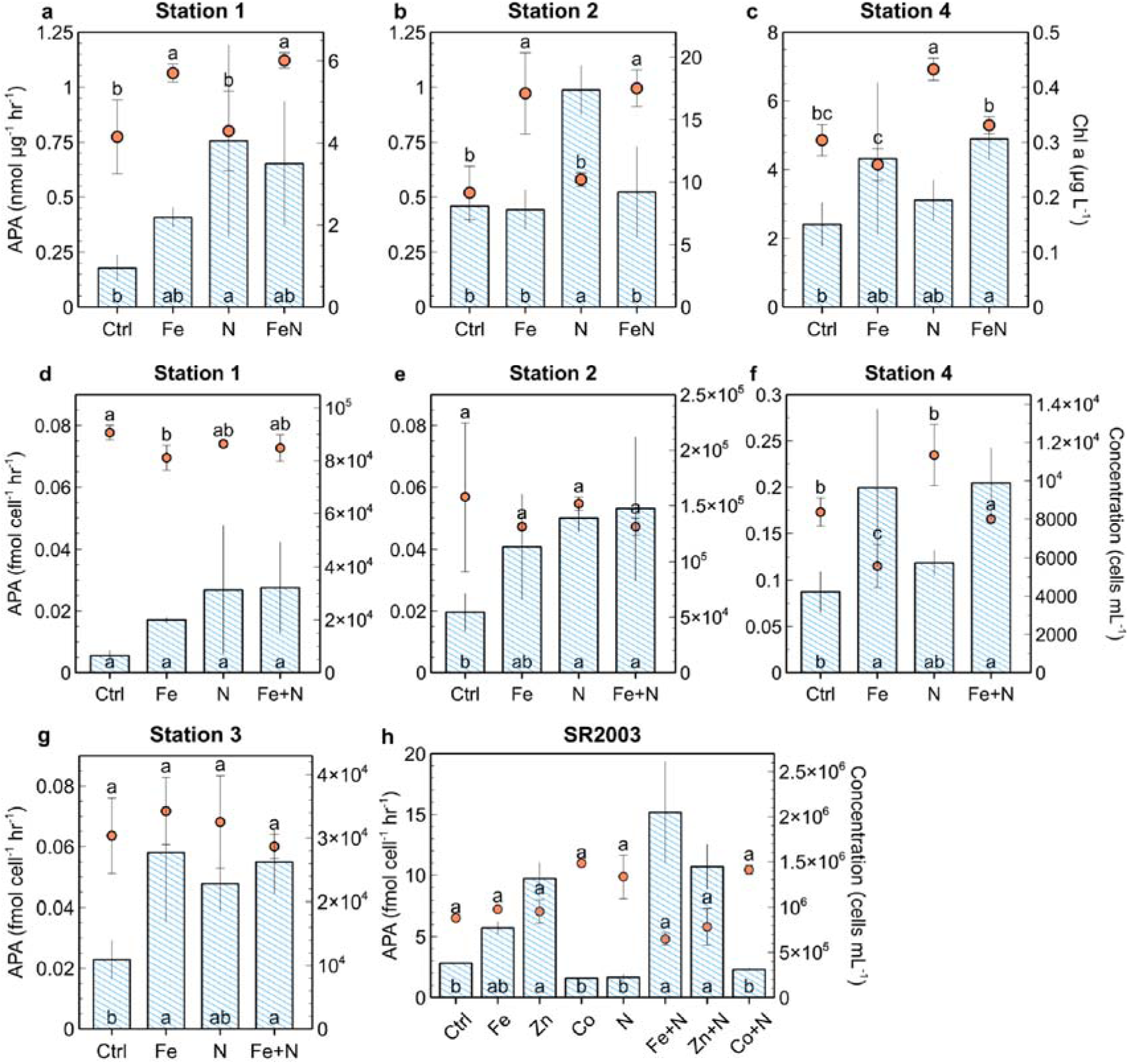
Microbial growth and alkaline phosphatase activity (APA) at experiment locations in the CCE. Microbial biomass (orange circles) and normalized APA (blue bars) using chlorophyll (Chl-*a*; **a-c**), phytoplankton counts (cells mL^-1^; **d-g**), or total microorganism counts (phytoplankton + bacteria; cells mL^-1^; **h**) at stations 1-4 and SR2003. Treatments lacking a shared letter are significantly different (paired t-test, p < 0.05). Error bars represent one standard error of the mean, n = 3 (**a-g**), 2 (**h**).

APA normalized to Chl-*a* increased significantly upon +N addition at coastal stations 1-2 (N vs. Ctrl, *p* < 0.01, **Fig. 3a,b**), and upon +Fe amendments at the offshore station 4 (FeN vs. Ctrl, *p* < 0.05; Fe vs. Ctrl, *p* = 0.066, **Fig. 3c**). Cell-specific APA also showed significant stimulation by +N at station 2 (N/FeN vs. Ctrl, *p* < 0.05, **Fig. 3d,e**), and by +Fe at station 4 (Fe/FeN vs. Ctrl, *p* < 0.05, **Fig. 3f**). At the transition station 3, cell-specific APA responded positively to +Fe treatments (Fe/FeN vs. Ctrl, *p* < 0.05, **Fig. 3g**). For SR2003, APA exhibited no significant changes when normalized to phytoplankton cell abundance (**Fig. S12**), but rose significantly upon Zn, FeN, or ZnN additions when normalized to all microorganisms including heterotrophic bacteria (**Fig. 3h**).

At station 2, +Fe treatments (Fe/FeN) caused a 7-10% increase in the relative abundance of diatoms (mostly *Thalassiosira*, *Chaetoceros* and *Skeletonema*; **Fig. S13**), and a 6-8% decline in dinoflagellates (**Fig. 4a**). Upon N addition, the presence of dinoflagellates increased by 11%, while diatoms and Prymnesiophyceae diminished slightly. At station 4, Fe amendments (Fe/FeN) resulted in a larger contribution of diatoms (2-4%), dinoflagellates (4-12%), and other eukaryotes (2-3%), whereas *Pelagomonas* became 20% less abundant (**Fig. 4b**). Added N had a negligible effect on the community composition dominated by *Pelagomonas*. Overall, the transcriptional profile of the eukaryotic community was distinct between stations and showed clear differences among treatments, with larger variance between Fe-containing samples (Fe/FeN) versus N or Ctrl (**Fig. 4c**).

**Figure 4.**
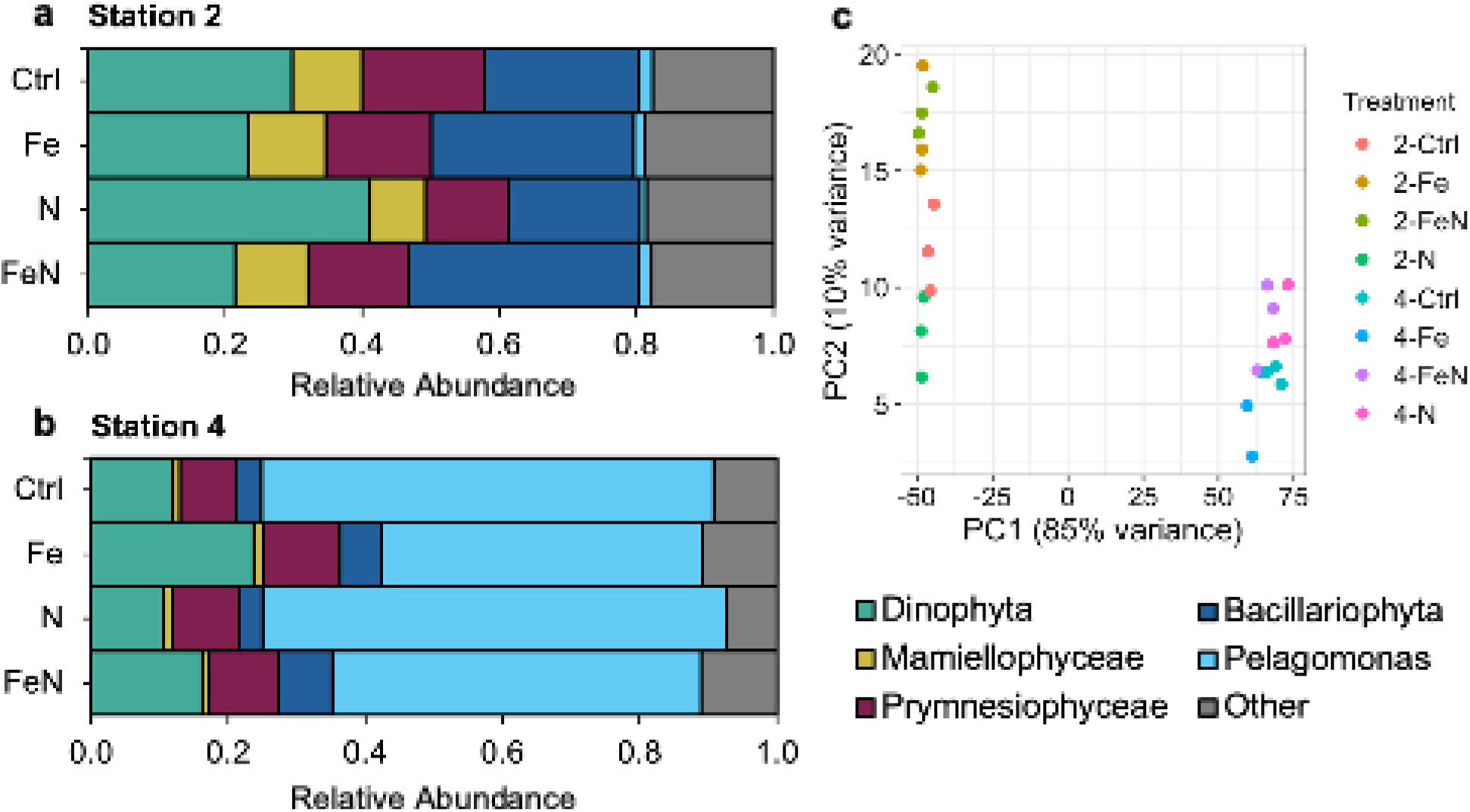
Microbial community responses to nutrient additions during incubation experiments in the CCE. a,b) Average relative abundances of major eukaryotic groups in bottle incubations at stations 2 and 4 inferred from poly(A)-selected mRNA (metatranscriptome). The average abundances were derived from the relative abundances of eukaryotic-assigned transcripts per million normalized reads (TPM). **c)** Principal Component Analysis (PCA) plots for eukaryotic mRNA within incubation experiments. Stations and treatments are denoted by color as described in the legend.

Eukaryotic community transcripts showed diminished Fe stress markers upon supplementing Fe at station 2 (Fe/FeN; **Fig. S14a**), and diminished N stress markers upon supplementing N at station 4 (N/FeN; **Fig. S14b**). In addition, amendments with one nutrient caused certain stress biomarkers for other nutrients to increase. For example, several molecular markers for P stress were elevated in +Fe incubations (*pstS, SLC34A*, *SQD1*), as well as +N incubations (*SLC34A*, *phoA*) at station 2 (**Fig. S14a**).

At the upwelling station 2, +N stimulated microbial community APA, and AP_aty_ was the most prevalent enzyme among the phosphatase-encoding genes in this treatment (34%; **Fig. S15a**). On a whole community level, the relative abundances of AP_aty_ and *phoA* genes increased in +N vs. Ctrl (**Fig. 5a**), with AP_aty_ displaying a significant fold-change (*p* < 0.001; **Fig. S16a**). On a taxonomic level, diatoms showed significant overrepresentations of AP_aty_ and *phoA* in +N vs. Ctrl (**Figs. 5c**), as well as P-stress markers *pstS* and *SLC34A* (**Figs. S17a, S18a**). Although other DOP-degrading genes were less abundant than AP_aty_ and *phoA*, several of them responded significantly to +N, including *phnY* and *ppa* in *Pelagomonas,* 5’-NT and PLC-PDE in mamiellophytes, and *ugpQ* and NUDIX in prymnesiophytes (**Fig. 5c**), yet these organisms did not display elevated P stress markers (**Fig. S17e, g, i**). Similar transcriptional responses were observed in the FeN treatment (e.g., significant increases in DOP hydrolases for all taxa in FeN vs. Ctrl; **Fig. S18**).

**Figure 5.**
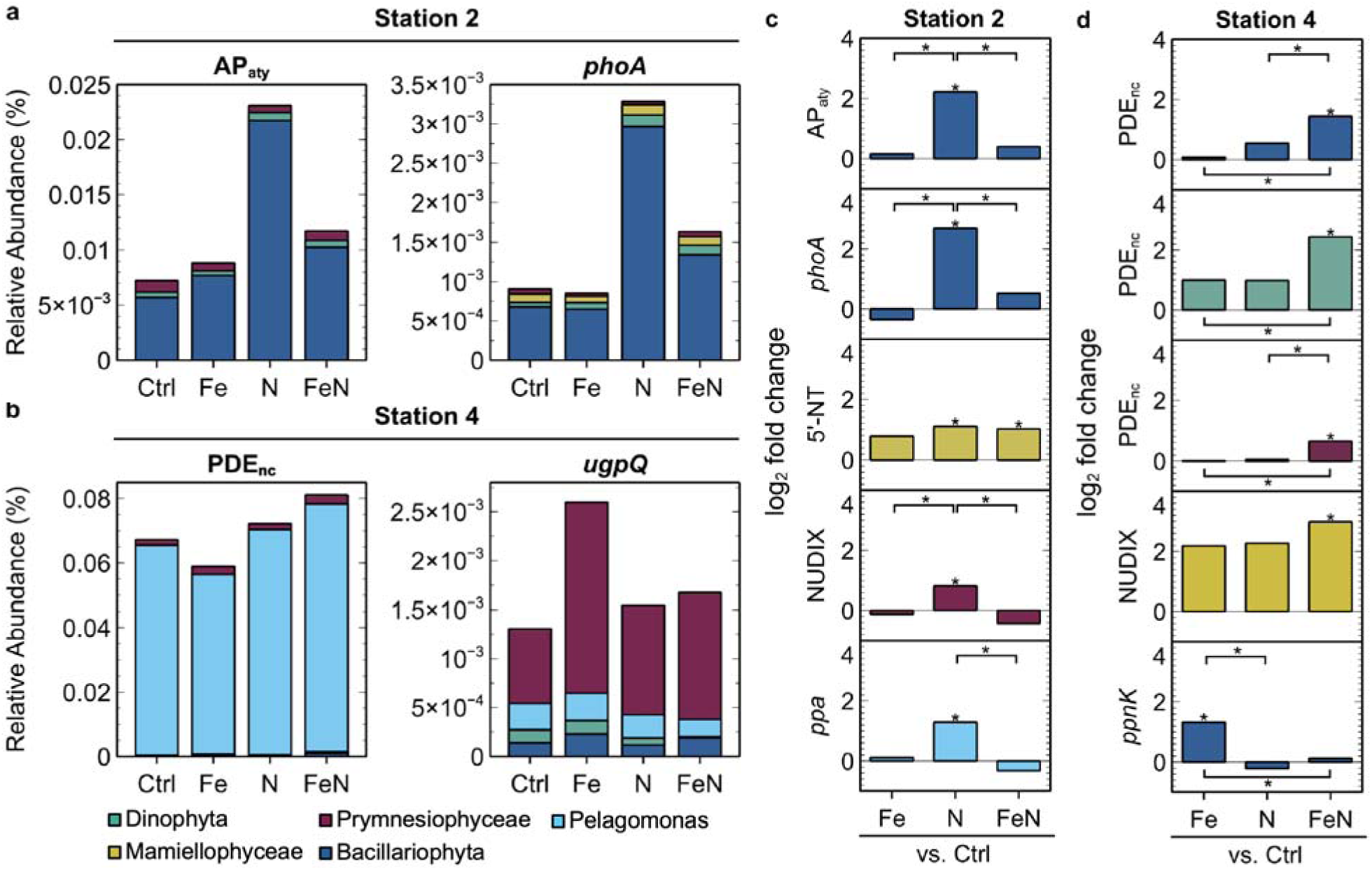
Taxon-specific transcript abundance and differential expression of phosphatases within incubation experiments. Relative abundance (%) of transcripts per million (**a, b**) and differentially expressed transcripts as log2-fold change proportion of expression (**c, d**) for major phytoplankton groups present. Fold-changes denoted with * were deemed statistically significant with an adjusted *p*-value < 0.05 (DESeq2 Wald test, Benjamini & Hochberg). Gene abbreviations are listed in Fig. 3.

At the offshore station 4, Fe/FeN additions stimulated microbial community APA, and PDE_nc_ was the dominant enzyme among the phosphatase-encoding genes for these treatments (64-72%; **Fig. S15b**). On a whole community level, the relative abundances of *ugpQ* and PDE_nc_ were elevated under +Fe and +FeN (**Fig. 5b**), respectively, compared to the control. In FeN treated samples, PDE_nc_ was significantly overrepresented in diatoms, dinoflagellates, and mamiellophytes (**Fig. 5d**), as well as NUDIX in mamiellophytes (**Fig. S18f**). In Fe treated samples, no previously considered phosphatase showed significant fold-changes within major taxa. However, Fe induced a significant spike in P stress markers within diatoms (*SLC34A*; **Fig. S17a, S18a**) alongside the NAD^+^ kinase *ppnK* (**Fig. 5d**; **Table S7**), which phosphorylates NAD under consumption of P-anhydrides.

Lastly, the prevalence of manually-curated AP_aty_ and PDE_nc_ orthologies in incubations was evaluated. PDE_nc_ expression was present and significantly regulated by nutrient availability in all major taxonomic groups (**Table S8**; **Fig. S19**). PDE_nc_ was also transcribed among the top 1% of the coding metatranscriptome pool in incubations at station 4 (**Table S9**). AP_aty_ genes were transcribed among the top 3% of all eukaryotic genes in incubations at station 2. For comparison, ISIPs were transcribed in the top 0.1-2% under Fe-limitation (station 2) and in the top 2-39% when Fe was not limiting (station 4). Both exceeded *phoA*, *phoD* and *phoX* at all stations, which only ranked in the top 14-26% expressed genes under equivalent conditions (**Table S9**), and were not always taxonomically present or environmentally regulated (**Table S8**; **Fig. S19**).

## Discussion

### Underlying P stress contributes to APA in a nutrient-rich upwelling region

The goal of this study was to examine the utilization of DOP along natural and experimental variations in nutrient availability. Typically, the activity of APs, especially PhoA, is induced in response to Pi-depletion and used as an indicator for nutritional P stress [10], a distinct metabolic state in cells experiencing low phosphate supply. The CCE was characterized as relatively P-replete by high ambient Pi concentrations of >0.7 μM, N:P ratios generally below 16, and primary limitation by either Fe or N (**Tables 1-2**). Despite this P status, we found measurable APA rates as well as eukaryotic genes for DOP utilization at all locations.

**Table 2.**
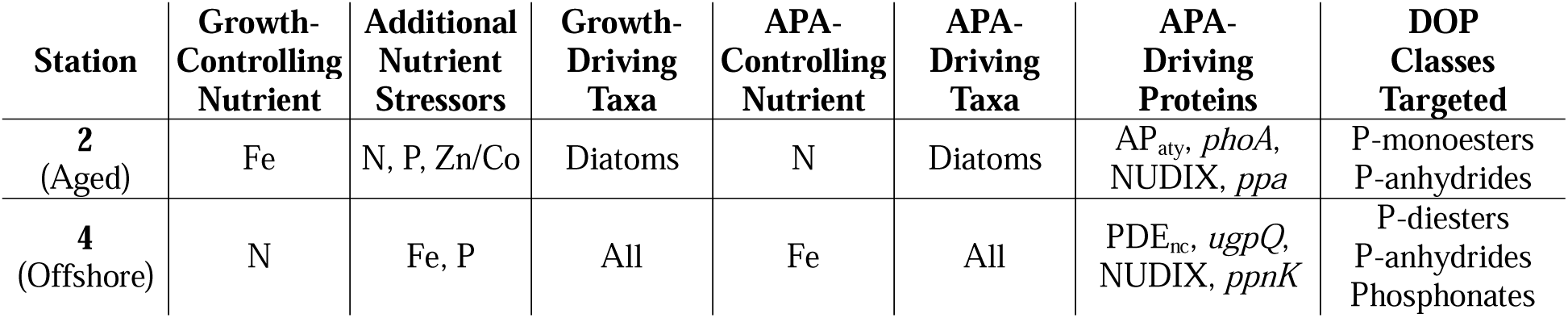
Main drivers of microbial responses within nutrient addition incubations.

APA in this study ranges within 8-38 nmol day^-1^, comparable to prior observations in the North Pacific [10,63], other eastern boundary upwelling systems [3,64], and background APA levels in Pi-replete subtropical oceans [10]. Consistent with our findings, several studies in Pacific gyres show biomass-normalized APA to be constitutive across phosphate gradients [10,12,65]. Different explanations for ‘basal’ or ‘background’ APA in high-Pi regions have been invoked, such as the regulation of PhoX by DOP rather than Pi (upregulated >90 nM DOP) [66], and constitutive or Pi-insensitive phosphatases of phytoplankton [3] or heterotrophic bacteria [67–69] that utilize DOP for carbon acquisition, i.e. PafA [68] or 5’-NTs [70].

Our results from this study provide evidence for multistressor landscapes of primary Fe limitation and additional N, P and Zn/Co stress during late upwelling (station 2), and primary N limitation with additional Fe and P stress offshore (station 4; **Table 2**). First, the low ambient N:P ratios (**Table 1**) do not necessary reflect an excess or replete P status for all taxa, since flexible microbial P stoichiometries due to metabolic plasticity can sometimes lead to non-Redfield patterns with diminished ratios in eutrophic areas and a below-Redfield global average [2]. Second, underlying P stress is supported by genetic nutrient stress indicators: Under ambient conditions, overrepresented *pstS*, *SLC34A*, and *SQD1* transcripts (**Fig. 2**), together with an above N:P ratio above 19 (**Table 1**), imply some degree of P stress for the whole eukaryotic community at station 1. In addition, all four P stress biomarkers were elevated for diatoms at station 2 (**Fig. S9**). Under incubation conditions, alleviation of proximal nutrient shortages often caused P stress genes to spike (**Fig. S14**, **S16**). For example, diatoms transcribed significantly more P stress biomarkers when either one or both Fe and N were added (**Fig. S17a, b**). Altogether, underlying P stress in one or more eukaryotic phytoplankton groups likely shapes DOP cycling in the CCE (**Table 2**), despite high Pi concentrations, as described for bloom and bust cycles [5].

### Community responses to nutrient additions and their molecular underpinnings

Metatranscriptomics were applied to elucidate molecular and taxonomic drivers of phytoplankton responses in growth and APA to Fe and N amendments. The basis of Fe-promoted APA is thought to be an increased supply of reactive cofactors for Fe-dependent DOP hydrolases [2], whereas N-promoted APA may be caused by a rise in macronutrient Redfield ratios toward P shortages [12].

#### Station 2 (aged upwelling)

At station 2, Fe-stress biomarkers, NO_3_:dFe ratios above ∼8 μmol/nmol [29], and incubations indicated proximal Fe-limitation of phytoplankton growth, as observed previously in the CCE [28–31]. This growth pattern is likely driven by chlorophyll-rich, large diatoms based on several indicators: First, community composition inferred from metatranscriptomes shows increased diatom contribution in +Fe and +FeN treated samples (**Fig. 4a**). Second, negative silica excess values (Si_ex_) present at stations 1-2 imply Fe-limitation in diatoms due to their silicic acid-containing frustules [30] (**Fig. S2**). Third, trends in Chl-*a* pigments were not mirrored using cell enumeration (**Table 1**, **Fig. 3b,e**), consistent with the removal of large chain-forming diatoms such as *Pseudo-nitzschia* (**Figs. 4a, S6b**) (see *Supplementary Material, Standard Procedures*).

At this station, microbial community APA increased in response to N additions. AP_aty_ transcripts from P-stressed diatoms likely dictated bulk APA due to their large contribution to the phosphatase pool (34%), 2.2-fold increased relative abundance, and significant fold-changes in response to +N (**Fig. 5c**), with secondary contributions from *phoA* and other proteins. This enhancement in APA was decoupled from diatom growth, presumably due to primary Fe limitation still stifling growth, analogous to reported co-limitation scenarios in the Atlantic [11]. The combination of +FeN did not lead to enhanced APA likely due to Fe-induced changes in microbial physiology and community structure (**Figs. 4a, S11**).

Both community responses in growth under +Fe/+FeN, and APA under +N were traced back to diatoms, potentially via niche partitioning into (1) growth-responsive genera which can compete successfully for inorganic P and accumulate under +Fe; and (2) APA-responsive genera which upregulate DOP acquisition mechanisms under +N but are stymied by the lack of Fe. Indeed, *Thalassiosira* and *Staurosira* expressed 49.1% and 38.4% of all diatom *phoA*, respectively, and *Pseudo-nitzschia* 92.3% of all AP_aty_ under +N (**Fig. S13**). In contrast, *Chatoceros* and *Skeletonema* contributed up to 40.9% of the diatom growth increase under +Fe/FeN, but negligibly expressed these phosphatases.

#### Station 4 (offshore zone)

At station 4, N-stress biomarkers, N:P ratios < 16, and incubations indicated that N was the primary limiting nutrient for phytoplankton growth. Metatranscriptomics suggest that N was incorporated into total biomass over the entire photosynthetic assemblage (**Fig. 4b**), which is also reflected in elevated phytoplankton cell counts (**Fig. S11**). Growth in +FeN likely differed from +N due to community composition changes associated with alleviating underlying Fe stress of several taxa (diatoms, prymnesiophytes, mamiellophytes; **Fig. S17**). Chl-*a* and cell counts aligned due to the absence of large chain-forming diatoms (**Fig. 4c,f**).

At this station, microbial community APA increased in response to Fe additions. No P-monoesterase was responsive to Fe; however, the relative abundance of PDE_nc_ - primarily from *Pelagomonas* - did increase by 12% in +FeN samples versus the control (**Fig. 5b**). Biochemical experiments demonstrate that several promiscuitive PDEs of terrestrial bacteria also target P-monoesters [71,72]. Furthermore, the metal cofactors for PDE_nc_ might involve binuclear Fe, Zn or Mn centers [73]. Hence, elevated APA upon +FeN addition can potentially be ascribed to Fe-responsive PDE_nc_ due to its environmental predominance (**Fig. 5b**) plus significant fold-changes in dinoflagellates, diatoms, and Prymnesiophyceae (**Fig. 5d**).

In +Fe samples, the relative abundance of *ugpQ* - mostly from Prymnesiophyceae - doubled compared to the control (**Fig. 5b**), making it the 2^nd^ most abundant phosphatase after PDE_nc_. In addition, significant overrepresentation of *ppnK* in diatoms implies an elevated P demand for NADP production due to mitigation of oxidative stress and metal homeostasis under sudden Fe influx [74,75], or resource allocation toward carbon fixation after replenishing Fe [76] in diatoms. This reallocation includes routing Fe toward Fe-S cluster biogenesis, also observed here (**Table S7**), and explains the significant decrease in Fe-demanding *phoD* and *phoX* (**Fig. S18**). We thus hypothesize that elevated APA under +Fe is related to *ugpQ* and *ppnK* through atypical Fe demands, unknown metabolic links, or indirect ecophysiological effects. Overall, diatoms were major drivers of community responses to nutrient fluctuations, and unclassified AP_aty_ and PDE_nc_ enzymes major drivers of APA trends (**Table 2**).

### Functional diversity among microbial taxa and implications for marine ecosystem measurements

P nutrition in different trophic states varied greatly between community members, highlighting the tailored occupation of different metabolic niches (**Fig. S19, Table S7**). Diatoms, dinoflagellates, and prymnesiophytes exhibited the widest functional diversity and transcribed all phosphatases tested (**Table S7**). However, diatom phosphatases exhibited the most significant changes in our incubation bioassays (**Fig. S18**), and dictated community responses instead of more abundant taxa (i.e. dinoflagellates). This is consistent with a study at the Oregon coast, where APs from diatoms APs mimicked *in situ* APA rates better than from other organisms [3]. This driving role is also reflected in diatoms’ normalized gene expression of AP_aty_, *phoA*, *phoD*, *phoX*, *ppa* and *ugpQ* being the highest among all taxa (**Fig. S19**). In comparison, the more streamlined green algae [77] and *Pelagomonas* [78] lacked several P-esterases and phosphonatases (**Table S7**). Notably, PDE_nc_ as well as *ppa* were most responsive overall to environmental cues based on the frequency of significant changes. Corroborating previous assumptions about harboring Fe cofactors [7,8], *phoD* (mamiellophytes), *phoX* (diatoms), and *psip1* (diatoms, dinoflagellates) reacted significantly to Fe availabilities (**Fig. S18**).

Our manually-curated gene compilations, AP_aty_ and PDE_nc_, which were not represented by canonical KOs, were more abundant, taxonomically represented, and environmentally responsive than well-studied *phoA*, *phoD* and *phoX* (**Tables S7-S8**). Their high percentile ranking among the coding metatranscriptome pool of incubation samples made PDE_nc_ and AP_aty_ comparable to highly expressed biomarkers, which can reach protein concentrations of up to 2.5 pM [56]. Given that functional gene assignments often rely on KEGG or similar databases [59], the results from AP_aty_ and PDE_nc_ illustrate that major molecular targets could be missed in studies solely relying on existing orthologies. This finding also highlights the need for more functional annotation of marine eukaryotic genes and their incorporation in routine ‘omics workflows.

Lastly, we anticipate DOP cycling in the CCE to encompass further mechanisms (see *Supplementary Material, Extended Discussion*) based on (1) additional protein classes including C-P lyases such as *phnYZ* (**Fig. S18; Table S8**), (2) additional environmental regulators including Zn or Co (**Fig. 3h, Fig. S12**), and (3) other organisms including zooplankton (**Table S9**), cyanobacteria (**Fig. S11**), and heterotrophic bacteria, which will be considered in future work.

## Conclusions

Eastern boundary upwelling systems rank among the most productive ocean biomes on Earth and provide crucial services for human society and planetary climate, yet their fate under future environmental changes remains unclear [79]. Recent studies demonstrate that marine primary productivity is hampered by multiple nutrient limitations in the global surface ocean [80]. Our study shows that even if P is not drawn down to levels that limit phytoplankton growth, several taxa in the CCE experience underlying P stress and access the DOP pool to meet nutritional P requirements. In aged, upwelled waters, Fe-limited diatoms drove increased APA upon N additions via Fe-free atypical AP_aty_ and *phoA*, since excess N exacerbated P demands. In offshore waters bordering the oligotrophic NPSG, however, the whole N-limited eukaryotic community contributed to increased APA upon Fe additions via unclassified, possibly Fe-containing PDE_nc_ by several taxa, elevated *ugpQ* abundances, and P-anhydride consuming *ppnK* in diatoms. These diverse responses highlight the occupation of different DOP-metabolizing niches by major organisms, as well as the broad functional diversity they apply toward P nutrition. Emerging evidence predicts that the Pacific subtropical gyres are shifting toward P limitation due to climate-change induced ocean stratification [81], enhanced N fixation [82], and anthropogenic N pollution [33]. Hence, we expect the mechanisms of microbial DOP utilization to become increasingly relevant in sustaining ocean fertility.

## Author contributions

V. S.: Conceptualization, Formal analysis, Funding acquisition, Investigation, Visualization, Writing – original draft. R. H. L.: Data curation, Investigation, Writing – review & editing. S. B.: Investigation. K. C. M.: Investigation. J. C. A.: Investigation. E. S.: Investigation. K. O. F.: Investigation. K. A. B.: Funding acquisition, Resources, Writing – review & editing. A. E. A.: Funding acquisition, Resources, Writing – review & editing. J. M. D.: Conceptualization, Funding acquisition, Project administration, Resources, Supervision, Writing – review & editing.

## Acknowledgements

The authors thank the captain, crew and science party of the *R/V Sally Ride* and *R/V Roger Revelle* for supporting this work. In particular, we thank Maxwell Fenton, Minerva M. Padilla, and Tyler H. Coale who assisted with sample collection. We also thank Michael R. Stukel for his contribution to the overall cruise operations as co-chief scientist, and Sydney Plummer, Monika Thukral, Stephanie A. Matthews, and Carl H. Lamborg for their assistance with shipboard experiments.

## Data availability

Dissolved inorganic nutrients for P2107 are publicly available at the Environmental Data Initiative (EDI) Data Portal [83]. RNA amplicon sequence data are deposited in the NCBI Sequence Read Archive under BioProject ID PRJNA1258289. Additional data is available from the authors upon request.

## References

1. Field CB et al. Primary production of the biosphere: Integrating terrestrial and oceanic components. Science. 1998;281(5374):237-40. doi:10.1126/science.281.5374.237.

2. Duhamel S et al. Phosphorus as an integral component of global marine biogeochemistry. Nat Geosci. 2021;14(6):359–68. doi:10.1038/s41561-021-00755-8.

3. Dyhrman ST, Ruttenberg KC. Presence and regulation of alkaline phosphatase activity in eukaryotic phytoplankton from the coastal ocean: Implications for dissolved organic phosphorus remineralization. Limnol Oceanogr. 2006;51(3):1381–90. doi:10.4319/lo.2006.51.3.1381.

4. Luo H et al. Depth distributions of alkaline phosphatase and phosphonate utilization genes in the North Pacific Subtropical Gyre. Aquat Microb Ecol. 2011;62(1):61–9. doi:10.3354/ame01458.

5. Trommer G et al. Phytoplankton phosphorus limitation in a North Atlantic coastal ecosystem not predicted by nutrient load. J Plankton Res. 2013;35(6):1207–19. doi:10.1093/plankt/fbt070.

6. O’Brie PJ, Herschlag D. Functional interrelationships in the alkaline phosphatase superfamily: Phosphodiesterase activity of *Escherichia coli* alkaline phosphatase. Biochemistry. 2001;40(19):5691–9. doi:10.1021/bi0028892.

7. Yong SC et al. A complex iron-calcium cofactor catalyzing phosphotransfer chemistry. Science. 2014;345(6201):1170-3. doi:10.1126/science.1254237.

8. Rodriguez F et al. Crystal structure of the *Bacillus subtilis* phosphodiesterase PhoD reveals an iron and calcium-containing active site. J Biol Chem. 2014;289(45):30889–99. doi:10.1074/jbc.M114.604892.

9. Wojciechowski CL, Cardia JP, Kantrowitz ER. Alkaline phosphatase from the hyperthermophilic bacterium *T. maritima* requires cobalt for activity. Protein Sci. 2002;11(4):903–11. doi:10.1110/ps.4260102.

10. Mahaffey C et al. Alkaline phosphatase activity in the subtropical ocean: Insights from nutrient, dust and trace metal addition experiments. Front Mar Sci. 2014;1(73):1–13. doi:10.3389/fmars.2014.00073.

11. Browning TJ et al. Iron limitation of microbial phosphorus acquisition in the tropical North Atlantic. Nat Commun. 2017;8(15465):1–7. doi:10.1038/ncomms15465.

12. Duhamel S, Dyhrman ST, Karl DM. Alkaline phosphatase activity and regulation in the North Pacific Subtropical Gyre. Limnol Oceanogr. 2010;55(3):1414–25. doi:10.4319/lo.2010.55.3.1414.

13. Duhamel S et al. Characterization of alkaline phosphatase activity in the North and South Pacific Subtropical Gyres: Implications for phosphorus cycling. Limnol Oceanogr. 2011;56(4):1244–54. doi:10.4319/lo.2011.56.4.1244.

14. Jickells TD et al. Global iron connections netween desert dust, ocean biogeochemistry, and climate. Science. 2005;308(5718):67-71. doi:10.1126/science.1105959.

15. Rouco M et al. Transcriptional patterns identify resource controls on the diazotroph *Trichodesmium* in the Atlantic and Pacific oceans. ISME J. 2018;12(6):1486–95. doi:10.1038/s41396-018-0087-z.

16. Young CL, Ingall ED. Marine dissolved organic phosphorus composition: Insights from samples recovered using combined electrodialysis/reverse osmosis. Aquat Geochem. 2010;16(4):563–74. doi:10.1007/s10498-009-9087-y.

17. Dyhrman ST et al. The transcriptome and proteome of the diatom *Thalassiosira pseudonana* reveal a diverse phosphorus stress response. PLoS One. 2012;7(3):e33768. doi:10.1371/journal.pone.0033768.

18. Chen X-H et al. Quantitative proteomics reveals common and specific Responses of a marine diatom *Thalassiosira pseudonana* to different macronutrient deficiencies. Front Microbiol. 2018;9(2761):2761. doi:10.3389/fmicb.2018.02761.

19. Martin P et al. Accumulation and enhanced cycling of polyphosphate by Sargasso Sea plankton in response to low phosphorus. Proc Natl Acad Sci. 2014;111(22):8089–94. doi:10.1073/pnas.1321719111.

20. Diaz JM et al. Dissolved organic phosphorus utilization by phytoplankton reveals preferential degradation of polyphosphates over phosphomonoesters. Front Mar Sci. 2018;5(380):1–17. doi:10.3389/fmars.2018.00380.

21. Diaz JM et al. Preferential utilization of inorganic polyphosphate over other bioavailable phosphorus sources by the model diatoms *Thalassiosira spp*. Environ Microbiol. 2019;21(7):2415–25. doi:10.1111/1462-2920.14630.

22. Whitney LP, Lomas MW. Phosphonate utilization by eukaryotic phytoplankton. Limnol Oceanogr Lett. 2019;4(1):18–24. doi:10.1002/lol2.10100.

23. Acker M et al. Phosphonate production by marine microbes: Exploring new sources and potential function. Proc Natl Acad Sci. 2022;119(11):e2113386119. doi:10.1073/pnas.2113386119.

24. Waggoner EM et al. Dissolved organic phosphorus bond-class utilization by *Synechococcus*. FEMS Microbiol Ecol. 2024;100(9):fiae099. doi:10.1093/femsec/fiae099.

25. Temperton B et al. Novel analysis of oceanic surface water metagenomes suggests importance of polyphosphate metabolism in oligotrophic environments. PLoS One. 2011;6(1):e16499. doi:10.1371/journal.pone.0016499.

26. Adams JC et al. Dissolved organic phosphorus utilization by the marine bacterium *Ruegeria pomeroyi* DSS-3 reveals chain length-dependent polyphosphate degradation. Environ Microbiol. 2022;24(5):2259–69. doi:10.1111/1462-2920.15877.

27. Closset I et al. Diatom response to alterations in upwelling and nutrient dynamics associated with climate forcing in the California Current System. Limnol Oceanogr. 2021;66(4):1578–93. doi:10.1002/lno.11705.

28. Hutchins DA et al. An iron limitation mosaic in the California upwelling regime. Limnol Oceanogr. 1998;43(6):1037–54. doi:10.4319/lo.1998.43.6.1037.

29. Andrew LK, Katherine B. Evidence for phytoplankton iron limitation in the southern California Current System. Mar Ecol: Prog Ser. 2007;342:91–103. doi:10.3354/meps342091.

30. Hogle SL et al. Pervasive iron limitation at subsurface chlorophyll maxima of the California Current. Proc Natl Acad Sci. 2018;115(52):13300–5. doi:10.1073/pnas.1813192115.

31. Forsch KO et al. Iron limitation and biogeochemical effects in southern California current coastal upwelling filaments. J Geophys Res: Oceans. 2023;128(11):e2023JC019961. doi:10.1029/2023JC019961.

32. Letscher RT, Moore JK. Preferential remineralization of dissolved organic phosphorus and non-Redfield DOM dynamics in the global ocean: Impacts on marine productivity, nitrogen fixation, and carbon export. Global Biogeochem Cycles. 2015;29(3):325–40. doi:10.1002/2014GB004904.

33. Kim I-N et al. Increasing anthropogenic nitrogen in the North Pacific Ocean. Science. 2014;346(6213):1102-6. doi:10.1126/science.1258396.

34. Deutsch C et al. Biogeochemical variability in the California Current System. Prog Oceanogr. 2021;196:102565. doi:10.1016/j.pocean.2021.102565.

35. Stukel MR et al. Contributions of mesozooplankton to vertical carbon export in a coastal upwelling system. Mar Ecol: Prog Ser. 2013;491:47–65. doi:10.3354/meps10453.

36. Strickland JDH, Parsons TR. A Practical Handbook of Seawater Analysis. 2 ed. vol 167. Canadian Bulletin of Fisheries and Aquatic Sciences. Fisheries Research Board of Canada; 1972:310.

37. CalCOFI Methods Manual. California Cooperative Oceanic Fisheries Investigations. Accessed Mar 06, 2025. https://calcofi.info/index.php/ccpublications/calcofi-methods

38. Hansen H, Koroleff F. Determination of Nutrients. 3 ed. Methods of Seawater Analysis. WILEY-VCH; 1999:159–228.

39. Monaghan EJ, Ruttenberg KC. Dissolved organic phosphorus in the coastal ocean: Reassessment of available methods and seasonal phosphorus profiles from the Eel River Shelf. Limnol Oceanogr. 1999;44(7):1702–14. doi:10.4319/lo.1999.44.7.1702.

40. Obata H, Karatani H, Nakayama E. Automated determination of iron in seawater by chelating resin concentration and chemiluminescence detection. Anal Chem. 1993;65(11):1524–8. doi:10.1021/ac00059a007.

41. Lohan MC, Aguilar-Islas AM, Bruland KW. Direct determination of iron in acidified (pH 1.7) seawater samples by flow injection analysis with catalytic spectrophotometric detection: Application and intercomparison. Limnol Oceanogr: Methods. 2006;4(6):164–71. doi:10.4319/lom.2006.4.164.

42. Lampe RH et al. Short-term acidification promotes diverse iron acquisition and conservation mechanisms in upwelling-associated phytoplankton. Nat Commun. 2023;14(1):7215. doi:10.1038/s41467-023-42949-1.

43. Love MI, Huber W, Anders S. Moderated estimation of fold change and dispersion for RNA-seq data with DESeq2. Genome Biol. 2014;15(12):550. doi:10.1186/s13059-014-0550-8.

44. Redfield AC. On the proportions of organic derivatives in sea water and their relation to the composition of plankton. James Johnstone Memorial Volume. University of Liverpool; 1934:176–92.

45. Dupont CL et al. Genomes and gene expression across light and productivity gradients in eastern subtropical Pacific microbial communities. ISME J. 2015;9(5):1076–92. doi:10.1038/ismej.2014.198.

46. Walworth NG et al. Why environmental biomarkers work: Transcriptome–proteome correlations and modeling of multistressor experiments in the marine bacterium *Trichodesmium*. J Proteome Res. 2022;21(1):77–89. doi:10.1021/acs.jproteome.1c00517.

47. Hockin NL et al. The response of diatom central carbon metabolism to nitrogen starvation Is different from that of green algae and higher plants. Plant Physiol. 2011;158(1):299–312. doi:10.1104/pp.111.184333.

48. Bender SJ et al. Transcriptional responses of three model diatoms to nitrate limitation of growth. Front Mar Sci. 2014;1:3. doi:10.3389/fmars.2014.00003.

49. Morrissey J et al. A novel protein, ubiquitous in marine phytoplankton, concentrates iron at the cell surface and facilitates uptake. Curr Biol. 2015;25(3):364–71. doi:10.1016/j.cub.2014.12.004.

50. McQuaid JB et al. Carbonate-sensitive phytotransferrin controls high-affinity iron uptake in diatoms. Nature. 2018;555(7697):534-7. doi:10.1038/nature25982.

51. Behnke J, LaRoche J. Iron uptake proteins in algae and the role of Iron Starvation-Induced Proteins (ISIPs). Eur J Phycol. 2020;55(3):339–60. doi:10.1080/09670262.2020.1744039.

52. Roche JL et al. Flavodoxin expression as an indicator of iron limitation in marine diatoms. J Phycol. 1995;31(4):520–30. doi:10.1111/j.1529-8817.1995.tb02545.x.

53. Graff van Creveld S et al. Divergent functions of two clades of flavodoxin in diatoms mitigate oxidative stress and iron limitation. eLife. 2023;12:e84392. doi:10.7554/eLife.84392.

54. Kellogg RM et al. Adaptive responses of marine diatoms to zinc scarcity and ecological implications. Nat Commun. 2022;13(1):1995. doi:10.1038/s41467-022-29603-y.

55. Kell RM et al. Zinc stimulation of phytoplankton in a low carbon dioxide, coastal Antarctic environment. bioRxiv. 2024:2023.11.05.565706. doi:10.1101/2023.11.05.565706.

56. Saito MA et al. Multiple nutrient stresses at intersecting Pacific Ocean biomes detected by protein biomarkers. Science. 2014;345(6201):1173-7. doi:10.1126/science.1256450.

57. Matsui H et al. Coordinated phosphate uptake by extracellular alkaline phosphatase and solute carrier transporters in marine diatoms. New Phytol. 2024;241(3):1210–21. doi:10.1111/nph.19410.

58. Wurch LL et al. Proteome changes driven by phosphorus deficiency and recovery in the brown tide-forming alga *Aureococcus anophagefferens*. PLoS One. 2011;6(12):e28949. doi:10.1371/journal.pone.0028949.

59. Cohen NR et al. Marine microeukaryote metatranscriptomics: Sample processing and bioinformatic workflow recommendations for ecological applications. Front Mar Sci. 2022;9:867007. doi:10.3389/fmars.2022.867007.

60. Lin X et al. Rapidly diverging evolution of an atypical alkaline phosphatase (PhoAaty) in marine phytoplankton: insights from dinoflagellate alkaline phosphatases. Front Microbiol. 2015;6:868. doi:10.3389/fmicb.2015.00868.

61. Li T et al. Identification and expression analysis of an atypical alkaline phosphatase in *Emiliania huxleyi*. Front Microbiol. 2018;9:2156. doi:10.3389/fmicb.2018.02156.

62. Torcello-Requena A et al. A distinct, high-affinity, alkaline phosphatase facilitates occupation of P-depleted environments by marine picocyanobacteria. Proc Natl Acad Sci. 2024;121(20):e2312892121. doi:10.1073/pnas.2312892121.

63. Sato M, Sakuraba R, Hashihama F. Phosphate monoesterase and diesterase activities in the North and South Pacific Ocean. Biogeosciences. 2013;10(11):7677–88. doi:10.5194/bg-10-7677-2013.

64. Sebastián M et al. Alkaline phosphatase activity and its relationship to inorganic phosphorus in the transition zone of the North-western African upwelling system. Prog Oceanogr. 2004;62(2):131–50. doi:10.1016/j.pocean.2004.07.007.

65. Suzumura M et al. Dissolved phosphorus pools and alkaline phosphatase activity in the euphotic zone of the Western North Pacific Ocean. Front Microbiol. 2012;3:99. doi:10.3389/fmicb.2012.00099.

66. Cerdan-Garcia E et al. Transcriptional responses of *Trichodesmium* to natural inverse gradients of Fe and P availability. ISME J. 2021;16(4):1055–64. doi:10.1038/s41396-021-01151-1.

67. Davis CE, Mahaffey C. Elevated alkaline phosphatase activity in a phosphate-replete environment: Influence of sinking particles. Limnol Oceanogr. 2017;62(6):2389–403. doi:10.1002/lno.10572.

68. Lidbury IDEA et al. A widely distributed phosphate-insensitive phosphatase presents a route for rapid organophosphorus remineralization in the biosphere. Proc Natl Acad Sci. 2022;119(5):e2118122119. doi:10.1073/pnas.2118122119.

69. Thomson B et al. Resolving the paradox: Continuous cell-free alkaline phosphatase activity despite high phosphate concentrations. Mar Chem. 2019;214:103671. doi:10.1016/j.marchem.2019.103671.

70. Ammerman JW, Azam F. Bacterial 5’-nucleotidase activity in estuarine and coastal marine waters: Characterization of enzyme activity. Limnol Oceanogr. 1991;36(7):1427–36. doi:10.4319/lo.1991.36.7.1427.

71. McLoughlin SY et al. Growth of *Escherichia coli* coexpressing phosphotriesterase and glycerophosphodiester phosphodiesterase, using paraoxon as the sole phosphorus source. Appl Environ Microbiol. 2004;70(1):404–12. doi:10.1128/AEM.70.1.404-412.2004.

72. Myers CL et al. Characterization of wall teichoic acid degradation by the bacteriophage 29 appendage protein GP12 using synthetic substrate analogs. J Biol Chem. 2015;290(31):19133–45. doi:10.1074/jbc.M115.662866.

73. Matange N, Podobnik M, Visweswariah Sandhya S. Metallophosphoesterases: structural fidelity with functional promiscuity. Biochem J. 2015;467(2):201–16. doi:10.1042/bj20150028.

74. Darby AC et al. Integrated transcriptomic and proteomic analysis of the global response of *Wolbachia* to doxycycline-induced stress. ISME J. 2013;8(4):925–37. doi:10.1038/ismej.2013.192.

75. Jamieson DJ. Oxidative stress responses of the yeast *Saccharomyces cerevisiae*. Yeast. 1998;14(16):1511–27. doi:10.1002/(SICI)1097-0061(199812)14:16<1511::AID-YEA356>3.0.CO;2-S.

76. Marchetti A et al. Comparative metatranscriptomics identifies molecular bases for the physiological responses of phytoplankton to varying iron availability. Proc Natl Acad Sci. 2012;109(6):E317–E25. doi:10.1073/pnas.1118408109.

77. Xu Y et al. Hidden genomic diversity drives niche partitioning in a cosmopolitan eukaryotic picophytoplankton. ISME J. 2024;18(1):wrae163. doi:10.1093/ismejo/wrae163.

78. Guérin N et al. Genomic adaptation of the picoeukaryote *Pelagomonas calceolata* to iron-poor oceans revealed by a chromosome-scale genome sequence. Commun Biol. 2022;5(1):983. doi:10.1038/s42003-022-03939-z.

79. Bograd SJ et al. Climate change impacts on eastern boundary upwelling systems. Annu Rev Mar Sci. 2023;15(Volume 15, 2023):303-28. doi:10.1146/annurev-marine-032122-021945.

80. Browning TJ, Moore CM. Global analysis of ocean phytoplankton nutrient limitation reveals high prevalence of co-limitation. Nat Commun. 2023;14(1):5014. doi:10.1038/s41467-023-40774-0.

81. Gerace S et al. Observed declines in upper ocean phosphate-to-nitrate availability. Proc Natl Acad Sci USA. 2025;122(6):e2411835122. doi:10.1073/pnas.2411835122.

82. Karl D et al. The role of nitrogen fixation in biogeochemical cycling in the subtropical North Pacific Ocean. Nature. 1997;388(6642):533-8. doi:10.1038/41474.

83. LTER CCE, Goericke R. Dissolved inorganic nutrients from CCE LTER process cruises, including 5 macro nutrients from water column bottle sample, 2006 - 2019 (ongoing). Accessed Mar 06, 2025. doi:10.6073/pasta/0e9975846c4bacf6deae5a3f53c6f9e1

