## Supplementary Material for "Atypical phosphatases drive dissolved organic phosphorus utilization by phosphorus-stressed phytoplankton in the California Current Ecosystem"

**for**

**Index**

**Standard Procedures	p. 02-05**

**Extended Discussion	p. 06**

**Figures S1-S19	p. 07-35**

**Tables S1-S9	p. 36-46**

**References	p. 47-48**

**Standard Procedures.**

*Chlorophyll*

Seawater (100 mL) was filtered onto 25 mm GF/F glass fiber filters with nominal pore size of 0.7 μm (Whatman), and filters were stored in the dark at -20 °C until analysis on-board the ship [1]. Samples were then extracted with 90% acetone in the dark (4 °C, 24 h) and measured using a 10AU fluorometer fitted with a red-sensitive photomultiplier tube (Turner Designs). Sample signals were calibrated using a chlorophyll-*a* standard (Sigma) and were corrected for phaeopigments by accounting for the fluorescence of extracts before and after acidification with 0.003 M HCl.

*Cell counts*

Seawater samples were preserved for flow cytometry analyses by pre-filtering through a 40 μm mesh to remove large particles per instrument requirements, fixing with 0.5% glutaraldehyde (final concentration), flash freezing in liquid nitrogen, and storing at -80 °C until analysis [2]. Microbial cell counts were conducted on a Guava EasyCyte HT flow cytometer (Millipore Sigma) with blue laser (488 nm) calibrated with instrument-specific beads. Samples were analyzed at a low flow rate (0.24 mL s^-1^) for 3 min. Phytoplankton groups were counted based on size-dependent forward scatter vs. red chlorophyll fluorescence at 675 nm (eukaryotes) or orange phycoerythrin fluorescence at 585 nm (*Synechococcus*). Size classes of eukaryotic phytoplankton were further distinguished based on forward scatter (pico-, nano- and large eukaryotes up to 40 μm). To enumerate bacteria, samples were diluted (1:100) with autoclaved filtered seawater (0.1 μm), and stained with SYBR Green I (Invitrogen) according to the manufacturer’s instructions in a 96-well plate in the dark at room temperature for 20 min. Bacterial cells were counted based on diagnostic forward scatter vs. SYBR green fluorescence at 530 nm.

*Soluble reactive phosphorus (SRP)*

Select samples for soluble reactive phosphorus (SRP) were analyzed with the following manual method [3]. Briefly, seawater was filtered through 25 mm GF/F glass fiber filters with nominal pore size of 0.7 μm (Whatman), which were previously combusted at 550 °C overnight, then acid-washed with 10% HCl overnight, and rinsed three times with Milli-Q water. The filtrate was collected in 60 mL acid-washed high density polyethylene bottles and frozen at -20 °C until analysis. SRP concentrations were measured as phosphomolybdate heteropoly acid using a standard colorimetric method [3] on a Cary 60 UV-Vis Spectrophotometer (Agilent) using a 4 cm glass spectrophotometry cell on triplicate subsamples. The detection limit, defined as three times the standard deviation of replicate blank measurements, was 13 nmol L^-1^ SRP. Data obtained by this method included station 1 (15 m), station 2 (20 m), station 3 (40 m), and station 4 (140 m; **Table 1**), and SRP depth profiles used to calculate DOP (**Fig. S3**).

*Dissolved organic phosphorus (DOP)*

Dissolved organic phosphorus (DOP) was defined as the difference between total dissolved phosphorus (TDP) and soluble reactive phosphorus (DOP = TDP-SRP). SRP values were obtained using the manual method described above. Samples for TDP were filtered through 25 mm GF/F glass fiber filters with nominal pore size of 0.7 μm (Whatman), which were previously combusted at 550 °C overnight, then acid-washed with 10% HCl overnight, and rinsed three times with Milli-Q water [4]. The filtrate was collected in 60 mL acid-washed high density polyethylene bottles and frozen at -20 °C until analysis. TDP concentrations were determined by the ash/hydrolysis method [4], in which 20 mL sample volume was treated with 0.4 mL of 0.17 M MgSO_4_ in an acid-cleaned, muffled glass jar and dried without cap at 95 °C overnight. Dried samples were ashed at 550 °C for 2 hours before cooling, acidifying with 3 mL 0.75 M HCl, and heating with cap at 80 °C for 20 min. Samples were then diluted to a total volume of 20 mL H_2_O and heated at 80 °C for another 10 min. Once at room temperature, samples were quantified using the manual method for SRP described above. Due to lack of a direct measurement, the DOP value at station 3 was modeled using Ocean Data View according to the ‘Estimating Z-Values at Arbitrary X/Y Points’ section of the ODV HowTo, Version 5, February 3 2024 [5]. A dataset of DOP concentrations at different depths along the survey track of P2107 in the CCE was used as isosurface variable for modeling (**Fig. S3b**). Desired values were estimated using the gridded field DIVA gridding procedure.

*Dissolved inorganic nutrients*

Surface concentrations of dissolved inorganic nutrients (ammonium, nitrate, nitrite, silicate, and phosphate) from the CCE-LTER process cruise P2107 were acquired following California Cooperative Oceanic Fisheries Investigations (CalCOFI) protocols [6]. Samples were dispensed into 30 mL acid-cleaned polypropylene screw-capped centrifuge tubes rinsed three times with sample before filling, and stored at -20 °C before analysis. Concentrations were measured using a colorimetric assay on a SEAL Analytical continuous segmented flow autoanalyzer 3 (AA3) at the Oceanographic Data Facility (ODF) Chemistry Laboratory, Scripps Institution of Oceanography. Data obtained by this method were ambient environmental surface concentrations (~2 m) throughout the P2107 survey transect (**Fig. 1**), and macronutrient depth profiles at stations 1-4 (**Fig. S2**). Silica excess (Si_ex_) values were calculated from inorganic nitrate and silicate data as described previously [7].

*Dissolved iron (dFe)*

Dissolved iron (dFe) was measured via chemiluminescence flow-injection analysis with hydrogen peroxide oxidation [8] (FeLume, Waterville Analytical), based on a manifold described by Lohan *et al.* [9]. Briefly, seawater was filtered using acid-cleaned 0.2 µm Acropak-200 capsule filters (VWR International) into acid-cleaned low-density polyethylene (Nalgene) bottles in a positive pressure clean van, then acidified to pH ~1.8 with HCl (Optima grade, Fisher Scientific). dFe was oxidized to Fe(III) with hydrogen peroxide (1% v/v, Optima grade), buffered in-line with an ammonium-acetate mixture, and selectively pre-concentrated on a column packed with resin (Toyopearl 650 M chelating resin) at pH ~3.5. After the sample was eluted with HCl (0.23 M, Optima grade), the production of radicals catalyzes a three-step oxidation reaction with hydrogen peroxide (0.23 M, Optima grade) and luminol (0.25 mM). dFe is eluted and oxidized with pH > 9 luminol-ammonia buffer, and the chemiluminescence (425 nm) was measured with a photomultiplier tube. Final dFe concentrations were then quantified via a Fe external standard curve (0, 0.4, 0.8, 3.2 nM prepared from a Fe in 2% HNO_3_ stock diluted in seawater) measured as described above. In-house and GEOTRACES consensus reference standards (geotraces.org) were measured in each analytical run to ensure accuracy and precision.

*Alkaline phosphatase activity (APA)*

Alkaline phosphatase activity (APA) assays [2] were carried out in a black, flat-bottom 96-well plate using 190 μL sample volume and 10 μL of the fluorogenic probe 4-methylumbelliferyl phosphate (MUF-P, Millipore Sigma). Reactions were initiated by addition of MUF-P (400 μM stock solution in H_2_O; 20 μM final concentration), mixed, and incubated at room temperature. Degradation of MUF-P to 4-methylumbelliferone (MUF) with concomitant release of PO_4_^3-^ was analyzed by a standard fluorescence technique (*5*) (excitation: 359 nm, emission: 449 nm) every 5-10 min for 3-24 h using a SpectraMax M3 multimode microplate reader (Molecular Devices). For control reactions, trace metal clean seawater from each station was filtered in-line with an acid-cleaned 0.2 µm Acropak-200 filter (VWR international) and stored at -20 °C prior to analysis. To determine background MUF-P autohydrolysis, filtered seawater samples were thawed, boiled (99 °C, 20 min), amended with nutrients (see *Field incubations*), and reacted with 20 μM MUF-P. Autohydrolysis rates were subtracted from the experimental APA rates. Data were also corrected for background fluorescence present in unamended controls and calibrated with a multipoint standard curve of MUF (10–500 nmol L^-1^) prepared in 0.2 μm-filtered seawater. MUF-P hydrolysis rates were calculated as the slope of the best fit line (R^2^ ≥ 0.98) of MUF concentration versus time. A final concentration of 20 μM MUF-P was assumed to be rate-saturating based on preliminary experiments with natural samples. Thus, the rates of APA reported herein represent maximum extracellular hydrolysis rates, consistent with previous investigations of APA [10].

*Metatranscriptomics*

Samples for eukaryotic mRNA [11] were collected by filtering 3 L or 1 L of seawater from Niskin or incubation bottles, respectively, through a 0.22 μm Sterivex-GP filter (Millipore). The filter was immediately sealed with both a luer-lock plug, previously cleaned with 1.2 mol L^−1^ HCl then Milli-Q, and Hemato-Seal^TM^ tube sealant, wrapped in aluminum foil, and flash frozen in liquid nitrogen. RNA was extracted using the NucleoMag RNA extraction kit (Machery-Nagel) on an Eppendorf epMotion 5075t, as described previously [12]. RNA quality was assessed using an Agilent 2200 TapeStation (RNA ScreenTape), and quantified using the Quant-iT^TM^ Ribogreen® RNA assay kit. Poly-A selected library construction and sequencing was performed at the UC Davis Genome Center, using an Illumina NovaSeq 6000. Raw reads were quality trimmed with trimmomatic v0.39 [13] and evaluated with FastQC v0.12.1 and MultiQC v1.9.dev0 [14]. Ribosomal RNA reads were also removed with SortMeRNA v4.3.6. The remaining reads were then assembled into contigs using both individual samples with rnaSPAdes v3.13.0 [15] and replicate samples with MEGAHIT v1.2.9 using the meta-large preset [16]. Assemblies were combined and deduplicated with dedupe.sh from BBMap v38.18. Proteins were predicted with GenemarkS-T [17], which were then clustered based on 98% similarity with MMSeqs2 v13-45111 and filtered for rRNA with SortMeRNA again. Reads were mapped to the predicted proteins with Bowtie2 v2.5.1 in local alignment mode [18]. Only reads with a MAPQ score ≥ 8 were retained. Taxonomic annotation of predicted proteins was performed with DIAMOND (v2.1.8) BLASTP searches against PhyloDB v1.076 [19,20] with an E-value cutoff of 10^-5^, minimum query coverage of 60%, and alignment sores in the top 95%. To avoid mis-annotations from contamination in the reference database, final taxonomic assignment was based on the Lineage Probability Index from the BLASTP hits with aforementioned cutoffs [19,21]. For functional annotation, DIAMOND (v2.1.8) BLASTP searches were performed against the Kyoto Encyclopedia of Genes and Genomes (KEGG; Release 94.1) [22]. KEGG Ortholog (KO) assignment was performed with KOfamKOALA which utilizes hmmsearch against KOfam, an HMM database of KOs [19,23]. Genes without KOs were manually annotated based on their KEGG gene annotation as listed in **Tables S1-S5**. Relative abundances (%) of eukaryotic transcripts were assessed by dividing the total number of transcripts per million (TPM) for each gene of interest by the total number of eukaryotic TPM. Z-score normalization was applied to standardize relative abundances to a mean of 0 and standard deviation of 1, by subtracting the dataset mean from each datapoint, then dividing by the dataset standard deviation.

**Extended Discussion.**

We anticipate DOP cycling in the CCE to encompass further mechanisms based on (1) additional protein classes, (2) additional environmental regulators, and (3) other organisms, detailed below.

(1) The MUF-P assay applied here was designed for P-monoesterases and does not target other protein classes, for which field bioassays are limited [24]. For instance, phosphonate-degrading *phnY* was significantly overrepresented in *Pelagomonas* for both +N and +FeN treatments at station 2 (**Fig. S16**), indicating that eukaryotic C-P lyase expression might also play a role in P nutrition, as known for bacterioplankton [25,26].
(2) Zn/Co levels have the potential to regulate DOP utilization, as Zn-limitation of APA has been observed in the field [10]. Indeed, biomarkers for Zn/Co stress were elevated at station 2 (**Fig. 2**), and phytoplankton growth proximally limited by Zn in SR2003 (**Fig. S11**). Here, Zn additions significantly stimulated APA when normalized to both eukaryotic and prokaryotic cell counts (**Fig. 3h**), while Co additions had no effect (**Fig. S11**).
(3) Activity exhibited cyanobacteria [27], heterotrophic bacteria [28-30], and zooplankton [31] can contribute to APA observations. *PhoA* sequences in particular originated from non-photosynthetic eukaryotes (87-98% “other”, **Fig. S10**) including zooplankton and rhizaria (**Table S9**). Our incubation experiments lacked organic carbon amendments and were designed to primarily stimulate phytoplankton rather than heterotrophs. Indeed, no trends were found when using cell enumerations of heterotrophic bacteria (data not shown). Yet, we anticipate that cyanobacteria, which were present at stations 1-3 (**Fig. S17**), and heterotrophic bacteria contribute substantially to DOP cycling in the CCE, which should be considered in future work.

**Fig. S1.**

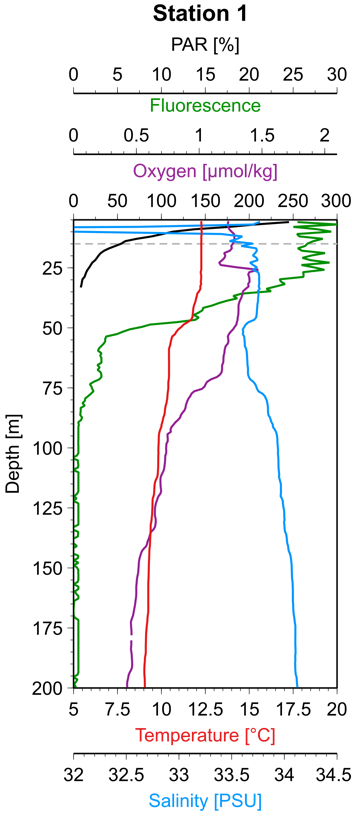

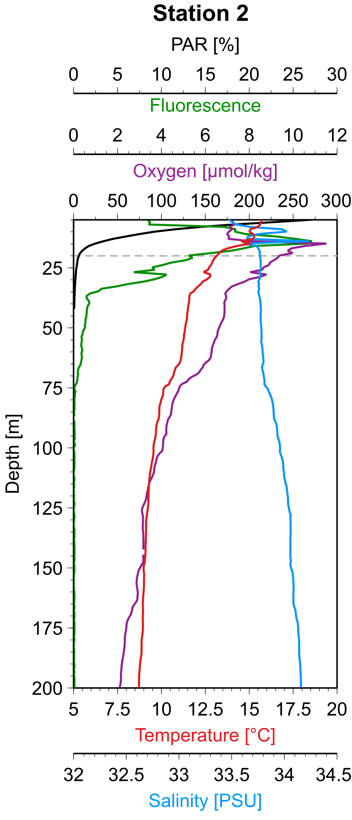

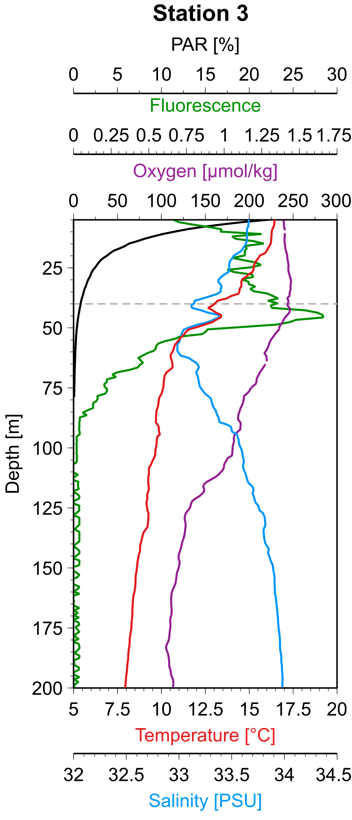

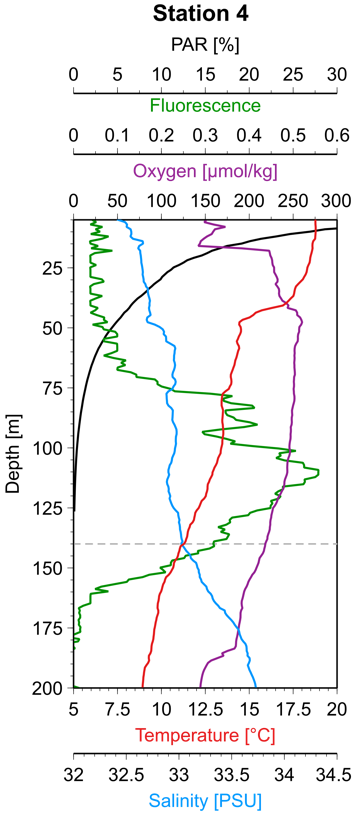

**Biogeochemical depth profiles at stations 1-4 in the California Current Ecosystem**. Water column depth profiles of photosynthetic active radiation/PAR (%; black), fluorescence (Volts; green), oxygen (mmol/kg; purple), temperature (°C; red) and salinity (PSU; blue) from the CTD (conductivity, temperature, depth) downcast prior to the start of bioassay experiments, with incubation source water depth denoted by a gray horizontal dashed line.

**Fig. S2.**

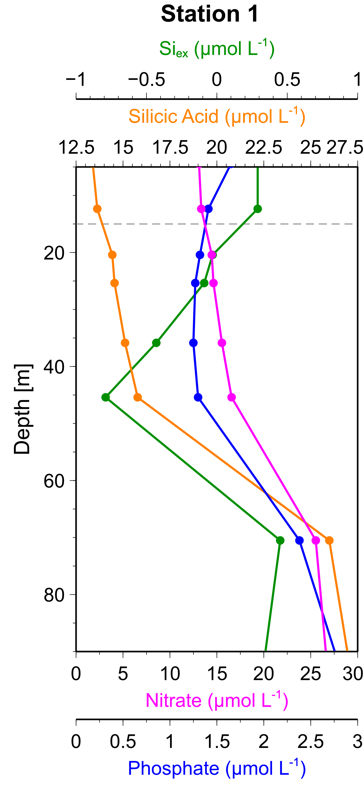

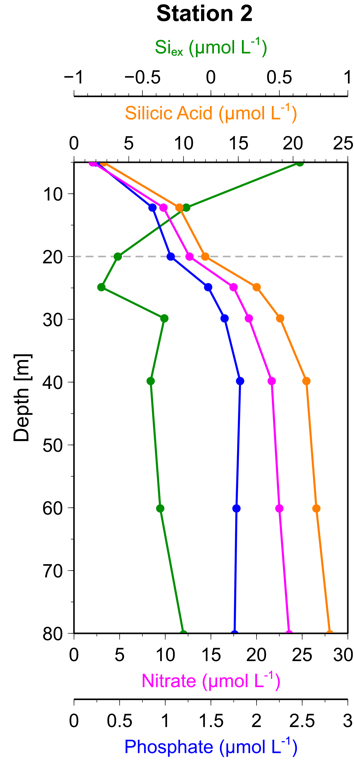

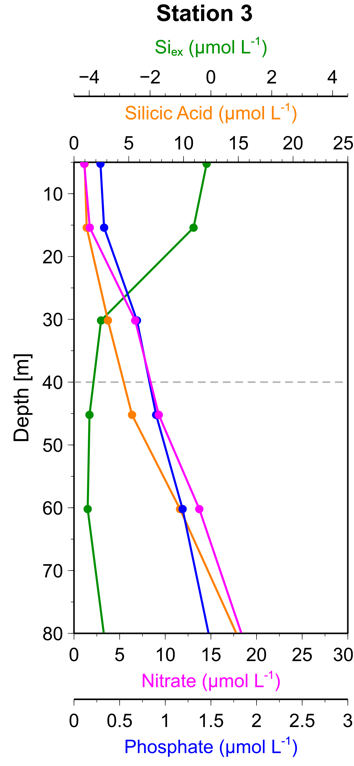

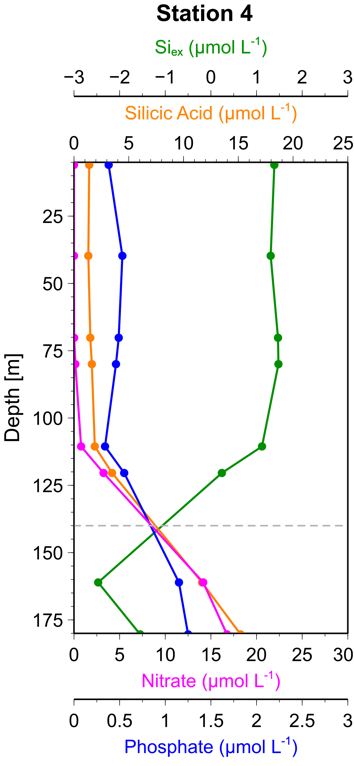

**Macronutrient depth profiles at stations 1-4 in the California Current Ecosystem**. Water column depth profiles of silica excess/Si_ex_ (μmol L^-1^; green), silicic acid (μmol L^-1^; orange), nitrate (μmol L^-1^; magenta) and phosphate (μmol L^-1^; blue) from the CTD (conductivity, temperature, depth) downcast prior to the start of bioassay experiments, with incubation source water depth denoted by a gray horizontal bar. Negative Si_ex_ values indicate iron-limitation of diatoms [7]. Note that initial phosphate concentrations within incubation bottles were determined via a different method (see *Standard Procedures*), and values are listed in the main manuscript (**Table 1**).

**Fig. S3.**

**A**

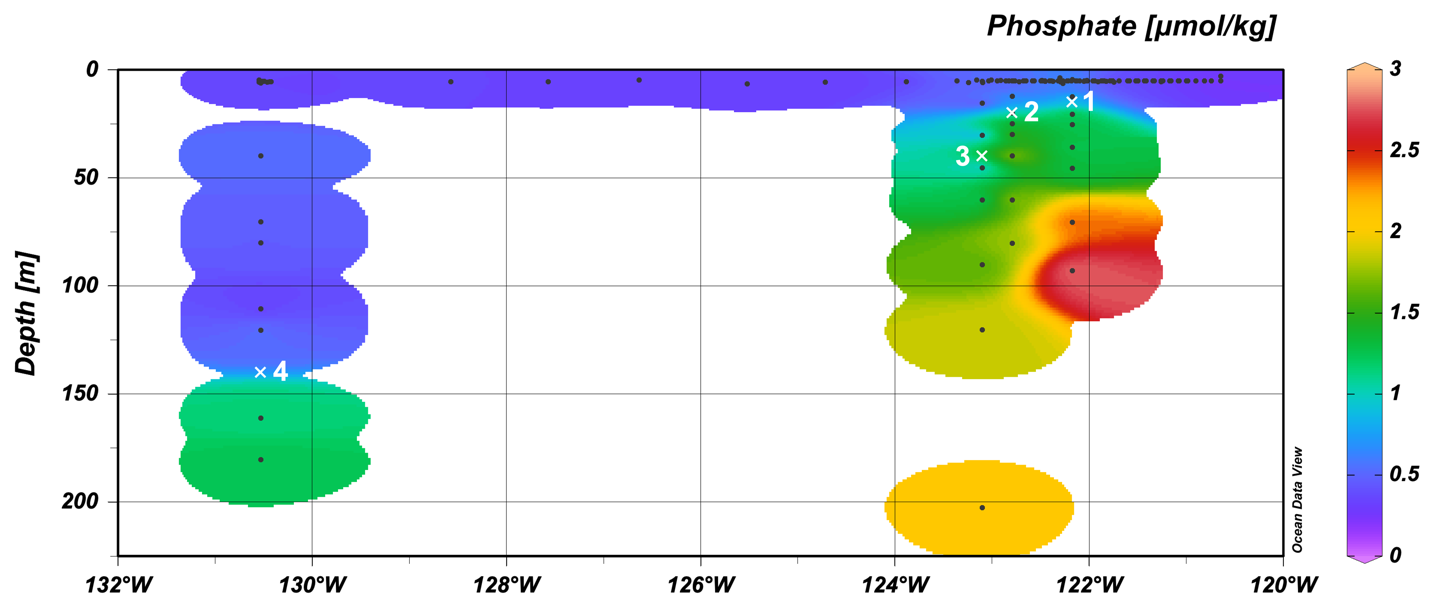

**B**

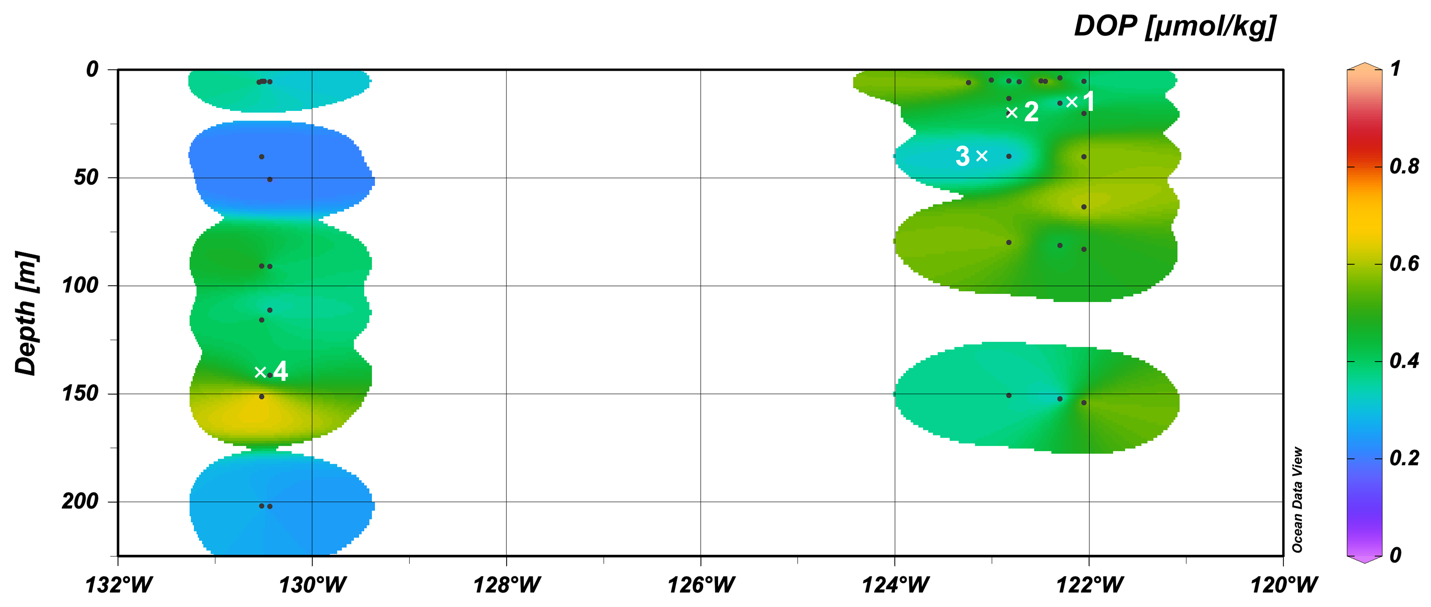

**Phosphorus concentrations at surface and with depth in the California Current Ecosystem**. Water column depth profiles of inorganic phosphate (PO_4_^3-^; **A**), and dissolved organic phosphate (DOP; **B**) from the CTD (conductivity, temperature, depth) downcast prior to the start of bioassay experiments, displayed as weighted-average gridded field. Stations 1-4 are denoted by white marks at the depths where water was collected for on-deck incubations.

**Fig. S4.**

**A	B**

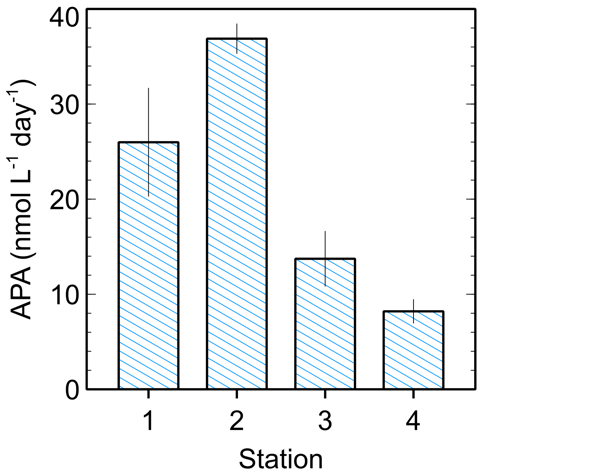

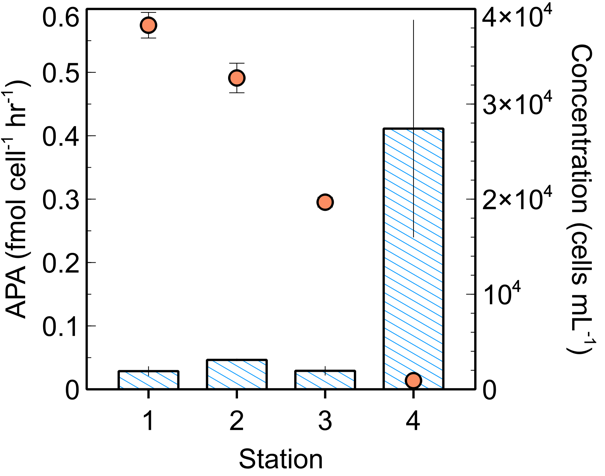

**Ambient alkaline phosphatase activity at stations 1-4 in the CCE.** Blue bars indicate *in situ* volumetric APA **(A)** or specific APA normalized to phytoplankton cell concentration **(B)**, and orange circles indicate cell concentration (cells mL^-1^) at stations 1 (Upwelling, 15 m), 2 (Aged, 20 m), 3 (Transition, 40 m) and 4 (Offshore, 140 m). Error bars represent one standard error of the mean, *n* = 3.

**Fig. S5.**

**A	B**

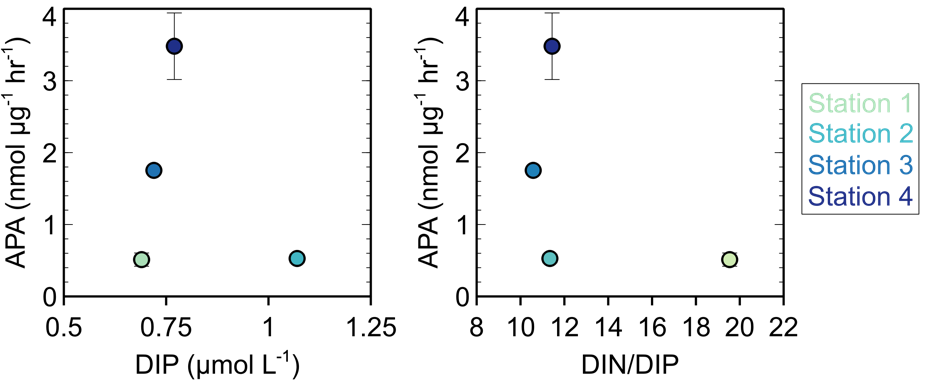

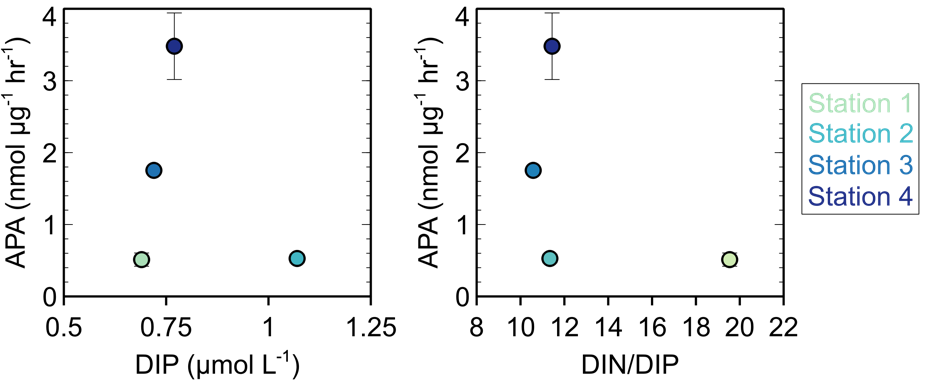

**Alkaline phosphatase activity versus DIP (A) and DIN/DIP ratios (B) at stations 1-4 in the California Current Ecosystem.** Indicated are stations 1 (Upwelling, 15 m), 2 (Aged, 20 m), 3 (Transition, 40 m) and 4 (Offshore, 140 m). DIP = dissolved inorganic phosphate, DIN = dissolved inorganic nitrate (nitrate + nitrite + ammonia).

**Fig. S6.**

**A**

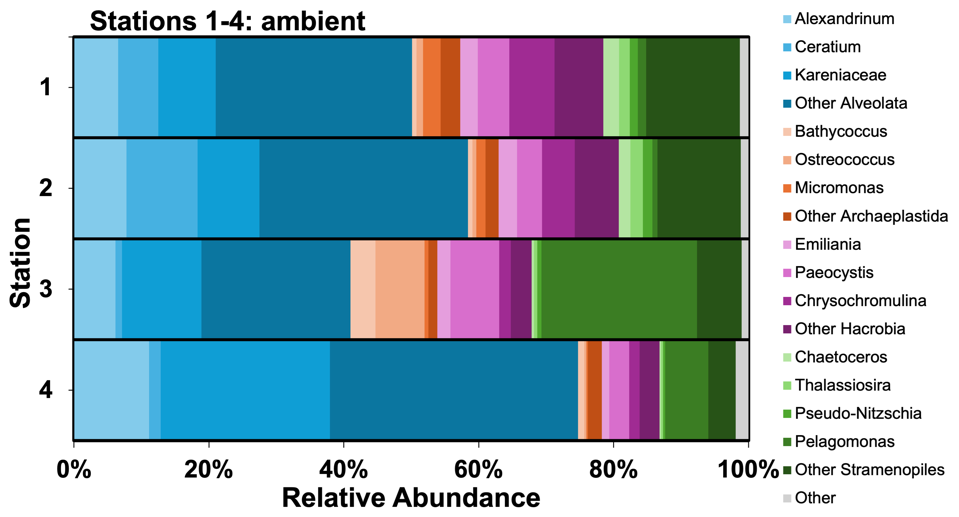

**B**

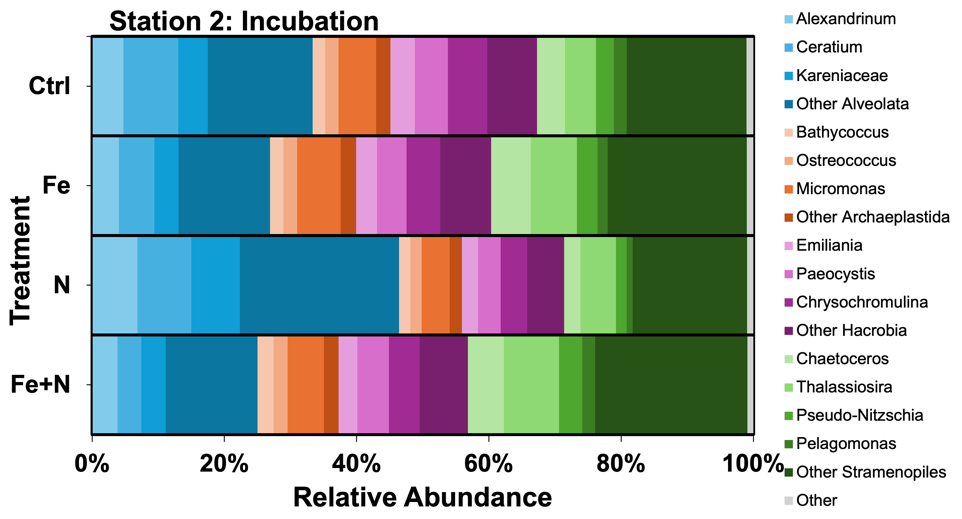

**C**

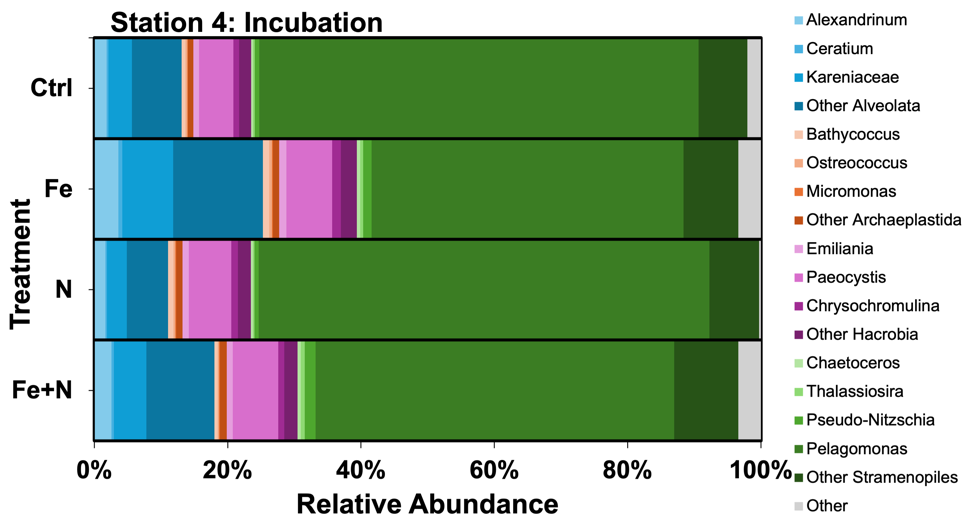

**Microbial community composition under ambient and incubation conditions in the CCE.** The average relative abundances of major photosynthetic eukaryotic taxa under ambient (**A**) and experimental conditions (**B**,**C**) from poly(A)-selected mRNA (metatranscriptome). The average abundances were derived from the relative abundances of eukaryotic-assigned transcripts per million normalized reads (TPM). Samples were from stations 1 (Upwelling, 15 m), 2 (Aged, 20 m), 3 (Transition, 40 m) and 4 (Offshore, 140 m).

**Fig. S7.**

**A**

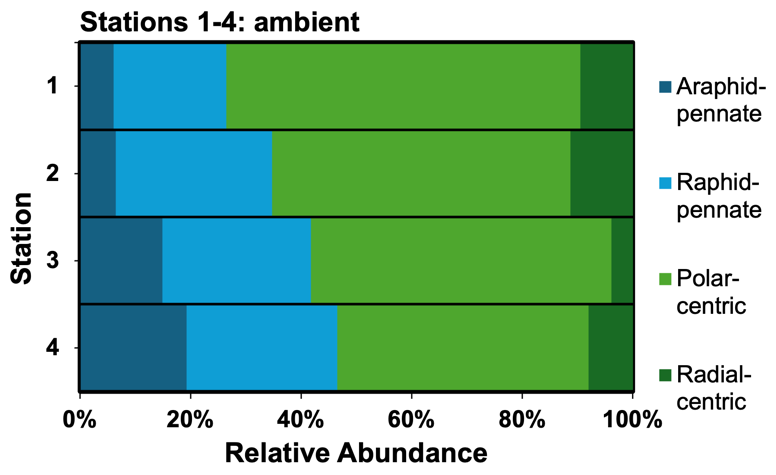

**B**

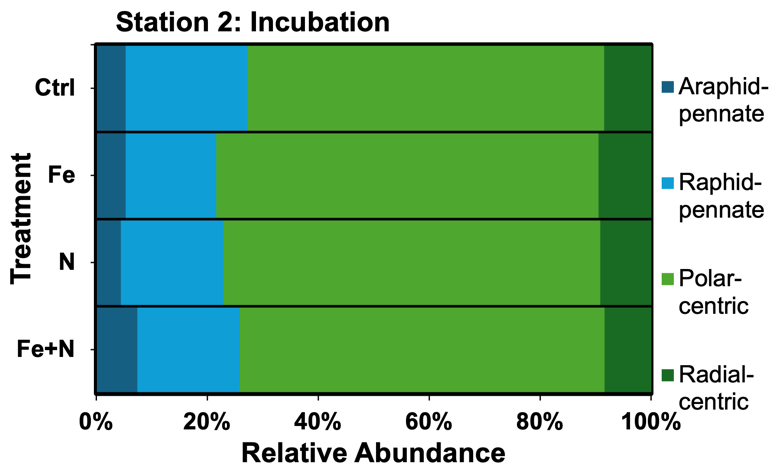

**C**

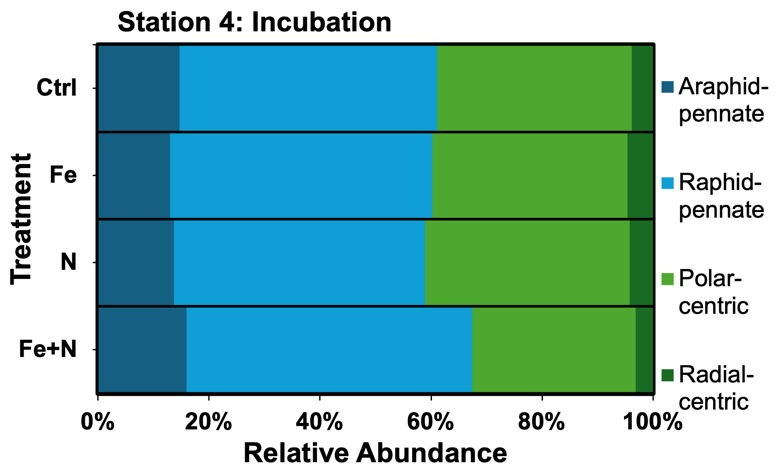

**Community structure of diatoms under ambient and incubation conditions in the CCE.** The average relative abundances of centric and pennate diatoms under ambient (**A**) and experimental conditions (**B**,**C**) from poly(A)-selected mRNA (metatranscriptome). The average abundances were derived from the relative abundances of eukaryotic-assigned transcripts per million normalized reads (TPM). Samples were from stations 1 (Upwelling, 15 m), 2 (Aged, 20 m), 3 (Transition, 40 m) and 4 (Offshore, 140 m).

**Fig. S8.**

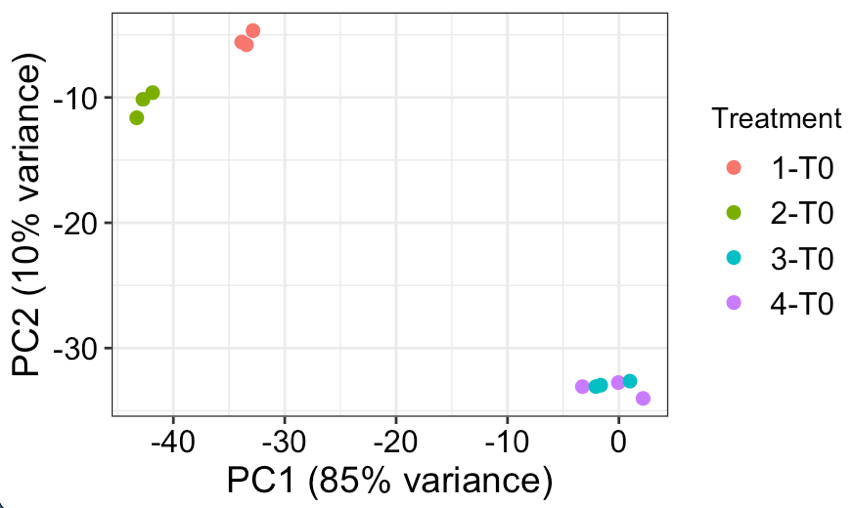

**Principal Component Analysis (PCA) for eukaryotic mRNA** **at stations 1-4 in the CCE**. Stations 1 (Upwelling, 15 m), 2 (Aged, 20 m), 3 (Transition, 40 m) and 4 (Offshore, 140 m) are denoted by color as described in the legend.

**Fig. S9.**

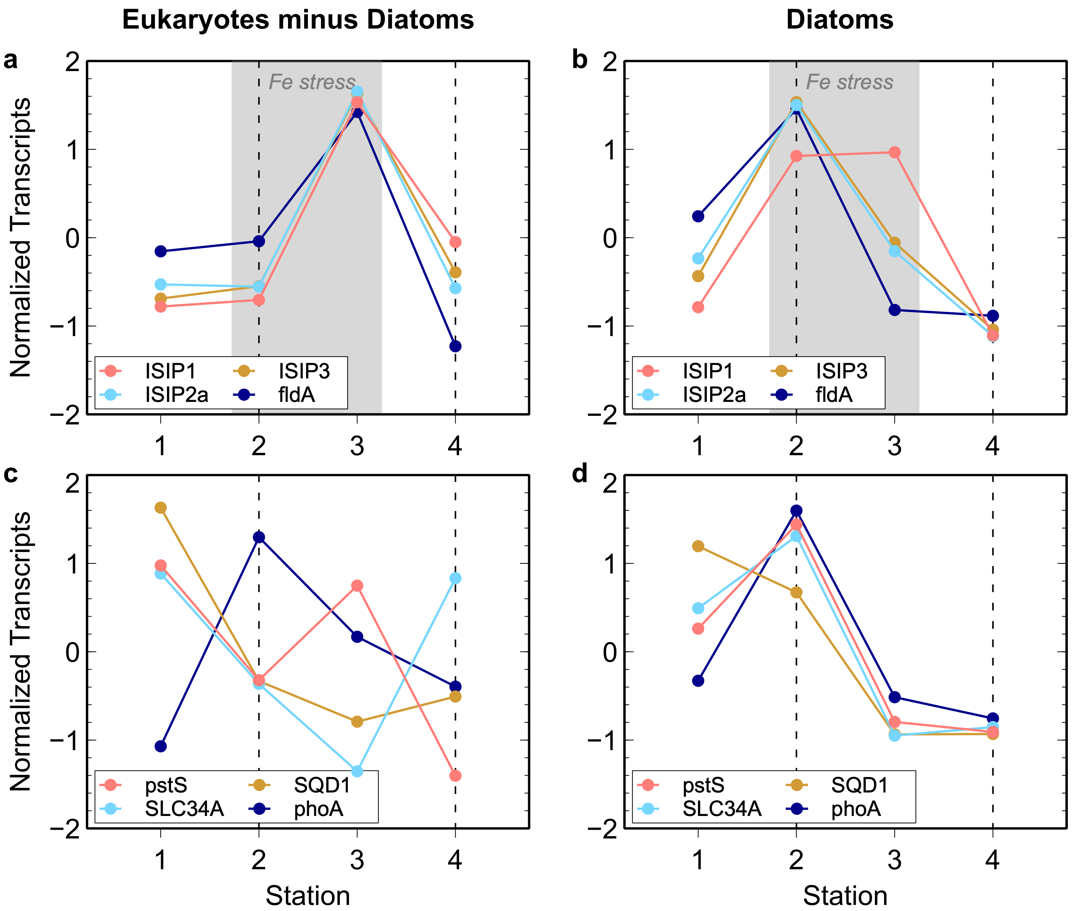

***In situ* taxon-dependent transcriptomic signatures of phytoplankton community nutrient stress at stations 1-4.** Established stress biomarkers for iron (Fe) and phosphorus (P) as z-score normalized relative abundances (%) of transcripts per million within the eukaryotic taxa except diatoms (**a, c**) and diatoms only (**b, d**). Dotted lines indicate stations selected for bioassays; shaded areas denote assigned nutrient stress for the whole eukaryotic community. ISIP1-3, iron stress responsive proteins 1-3; *fldA*, flavodoxin A; *pstS*, high-affinity phosphate transport system substrate-binding protein; SLC34A, sodium-dependent phosphate cotransporter; SQD1, UDP-sulfoquinovose synthase; *phoA*, alkaline phosphatase PhoA.

**Fig. S10.**

**A**

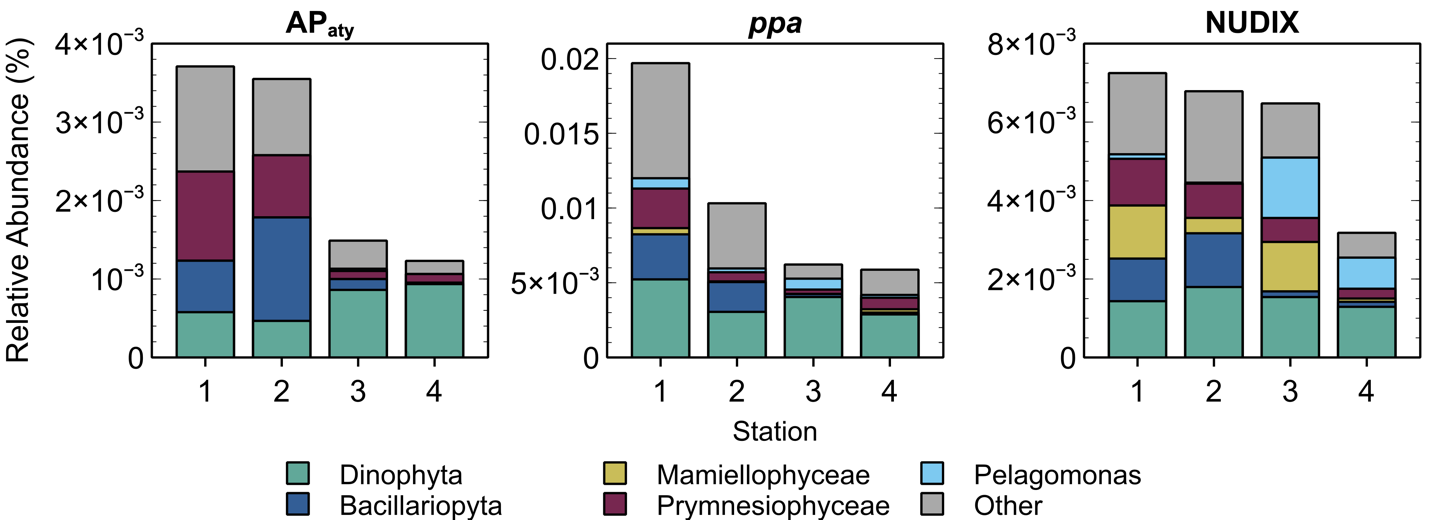

**B**

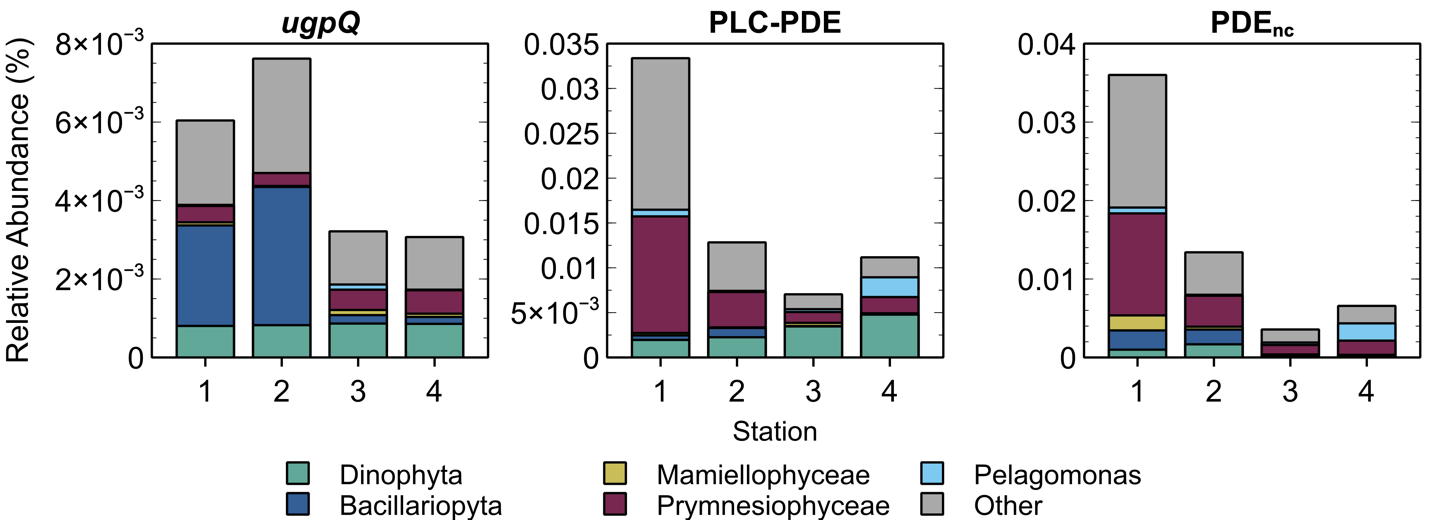

**C**

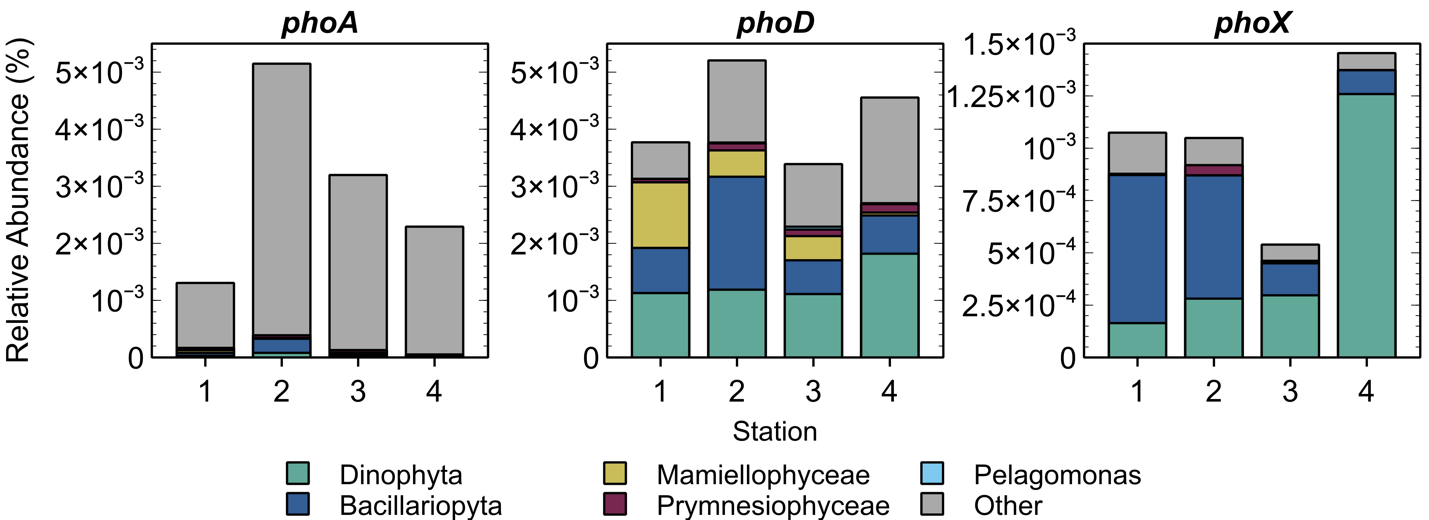

**Eukaryotic taxonomic assignment of DOP-degrading enzymes under ambient conditions.** Most abundant DOP hydrolases (**A-B**) and main AP isoforms (**C**) at stations 1-4 among all eukaryotic taxa indicated as relative abundance (%) of transcripts per million (TPM). AP_aty_, atypical alkaline phosphatase; ppa, inorganic pyrophosphatase; NUDIX, (di)nucleoside polyphosphate hydrolase; PLC-PDE, PLC-like phosphodiesterase; PDE_nc_, unclassified phosphodiesterase; *phoA*, alkaline phosphatase PhoA; *phoD*, alkaline phosphatase PhoD; *phoX*, alkaline phosphatase PhoX.

**Fig. S11.**

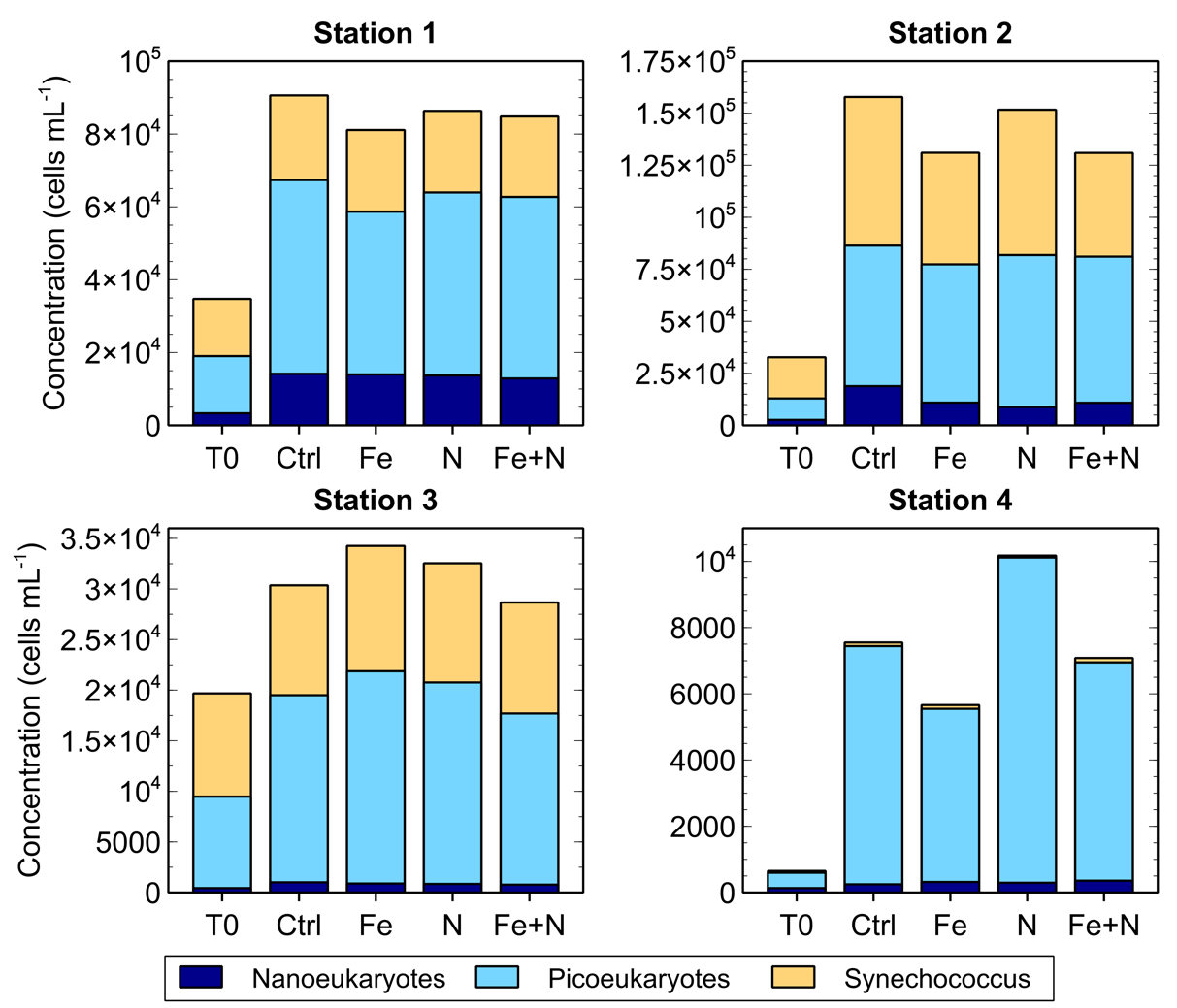

**Cell abundances of major phytoplankton size classes in field incubation bioassay experiments.** Nanoeukaryotes (dark blue), picoeukaryotes (light blue), and *Synechococcus* (yellow) abundances (cells mL^-1^) as determined by flow cytometry analysis. Large organisms (> 40 μm), e.g. diatom chains, were filtered out due to instrument requirements.

**Fig. S12.**

**A**

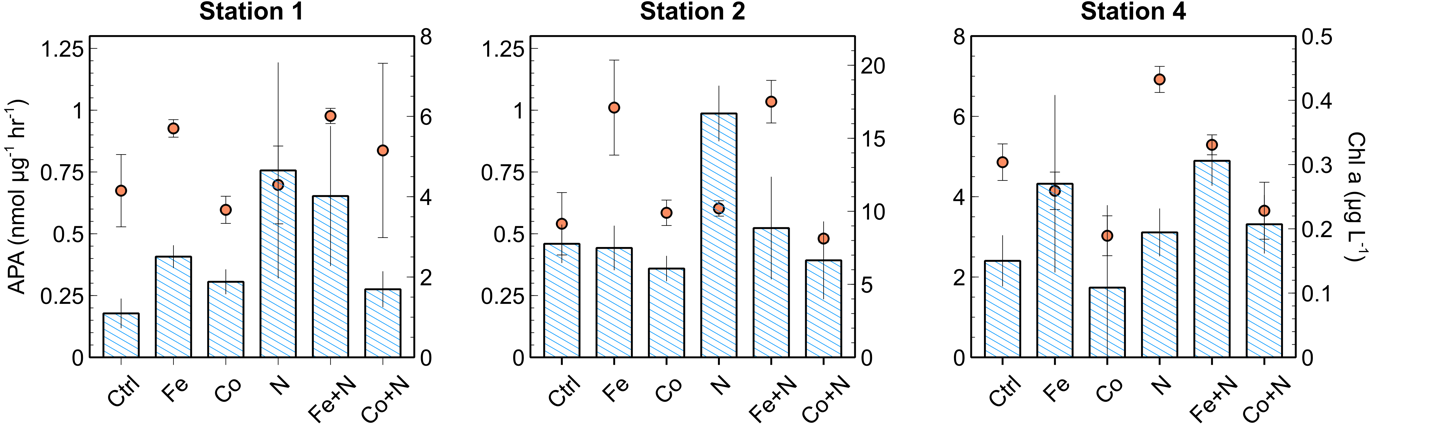

**B**

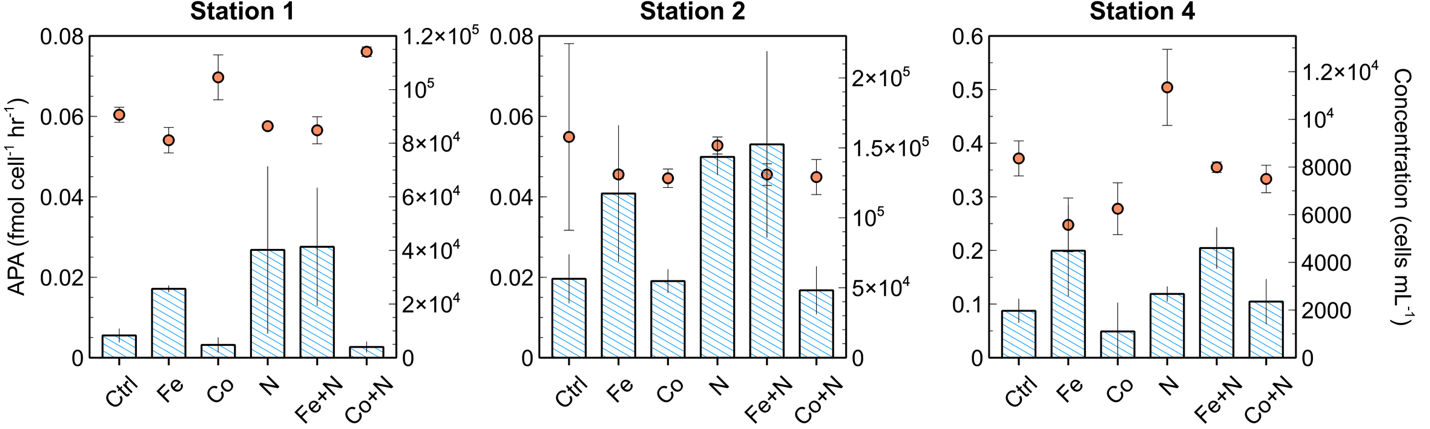

**C**

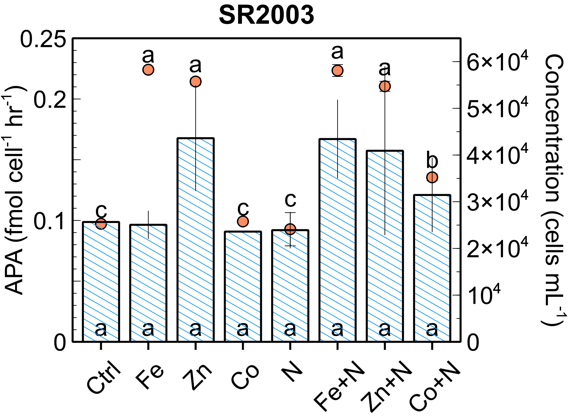

**Microbial growth and alkaline phosphatase activity (APA) at experiment locations in the CCE including Co treatments.** Microbial biomass (orange circles) and normalized APA (blue bars) using chlorophyll-a (Chl-*a*; **A**) or phytoplankton cell concentration (cells mL^-1^; **B**) at experimental stations 1 (Upwelling), 2 (Aged), 4 (Offshore), and SR2003. Treatments lacking a shared letter are significantly different (paired t-test, p < 0.05). Error bars represent one standard error of the mean, n = 3 (**A-B**), 2 (**C**).

**Fig. S13.**

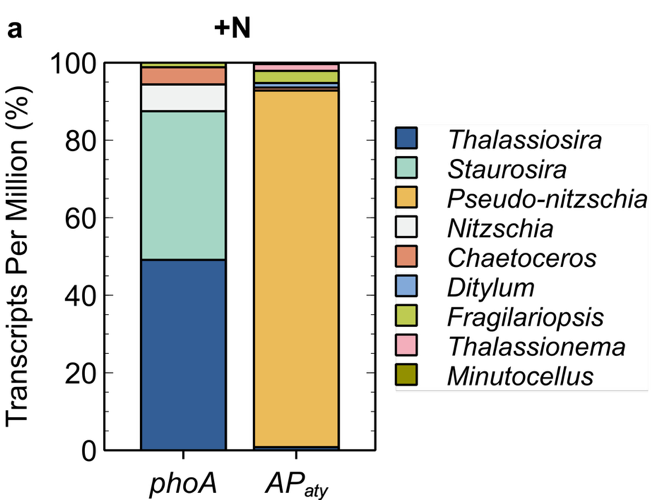

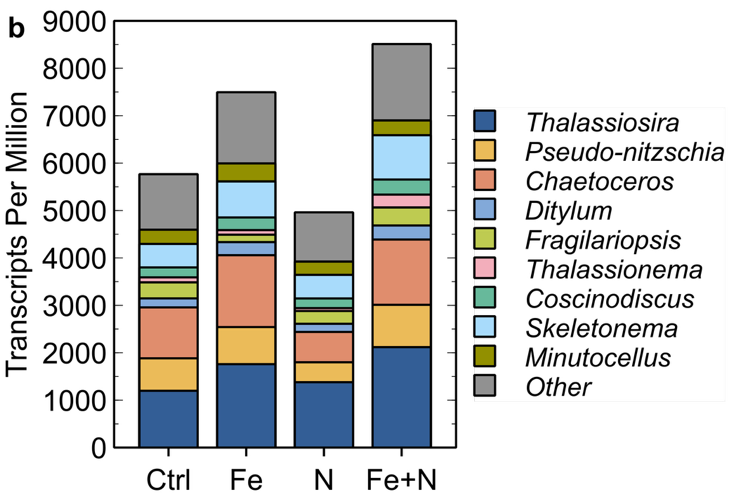

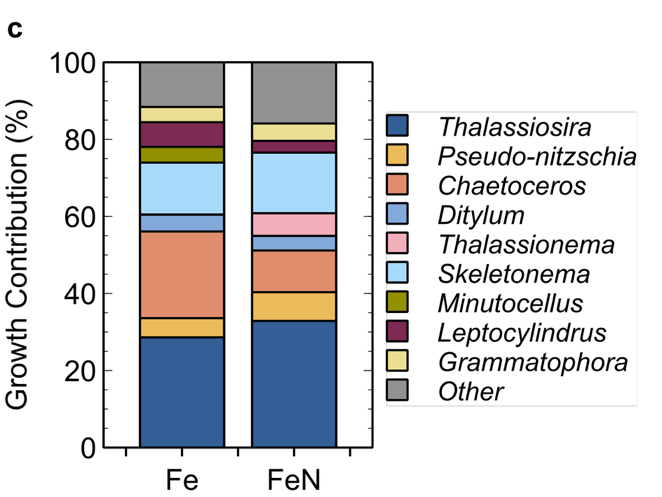

**Diatom species-specific phosphatase genes *phoA* and AP_aty_, community composition, and growth contribution in incubations at station 2.** Abundance is indicated as percent (**a**) or absolute (**b**) transcripts per million (TPM). Growth contribution (%) of Fe vs. Ctrl and FeN vs. Ctrl was calculated as the growth increase of a certain species divided by the growth increase of all diatoms in TPM, normalized so that the contribution of declining (negative) species equals 0 (**c**). *phoA*, alkaline phosphatase PhoA; AP_aty_, atypical alkaline phosphatase.

**Fig. S14.**

**A	Station 2 (Aged)**

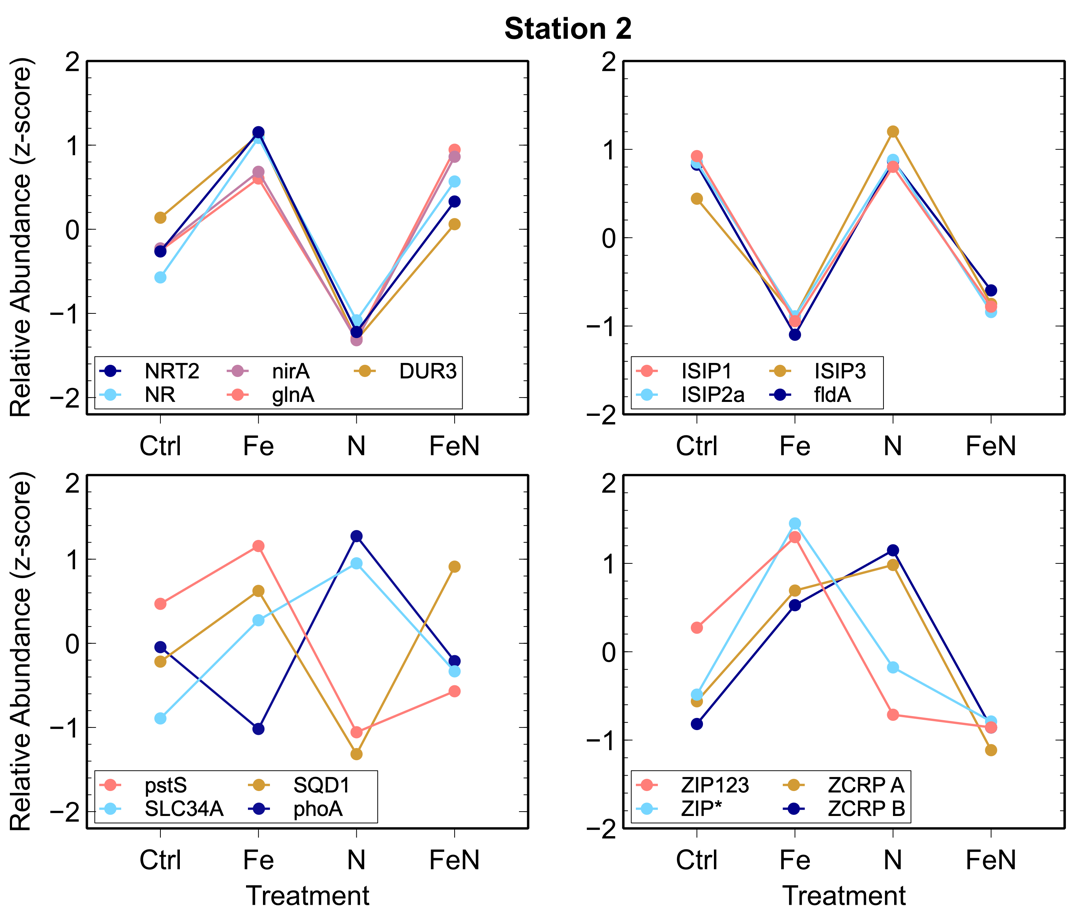

**B	Station 4 (Offshore)**

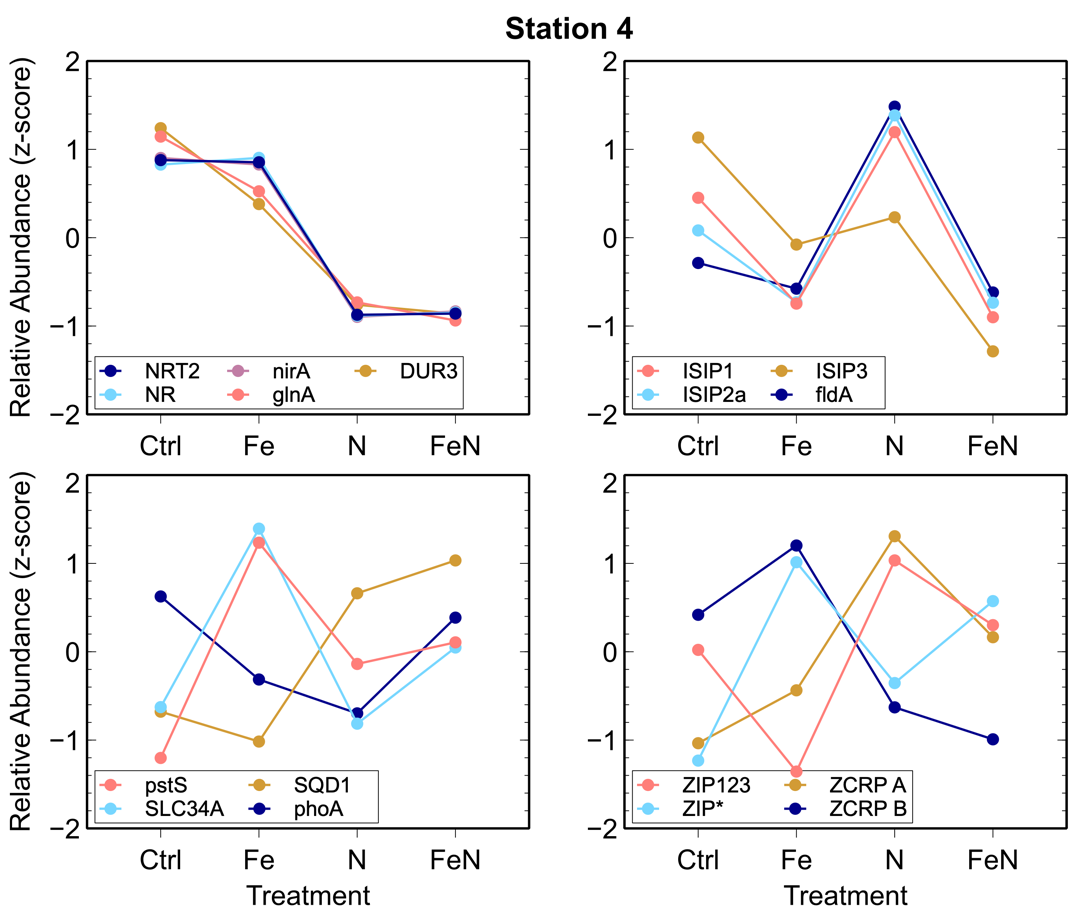

**Transcriptomic signatures of phytoplankton community nutrient stress during incubation bioassays.** Established stress biomarkers for nitrogen (N), iron (Fe), phosphorus (P) and zinc/cobalt (Zn/Co) as z-score normalized relative abundances (%) of transcripts per million after incubation bioassays at stations 2 (coastal; **A**) and 4 (offshore; **B**). NRT2, nitrate/nitrite transporter; NR, NAD(P)H-nitrate reductase; *nirA*, ferredoxin-nitrite reductase; *glnA*, glutamine synthetase; DUR3, urea-proton symporter; ISIP1-3, iron stress responsive proteins 1-3; *psbC*, photosystem II CP43 chlorophyll apoprotein; *fldA*, flavodoxin A; *pstS*, high-affinity phosphate transport system substrate-binding protein; SLC34A, sodium-dependent phosphate cotransporter; SQD1, UDP-sulfoquinovose synthase; *phoA*, alkaline phosphatase PhoA; ZIP123 and ZIP*, solute carrier family 39 (zinc transporter); ZCRP A/B, zinc/cobalt responsive proteins A/B.

**Fig. S15.**

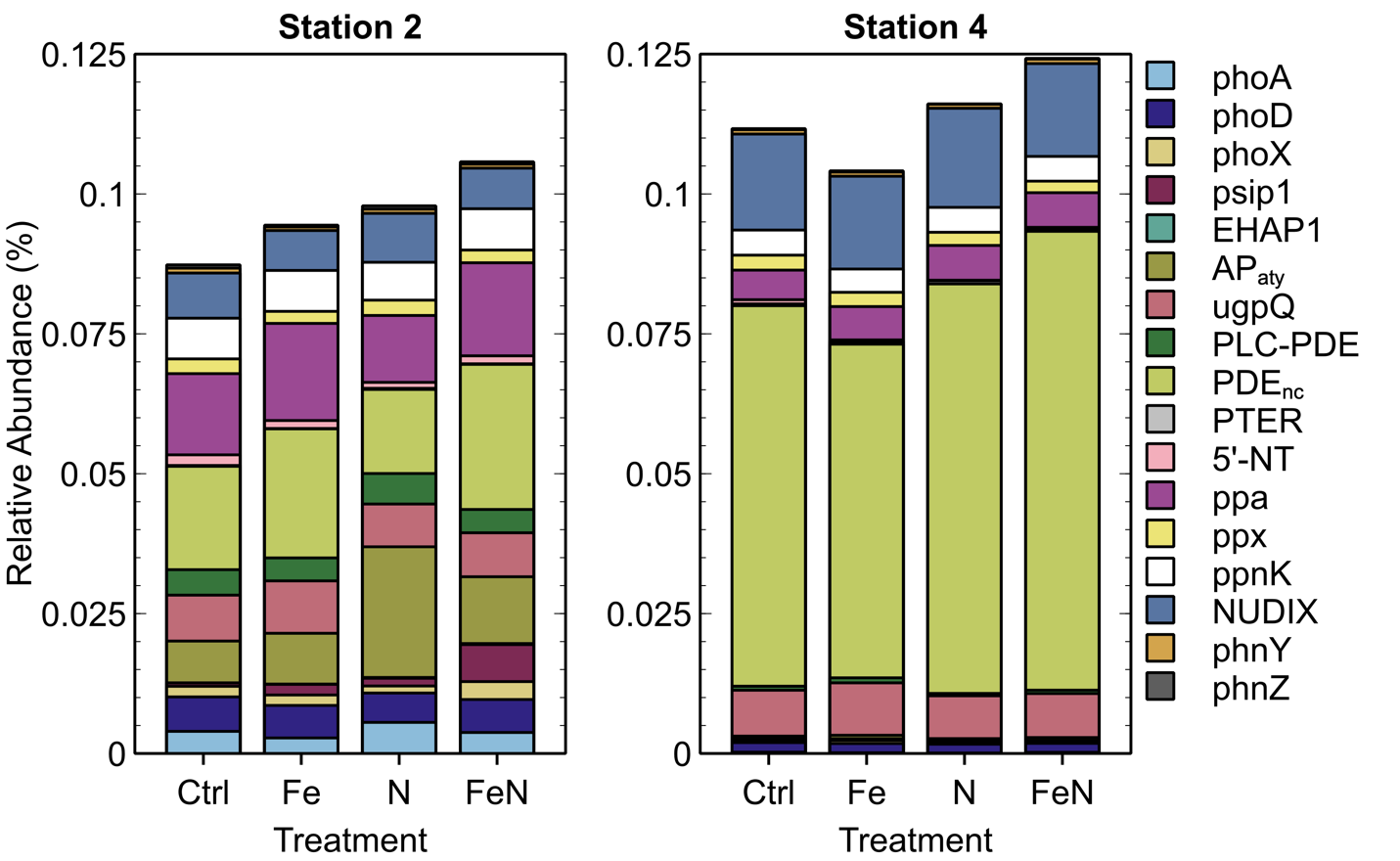

**Relative abundance of DOP-degrading enzymes during incubation bioassays**, indicated as relative abundance (%) of transcripts per million (TPM). *phoA*, alkaline phosphatase PhoA; *phoD*, alkaline phosphatase PhoD; *phoX*, alkaline phosphatase PhoX; psip1, alkaline phosphatase Psip1; EHAP1, *Emiliania huxleyi* AP1-type alkaline phosphatase; AP_aty_, atypical alkaline phosphatase; *ugpQ*, glycerophosphoryl diester phosphodiesterase; PLC-PDE, PLC-like phosphodiesterase; PDE_nc_, unclassified phosphodiesterase; PTER, phosphotriesterases; 5NT, 5’-nucleotidases; ppa, inorganic pyrophosphatase; ppx, exopolyphosphatase; NUDIC, (di)nucleoside polyphosphate hydrolase; *phnY*, 2-aminoethylphosphonate dioxygenase; *phnZ*, 2-amino-1-hydroxy-ethylphosphonate dioxygenase.

**Fig. S16.**

**A	Station 2 (Aged)**

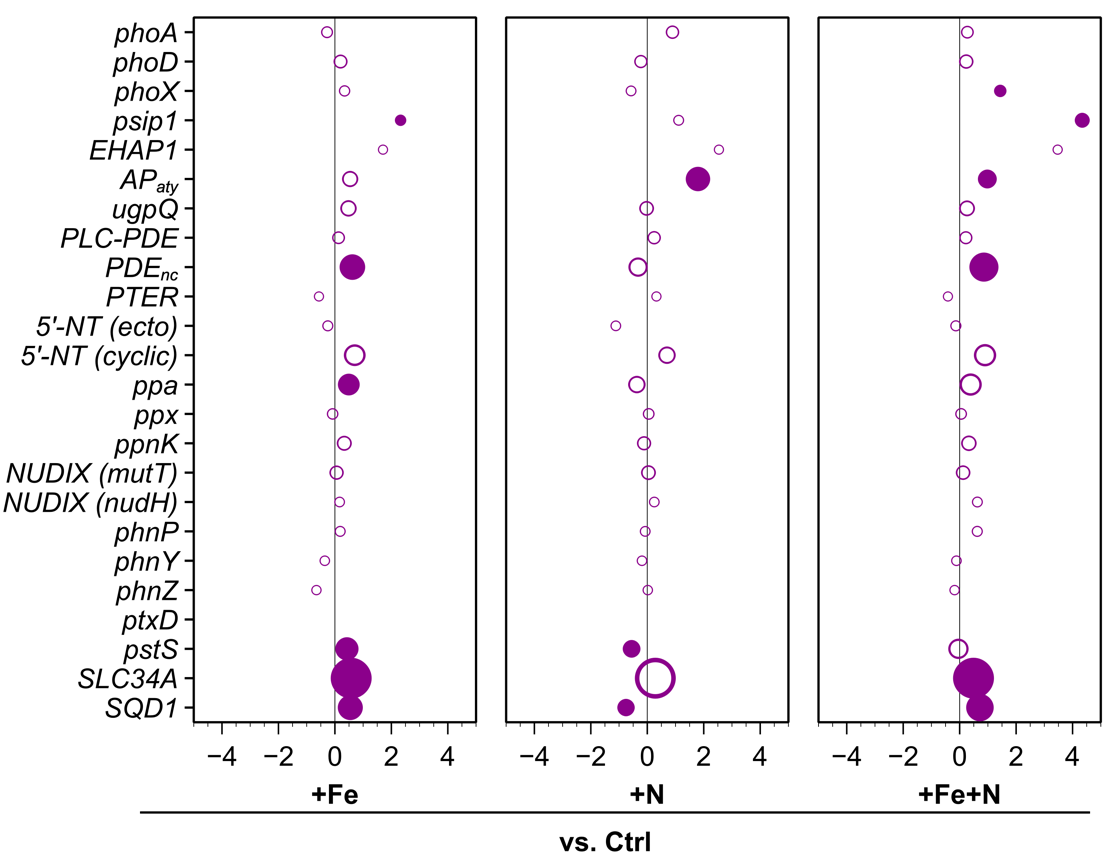

**B	Station 4 (Offshore)**

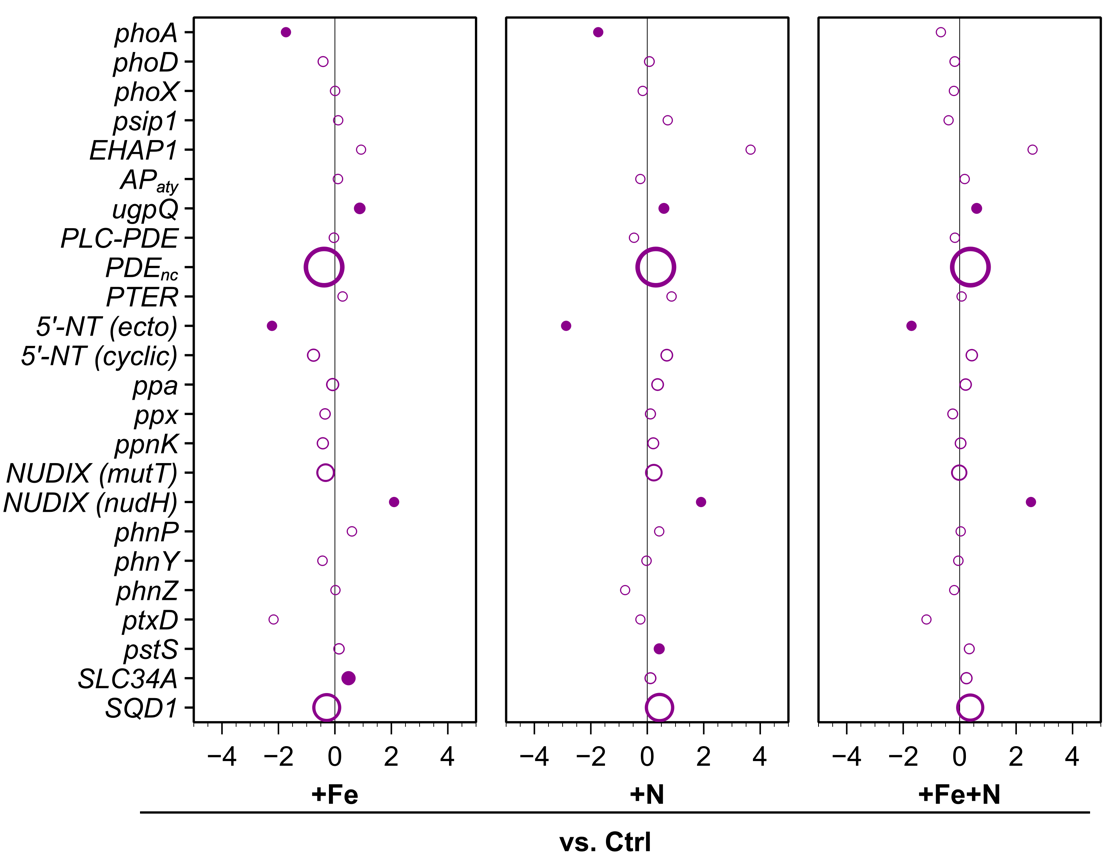

**Community-level differential expression of DOP utilizing enzymes within incubation experiments.** Differentially expressed transcripts are shown as log2-fold change proportion of expression for the eukaryotic microbial community. Circle sizes are weighted by relative abundance (%) of transcripts per million (TPM) normalized to scale between 0.5-2.0, and solid-colored circles denote statistical significance with an adjusted *p*-value < 0.05 (DESeq2 Wald test, Benjamini & Hochberg).

**Fig. S17.**

**A**

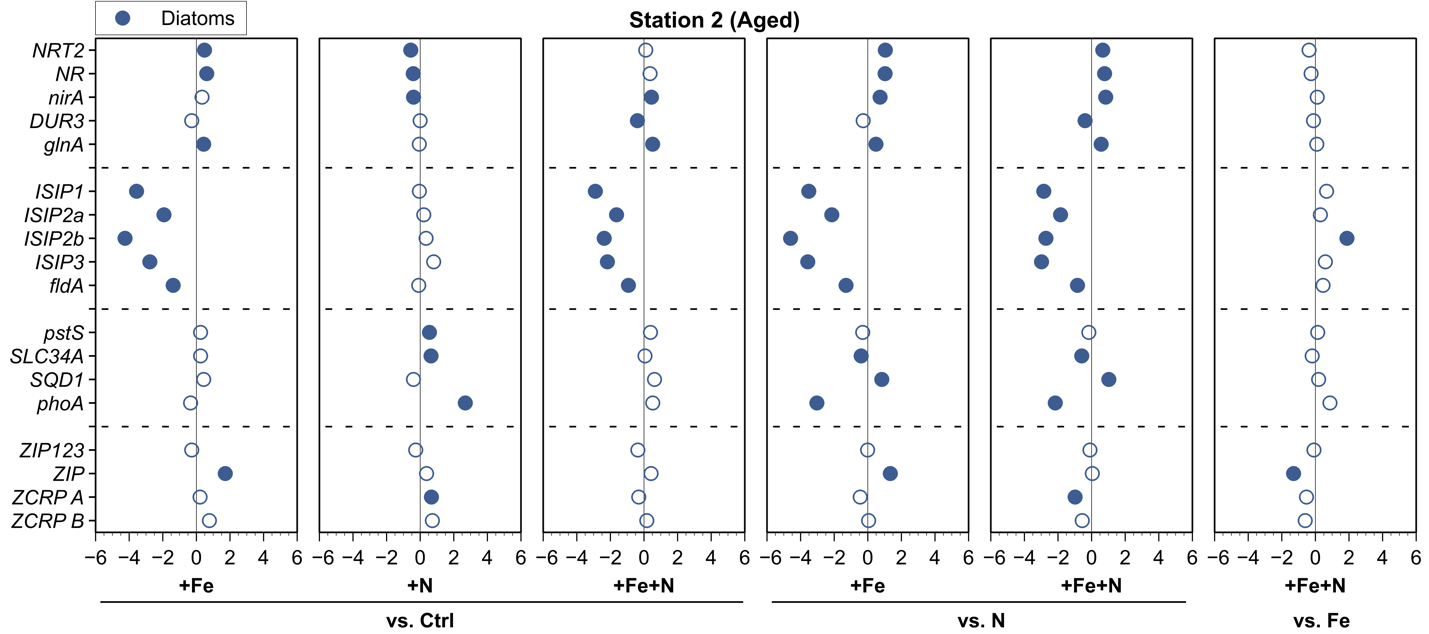

**B**

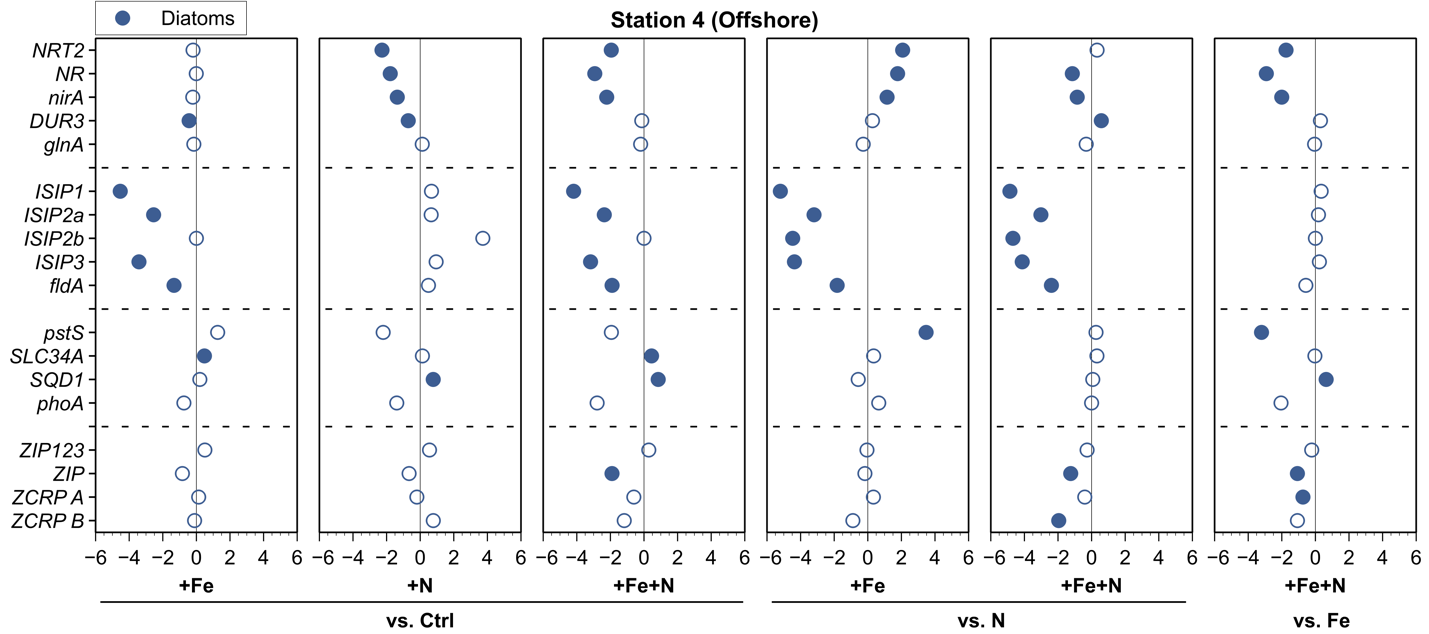

**C**

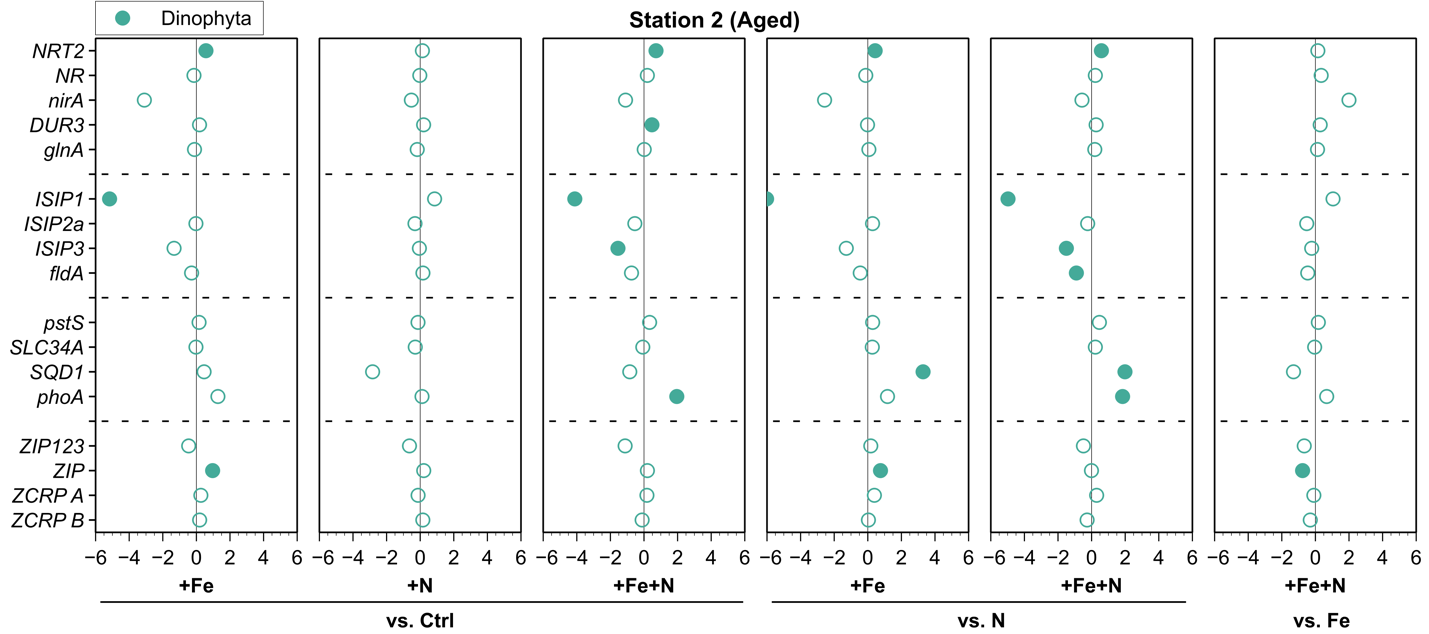

**D**

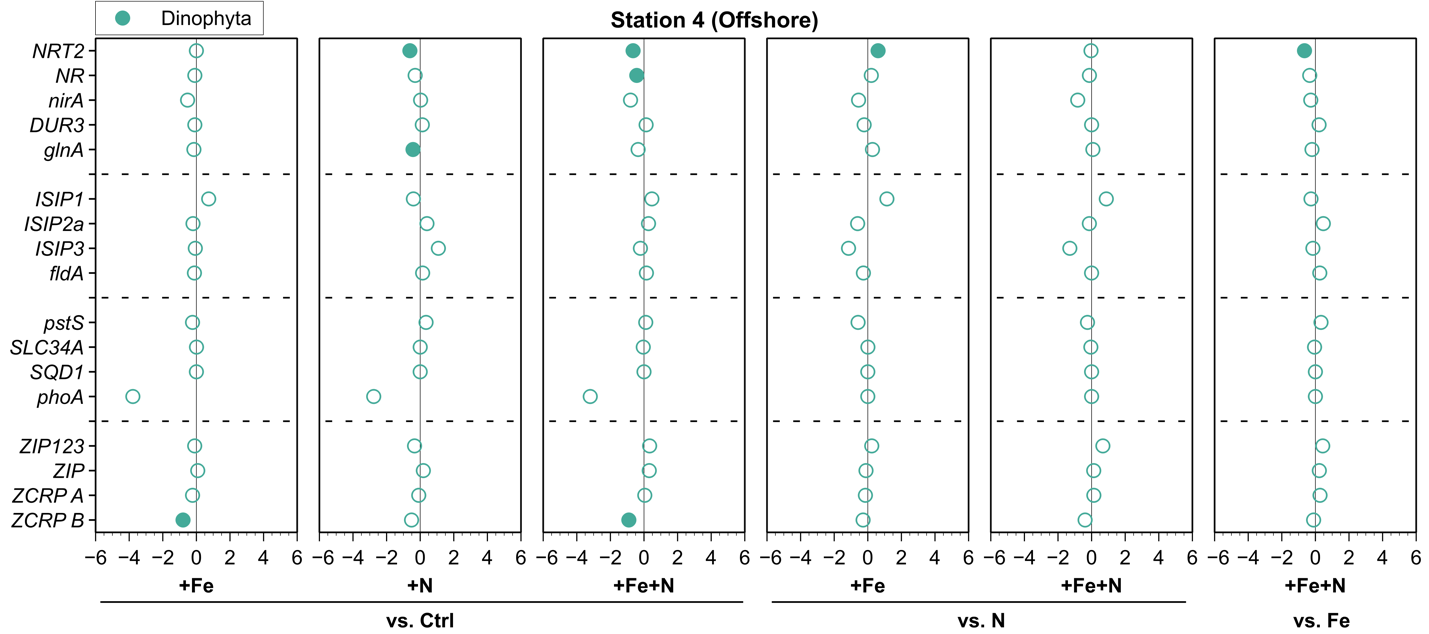

**E**

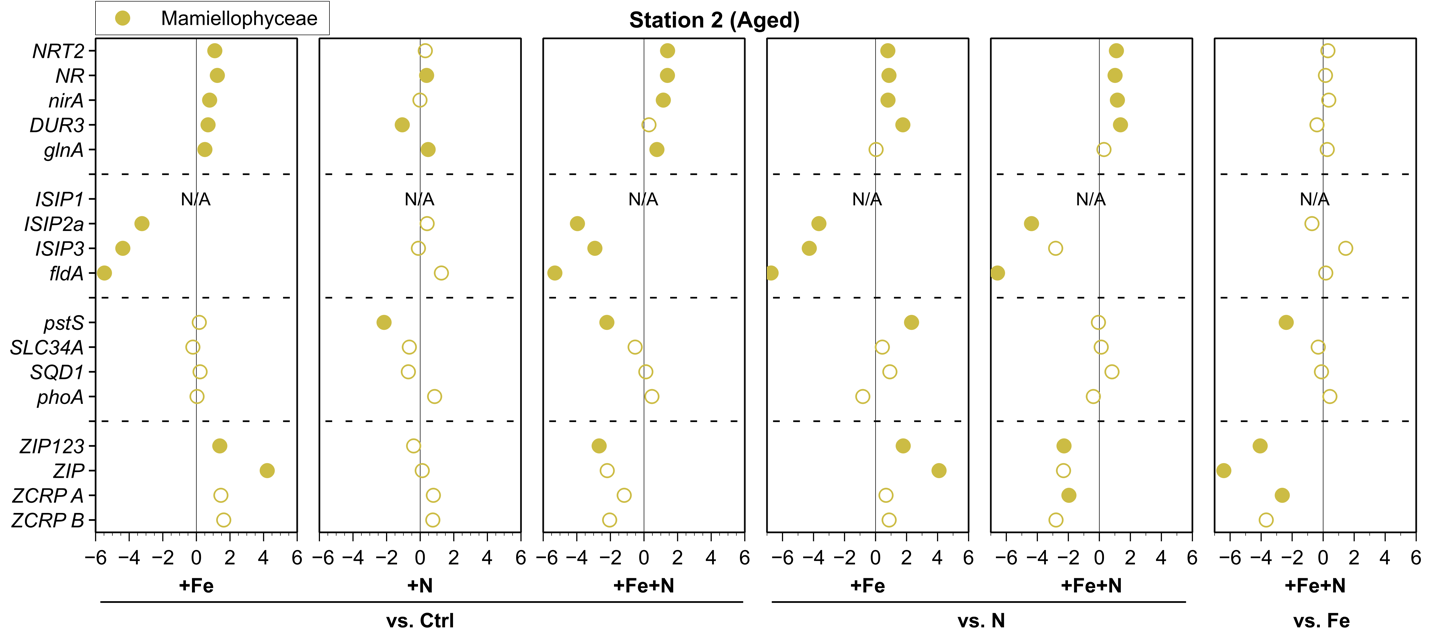

**F**

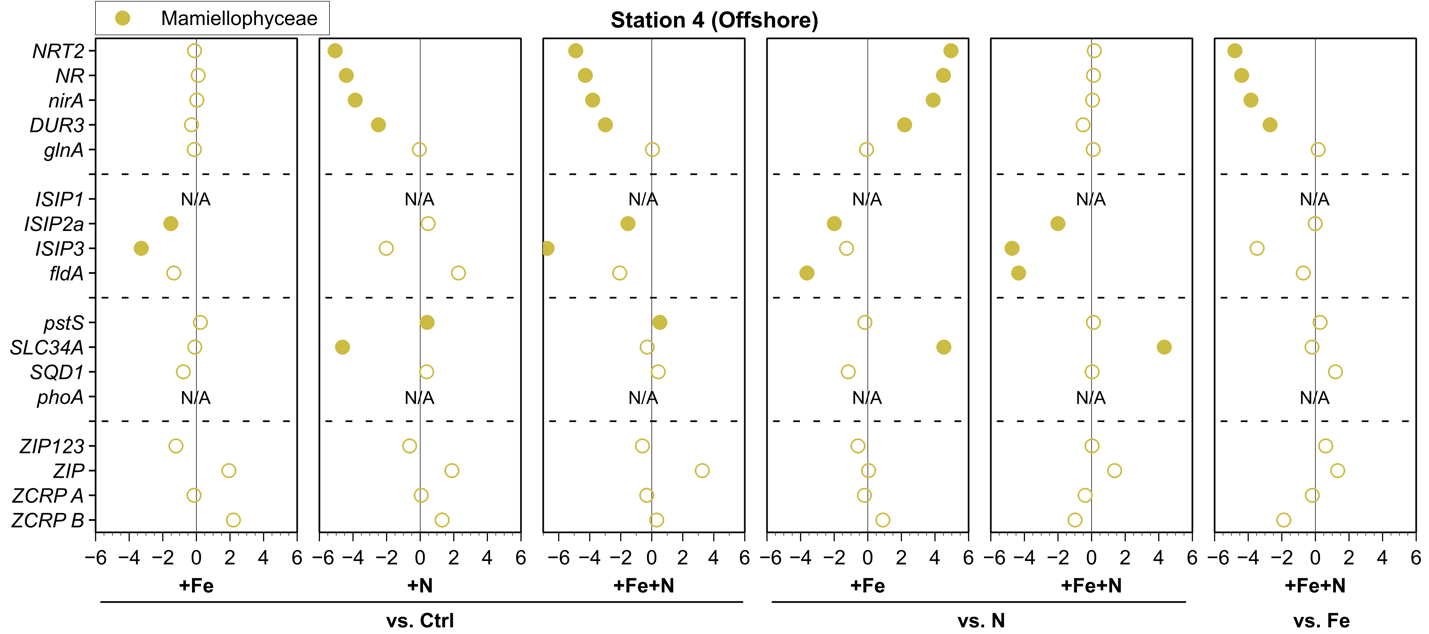

**G**

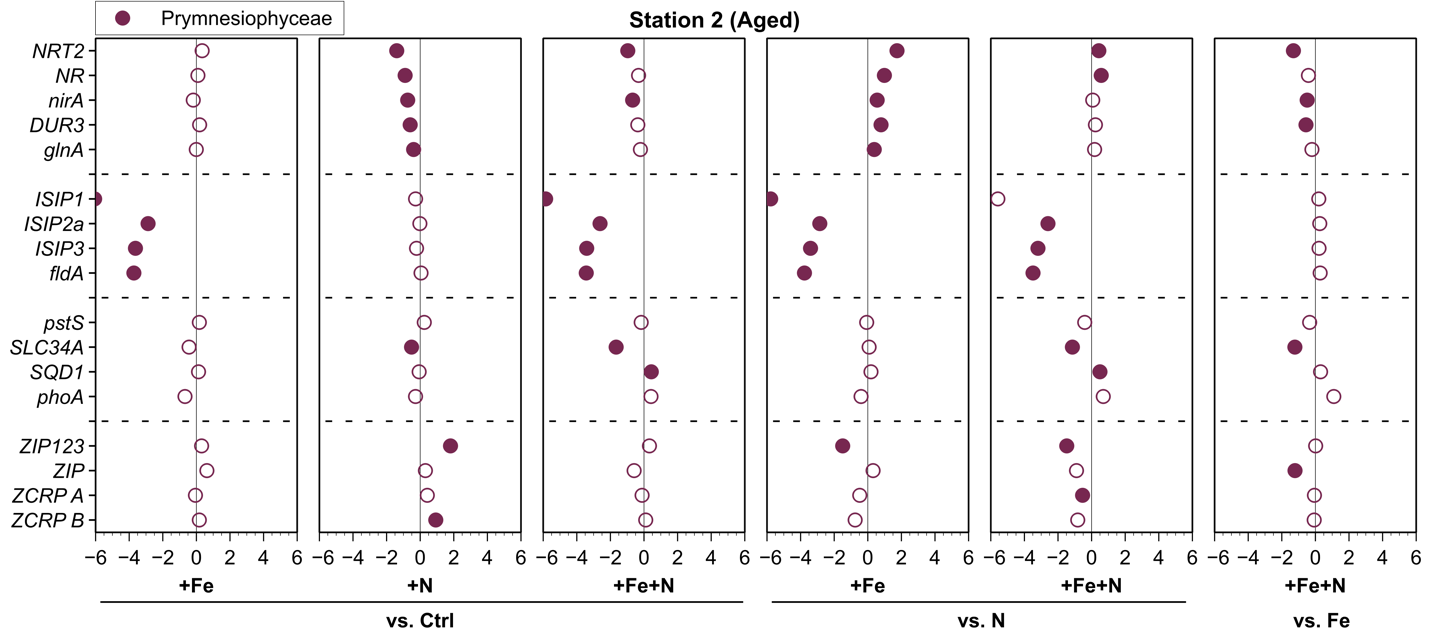

**H**

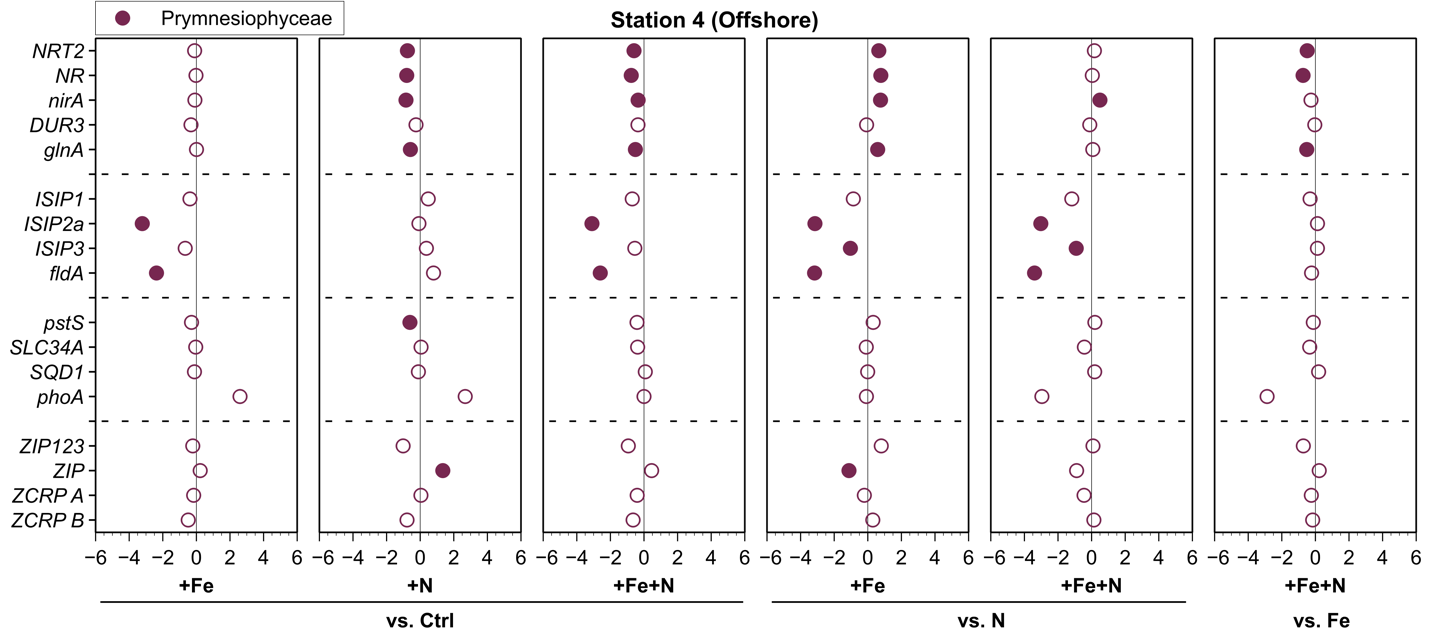

**I**

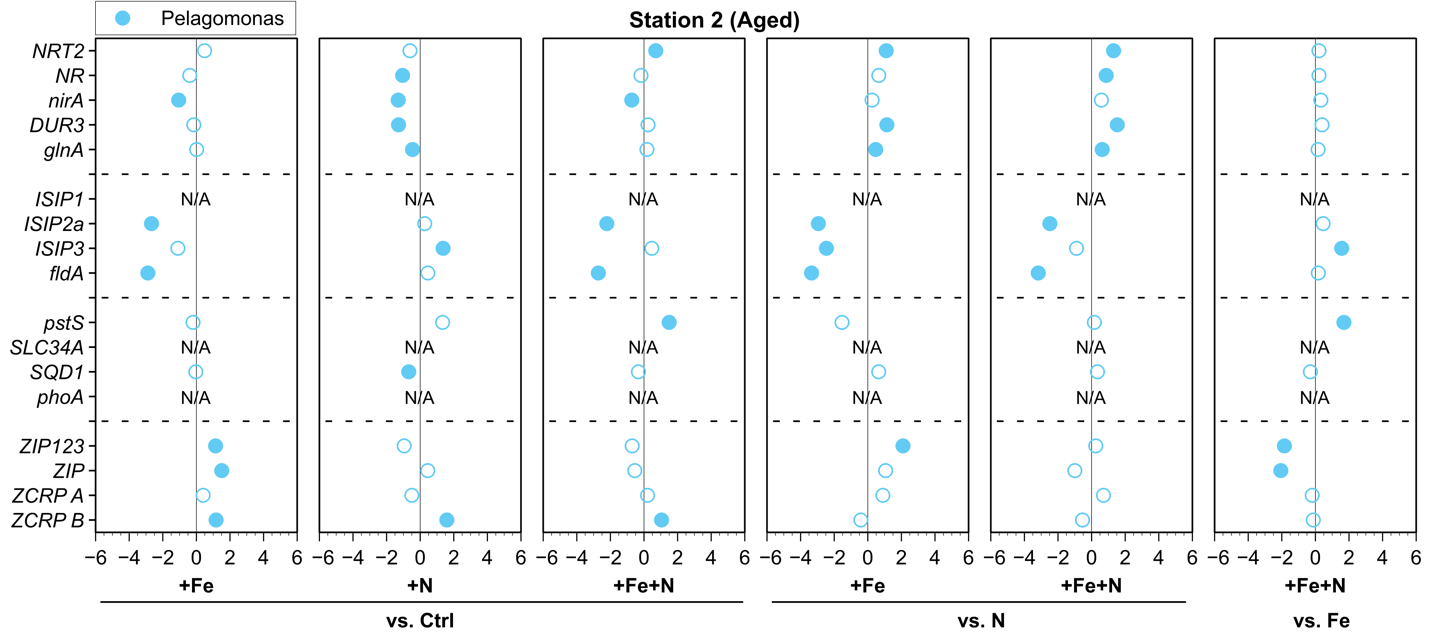

**J**

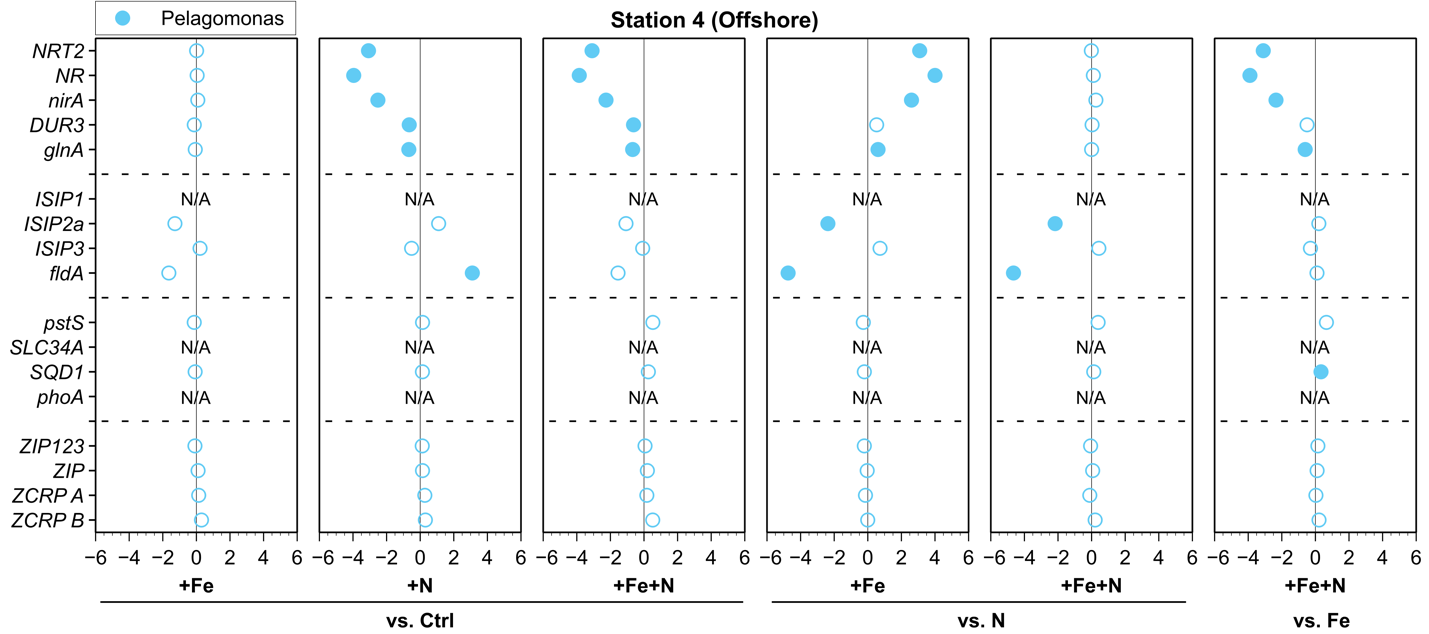

**Taxon-specific differential expression of nutrient stress biomarkers within incubation experiments.** Differentially expressed transcripts are shown as log2-fold change proportion of expression for major taxonomic groups diatoms (**A-B**), dinoflagellates (**C-D**), mamiellophytes (**E-F**), prymnesiophytes (**G-H**), and *Pelagomonas* (**I-J**), and solid-colored circles denote statistical significance with an adjusted *p*-value < 0.05 (DESeq2 Wald test, Benjamini & Hochberg).

**Fig. S18.**

**A**

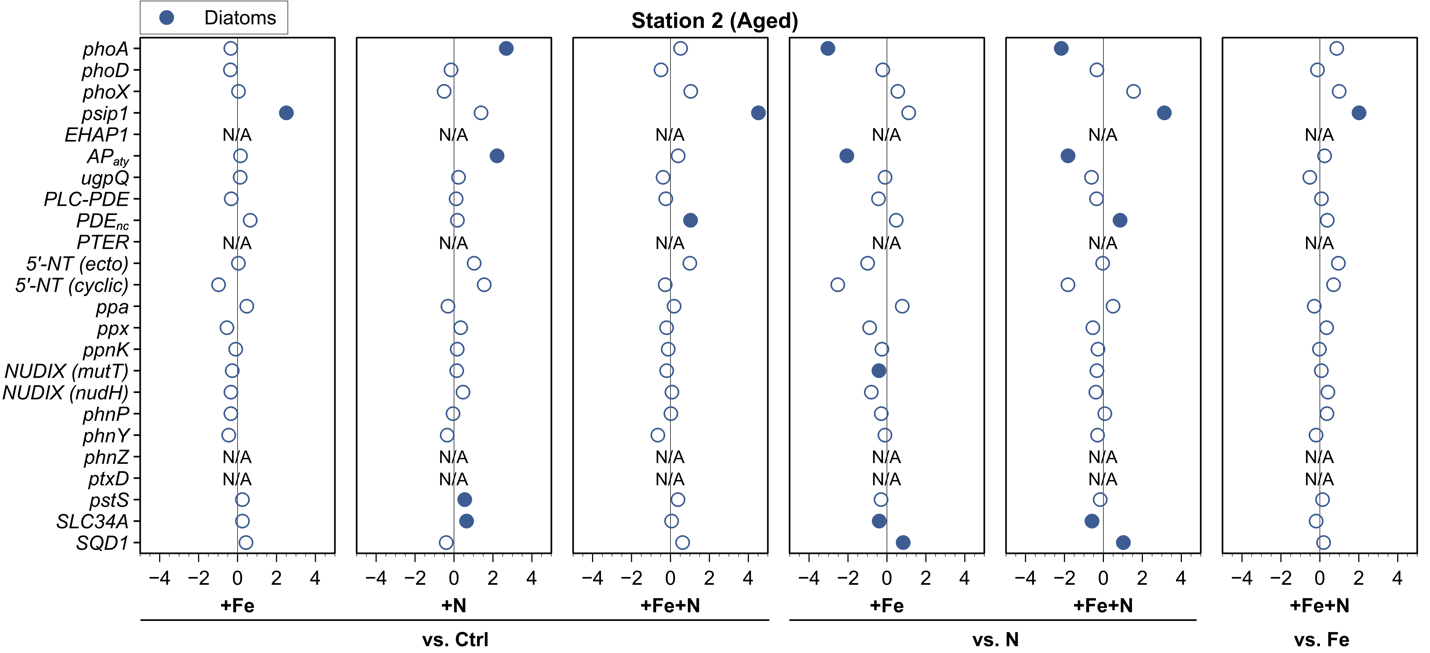

**B**

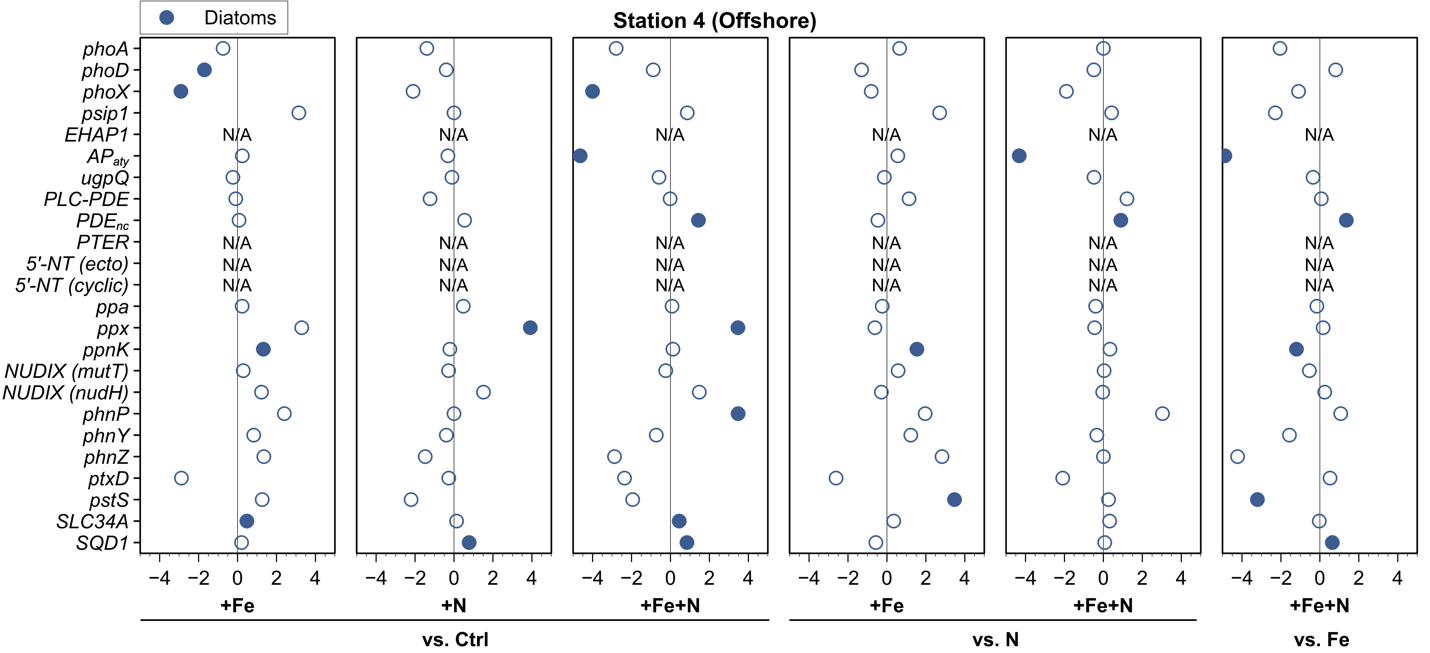

**C**

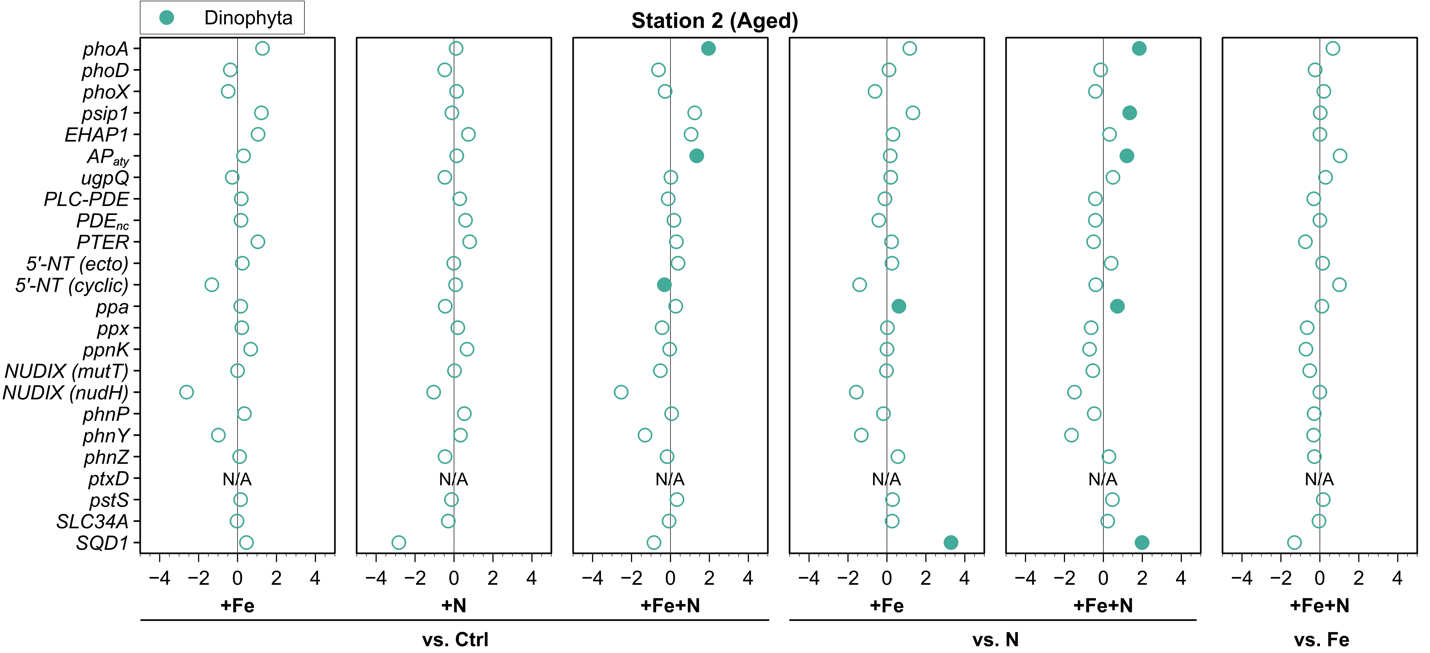

**D**

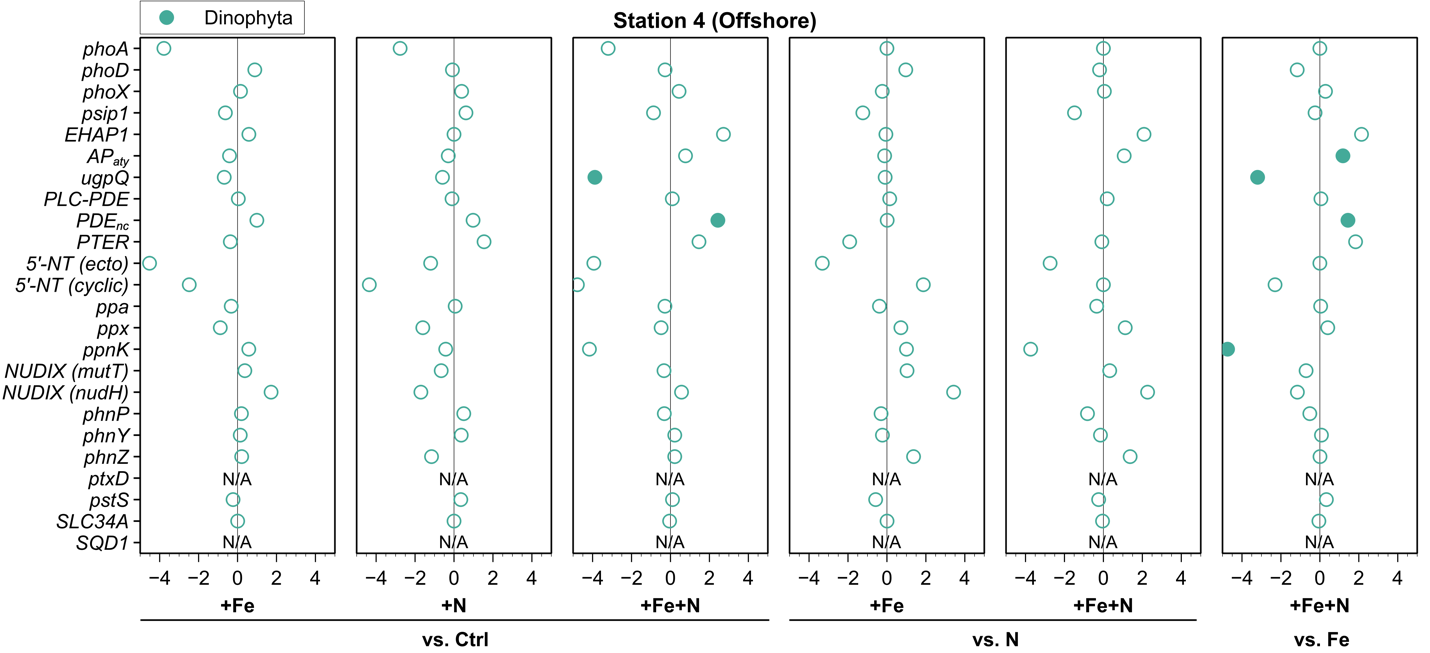

**E**

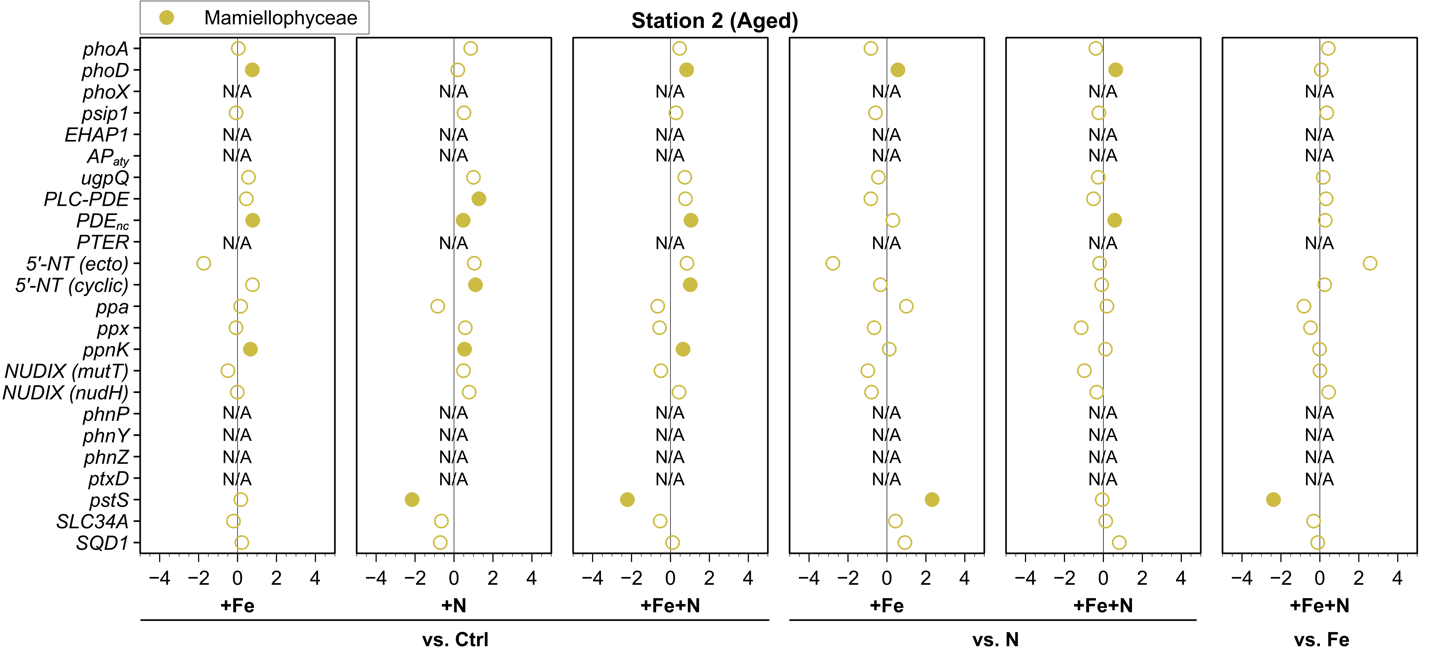

**F**

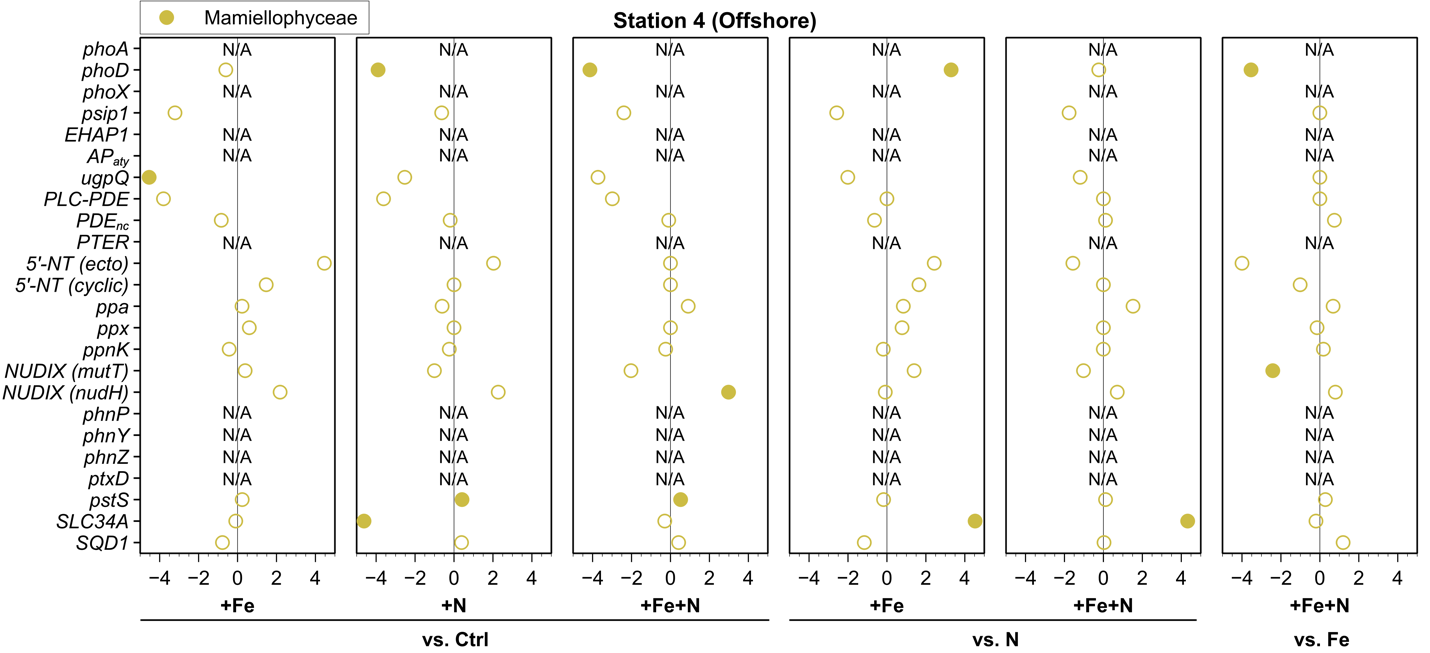

**G**

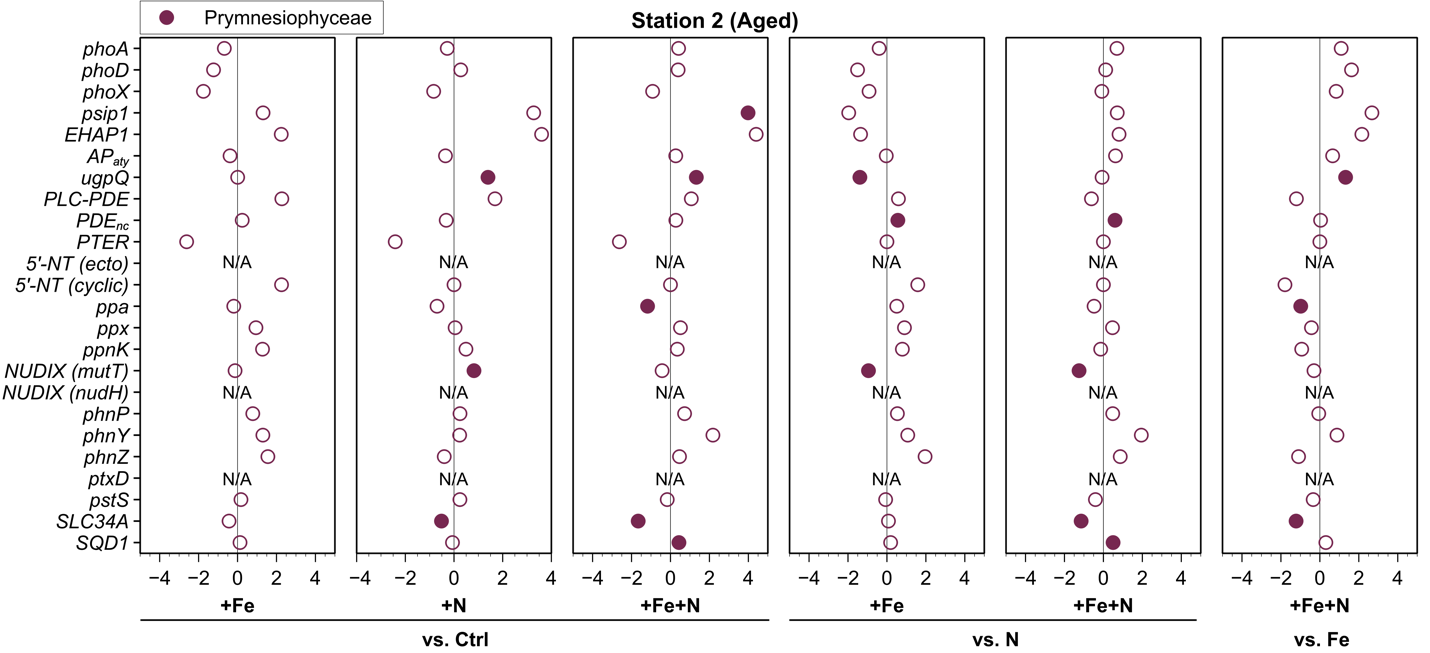

**H**

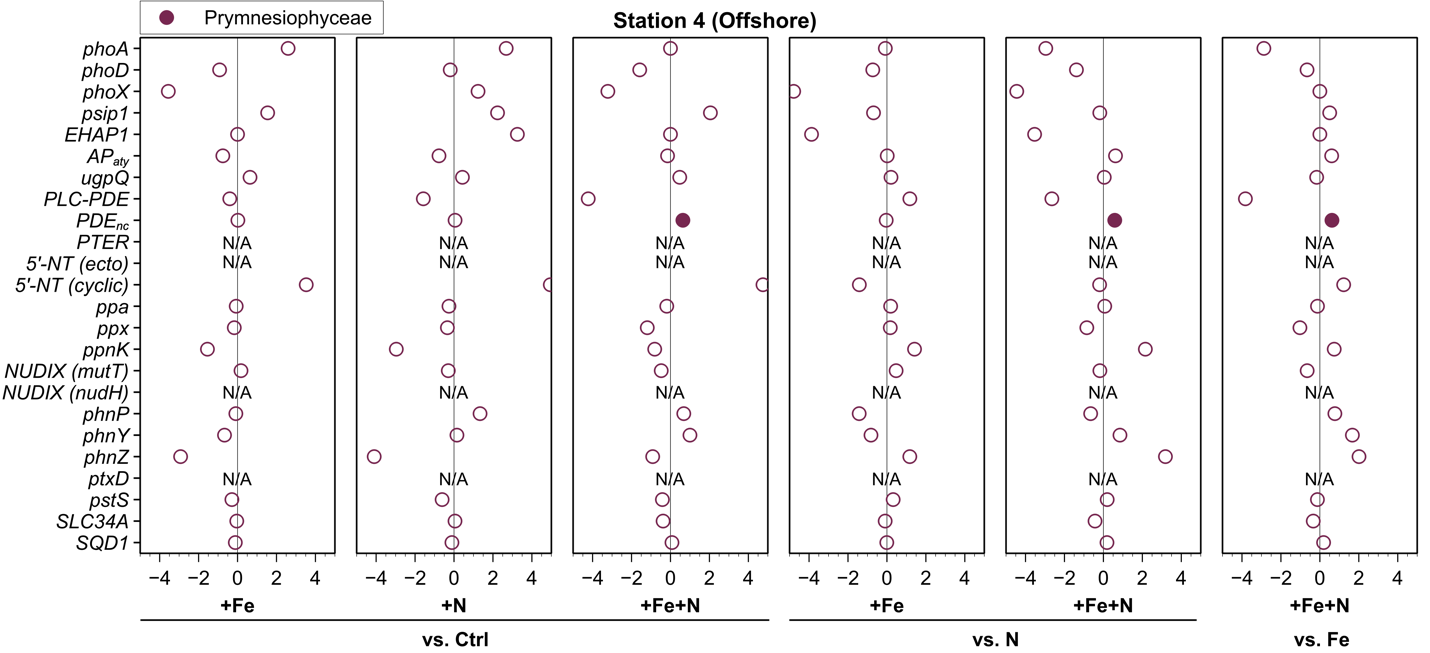

**I**

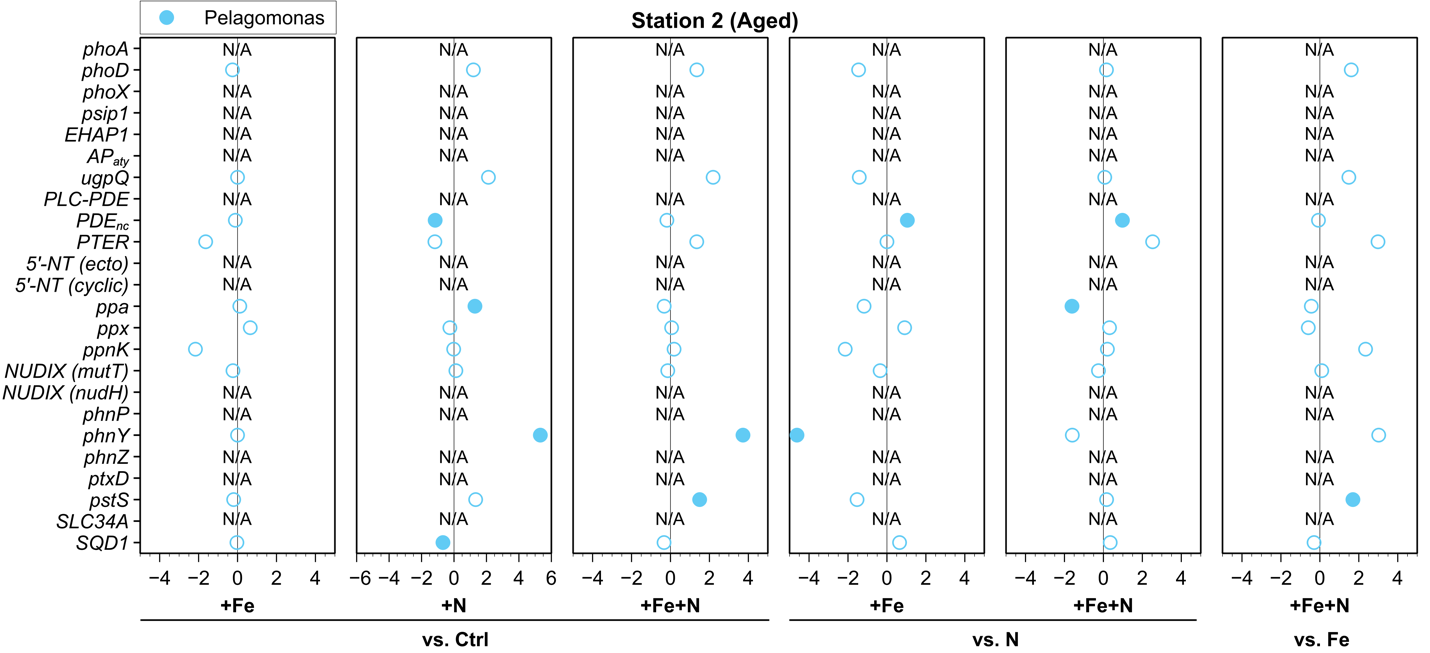

**J**

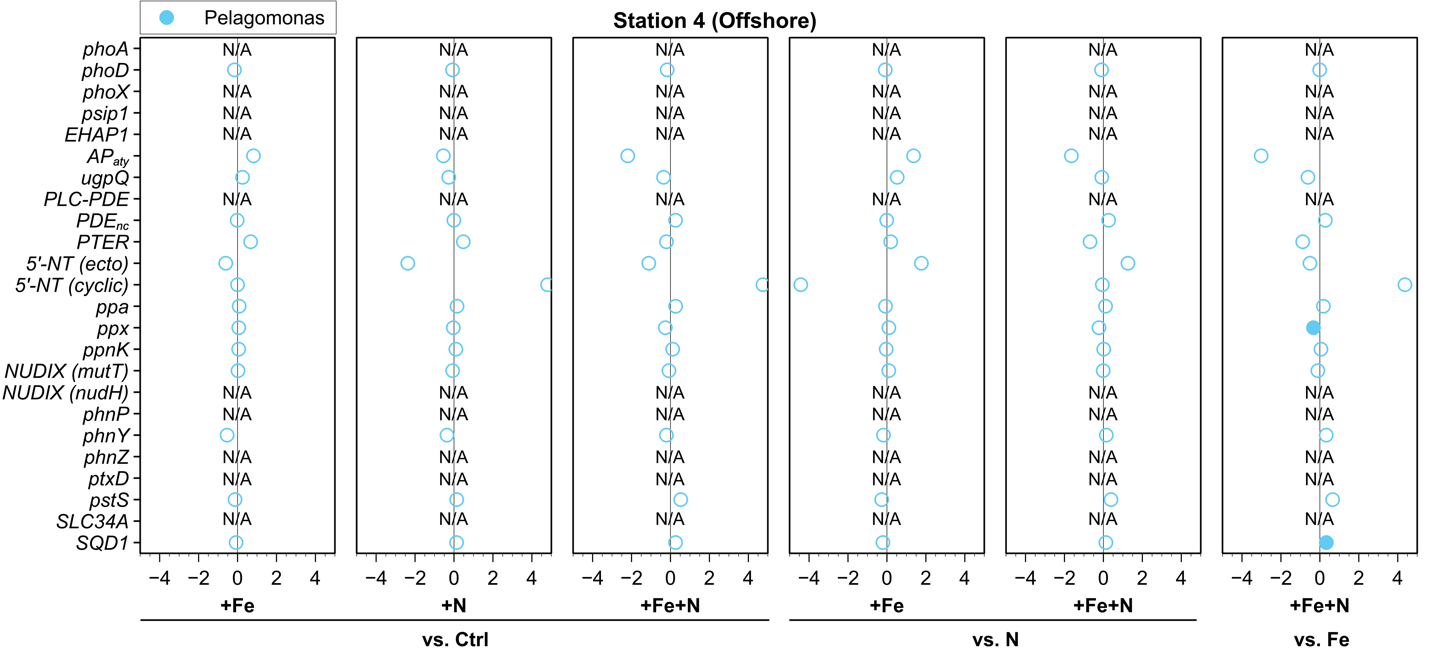

**Taxon-specific differential expression of DOP utilizing enzymes within incubation experiments.** Differentially expressed transcripts are shown as log2-fold change proportion of expression for major taxonomic groups diatoms (**A-B**), dinoflagellates (**C-D**), mamiellophytes (**E-F**), prymnesiophytes (**G-H**), and *Pelagomonas* (**I-J**), and solid-colored circles denote statistical significance with an adjusted *p*-value < 0.05 (DESeq2 Wald test, Benjamini & Hochberg).

**Fig. S19.**

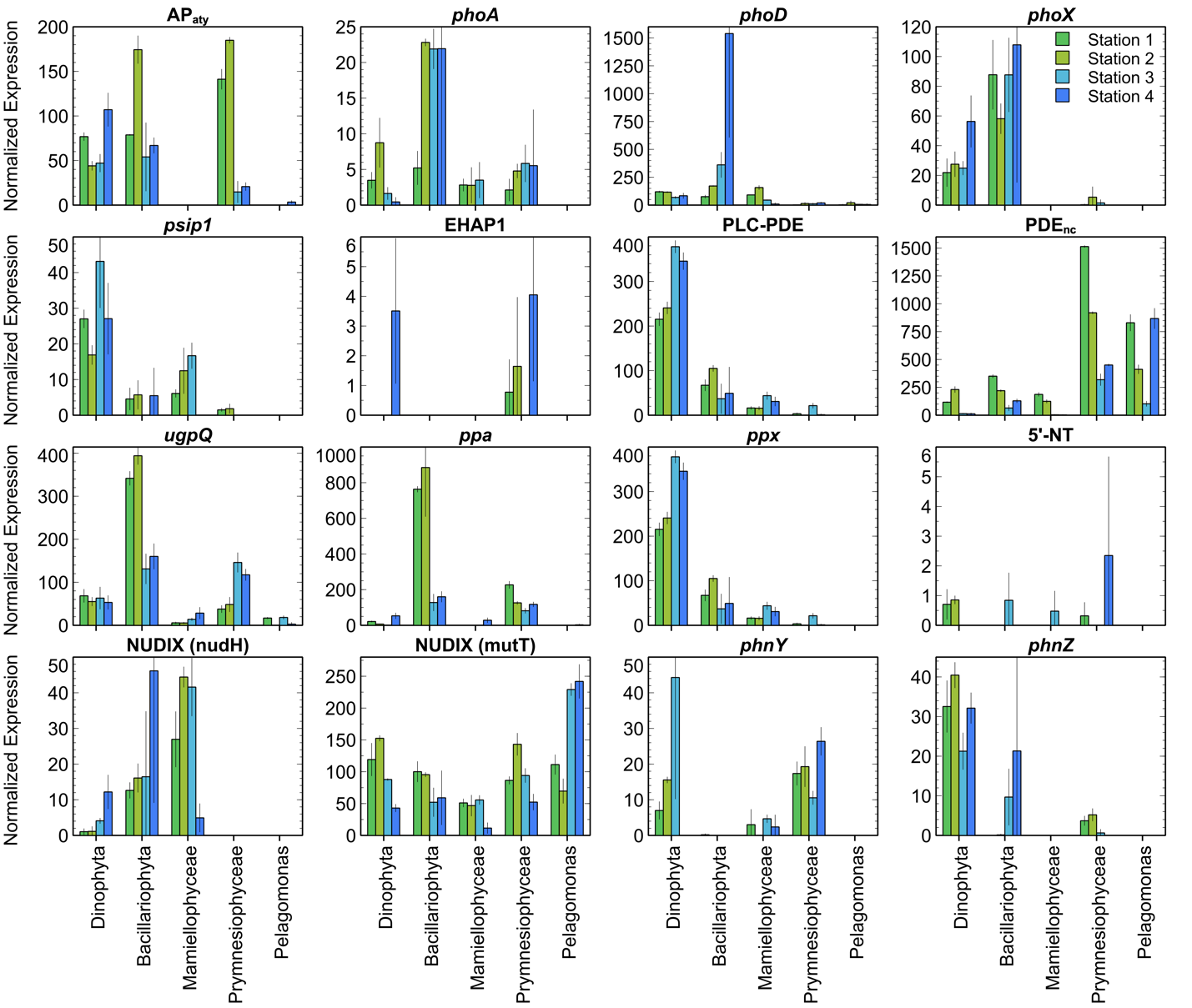

**Taxon-specific normalized transcript expression of DOP utilizing enzymes under ambient conditions in the CCE.** Expression is indicated as transcripts per million (TPM) normalized using DESeq2 v1.22.2 for each taxonomic group.

**Table S1. ISIP genes.** Sequences were assigned based on sequence similarity to literature references[32] via protein BLAST search (expectation value cutoff E 10^-25^).

| **Clade** | **Class** | **Species** | **Gene** |
| --- | --- | --- | --- |
| **ISIP1** | | | |
| Stramenopiles | Bacillariophyceae | *Phaeodactylum tricornutum* | [pti:PHATRDRAFT_55031](https://www.genome.jp/entry/pti:PHATRDRAFT_55031) |
| Stramenopiles | Bacillariophyceae | *Fragilariopsis cylindrus* | fcy:FRACYDRAFT_236570 |
| Stramenopiles | Bacillariophyceae | *Fragilariopsis cylindrus* | fcy:FRACYDRAFT_241515 |
| Stramenopiles | Bacillariophyceae | *Fragilariopsis cylindrus* | fcy:FRACYDRAFT_240619 |
| **ISIP2a** | | | |
| Stramenopiles | Bacillariophyceae | *Phaeodactylum tricornutum* | [pti:PHATRDRAFT_54465](https://www.genome.jp/entry/pti:PHATRDRAFT_54465) |
| Stramenopiles | Bacillariophyceae | *Phaeodactylum tricornutum* | pti:PHATRDRAFT_47240 |
| Stramenopiles | Bacillariophyceae | *Phaeodactylum tricornutum* | pti:PHATRDRAFT_54466 |
| Stramenopiles | Bacillariophyceae | *Phaeodactylum tricornutum* | pti:PHATRDRAFT_54708 |
| Stramenopiles | Bacillariophyceae | *Phaeodactylum tricornutum* | pti:PHATRDRAFT_49986 |
| Stramenopiles | Bacillariophyceae | *Phaeodactylum tricornutum* | pti:PHATRDRAFT_45708 |
| Stramenopiles | Bacillariophyceae | *Fragilariopsis cylindrus* | fcy:FRACYDRAFT_272749 |
| Stramenopiles | Bacillariophyceae | *Fragilariopsis cylindrus* | fcy:FRACYDRAFT_260427 |
| Stramenopiles | Bacillariophyceae | *Fragilariopsis cylindrus* | fcy:FRACYDRAFT_232971 |
| Stramenopiles | Dictyochophyceae | *Aureococcus anophagefferens* | aaf:AURANDRAFT_64252 |
| Hacrobia | Prymnesiophyceae | *Emiliania huxleyi* | ehx:EMIHUDRAFT_219446 |
| Hacrobia | Prymnesiophyceae | *Emiliania huxleyi* | ehx:EMIHUDRAFT_100149 |
| Hacrobia | Prymnesiophyceae | *Emiliania huxleyi* | ehx:EMIHUDRAFT_100764 |
| Hacrobia | Prymnesiophyceae | *Emiliania huxleyi* | ehx:EMIHUDRAFT_208161 |
| Hacrobia | Prymnesiophyceae | *Emiliania huxleyi* | ehx:EMIHUDRAFT_247589 |
| Hacrobia | Prymnesiophyceae | *Emiliania huxleyi* | ehx:EMIHUDRAFT_113079 |
| Hacrobia | Prymnesiophyceae | *Emiliania huxleyi* | ehx:EMIHUDRAFT_114466 |
| Hacrobia | Prymnesiophyceae | *Emiliania huxleyi* | ehx:EMIHUDRAFT_455896 |
| Hacrobia | Prymnesiophyceae | *Emiliania huxleyi* | ehx:EMIHUDRAFT_225221 |
| Chlorophyta | Mamiellophyceae | *Ostreococcus lucimarinus* | olu:OSTLU_26537 |
| **ISIP2b** | | | |
| Stramenopiles | Bacillariophyceae | *Phaeodactylum tricornutum* | [pti:PHATRDRAFT_54987](https://www.genome.jp/entry/pti:PHATRDRAFT_54987) |
| **ISIP3** | | | |
| Stramenopiles | Bacillariophyceae | *Thalassiosira pseudonana* | tps:THAPSDRAFT_bd1757 |
| Stramenopiles | Bacillariophyceae | *Phaeodactylum tricornutum* | [pti:PHATRDRAFT_47674](https://www.genome.jp/entry/pti:PHATRDRAFT_47674) |
| Stramenopiles | Bacillariophyceae | *Fragilariopsis cylindrus* | fcy:FRACYDRAFT_182258 |
| Stramenopiles | Bacillariophyceae | *Fragilariopsis cylindrus* | fcy:FRACYDRAFT_268337 |
| Hacrobia | Prymnesiophyceae | *Emiliania huxleyi* | ehx:EMIHUDRAFT_120809 |
| Hacrobia | Cryptophyceae | *Guillardia theta* | gtt:GUITHDRAFT_112878 |

**Table S2. ZCRP genes.** Sequences were assigned based on sequence similarity to literature references[33] via protein BLAST search (expectation value cutoff E 10^-25^).

| **Clade** | **Class** | **Species** | **Gene** |
| --- | --- | --- | --- |
| **ZCRP_A** | | | |
| Alveolata | [Dinophyceae](https://en.wikipedia.org/wiki/Dinophyceae) | *Breviolum minutum* | smin:v1.2.026665.t1 |
| Alveolata | [Dinophyceae](https://en.wikipedia.org/wiki/Dinophyceae) | *Breviolum minutum* | smin:v1.2.015561.t1 |
| Alveolata | [Dinophyceae](https://en.wikipedia.org/wiki/Dinophyceae) | *Breviolum minutum* | smin:v1.2.024197.t2 |
| Alveolata | [Dinophyceae](https://en.wikipedia.org/wiki/Dinophyceae) | *Breviolum minutum* | smin:v1.2.035355.t1 |
| Alveolata | [Dinophyceae](https://en.wikipedia.org/wiki/Dinophyceae) | *Breviolum minutum* | smin:v1.2.035355.t2 |
| Stramenopiles | Bacillariophyceae | *Thalassiosira pseudonana* | [tps:THAPSDRAFT_3054](https://www.genome.jp/entry/tps:THAPSDRAFT_3054) |
| Stramenopiles | Bacillariophyceae | *Phaeodactylum tricornutum* | [pti:PHATRDRAFT_41508](https://www.genome.jp/entry/pti:PHATRDRAFT_41508) |
| Stramenopiles | Bacillariophyceae | *Fragilariopsis cylindrus* | fcy:FRACYDRAFT_224667 |
| Stramenopiles | Bacillariophyceae | *Fragilariopsis cylindrus* | fcy:FRACYDRAFT_184852 |
| Stramenopiles | Dictyochophyceae | *Aureococcus anophagefferens* | aaf:AURANDRAFT_12902 |
| Stramenopiles | Dictyochophyceae | *Aureococcus anophagefferens* | aaf:AURANDRAFT_52124 |
| Stramenopiles | Dictyochophyceae | *Aureococcus anophagefferens* | aaf:AURANDRAFT_12916 |
| Stramenopiles | Dictyochophyceae | *Aureococcus anophagefferens* | aaf:AURANDRAFT_60046 |
| Stramenopiles | Dictyochophyceae | *Aureococcus anophagefferens* | aaf:AURANDRAFT_12944 |
| Stramenopiles | Dictyochophyceae | *Aureococcus anophagefferens* | aaf:AURANDRAFT_64741 |
| Hacrobia | Prymnesiophyceae | *Emiliania huxleyi* | ehx:EMIHUDRAFT_419514 |
| Hacrobia | Prymnesiophyceae | *Emiliania huxleyi* | ehx:EMIHUDRAFT_456427 |
| Hacrobia | Prymnesiophyceae | *Emiliania huxleyi* | ehx:EMIHUDRAFT_68047 |
| Hacrobia | Prymnesiophyceae | *Emiliania huxleyi* | ehx:EMIHUDRAFT_465373 |
| Hacrobia | Prymnesiophyceae | *Emiliania huxleyi* | ehx:EMIHUDRAFT_43480 |
| Hacrobia | Prymnesiophyceae | *Emiliania huxleyi* | ehx:EMIHUDRAFT_54553 |
| Hacrobia | Prymnesiophyceae | *Emiliania huxleyi* | ehx:EMIHUDRAFT_417487 |
| Hacrobia | Cryptophyceae | *Guillardia theta* | gtt:GUITHDRAFT_158114 |
| Hacrobia | Cryptophyceae | *Guillardia theta* | gtt:GUITHDRAFT_157306 |
| Chlorophyta | Chlorophyceae | *Volvox carteri* | vcn:VOLCADRAFT_107727 |
| Chlorophyta | Chlorophyceae | *Chlorella variabilis* | cvr:CHLNCDRAFT_142112 |
| Chlorophyta | Mamiellophyceae | *Bathycoccus prasinos* | bpg:Bathy14g01250 |
| Chlorophyta | Mamiellophyceae | *Micromonas commoda* | mis:MICPUN_108724 |
| Chlorophyta | Mamiellophyceae | *Micromonas commoda* | mis:MICPUN_94775 |
| Chlorophyta | Mamiellophyceae | *Micromonas commoda* | mis:MICPUN_60337 |
| Chlorophyta | Mamiellophyceae | *Micromonas pusilla* | mpp:MICPUCDRAFT_30963 |
| Chlorophyta | Mamiellophyceae | *Ostreococcus lucimarinus* | olu:OSTLU_29990 |
| Chlorophyta | Mamiellophyceae | *Ostreococcus lucimarinus* | olu:OSTLU_40657 |
| Chlorophyta | Mamiellophyceae | *Ostreococcus tauri* | ota:OT_ostta02g04860 |
| Chlorophyta | Mamiellophyceae | *Ostreococcus tauri* | ota:OT_ostta06g02290 |
| **ZCRP_B** | | | |
| Alveolata | [Dinophyceae](https://en.wikipedia.org/wiki/Dinophyceae) | *Breviolum minutum* | smin:v1.2.024646.t1 |
| Alveolata | [Dinophyceae](https://en.wikipedia.org/wiki/Dinophyceae) | *Breviolum minutum* | smin:v1.2.024646.t2 |
| Stramenopiles | Bacillariophyceae | *Thalassiosira pseudonana* | [tps:THAPSDRAFT_bd938](https://www.genome.jp/entry/tps:THAPSDRAFT_bd938) |
| Stramenopiles | Bacillariophyceae | *Phaeodactylum tricornutum* | [pti:PHATRDRAFT_49168](https://www.genome.jp/entry/pti:PHATRDRAFT_49168) |
| Stramenopiles | Dictyochophyceae | *Aureococcus anophagefferens* | aaf:AURANDRAFT_8505 |
| Hacrobia | Prymnesiophyceae | *Emiliania huxleyi* | ehx:EMIHUDRAFT_113427 |
| Hacrobia | Prymnesiophyceae | *Emiliania huxleyi* | ehx:EMIHUDRAFT_198033 |

**Table S3. Alkaline phosphatase genes.** Sequences were combined from main KEGG orthologies, literature references as indicated in the table, and manually assigned genes (denoted ‘M’) based on sequence similarity to references via protein BLAST search (expectation value cutoff E 10^-25^), Pfam (protein family) or CDD (conserved domain database) classification, and multiple sequence alignments. Literature sequences without entries in the KEGG GENES database were omitted.

| **Clade** | **Class** | **Species** | **Gene** | **Type** | **Ref.** |
| --- | --- | --- | --- | --- | --- |
| **phoA (KO K01077)** | | | | | |
| Stramenopiles | Bacillariophyceae | *Thalassiosira pseudonana* | tps:THAPSDRAFT_261067 | PhoA | ^[34,35]^ |
| Stramenopiles | Bacillariophyceae | *Fragilariopsis cylindrus* | fcy:FRACYDRAFT_187960 | PhoA | M |
| Stramenopiles | Bacillariophyceae | *Phaeodactylum tricornutum* | pti:PHATRDRAFT_49678 | PhoA | ^[36,37]^ |
| **phoD (KO K01113)** | | | | | |
| Stramenopiles | Bacillariophyceae | *Thalassiosira pseudonana* | tps:THAPSDRAFT_20880 | PhoD | ^[34]^ |
| Hacrobia | Cryptophyceae | *Guillardia theta* | gtt:GUITHDRAFT_140586 | PhoD | M |
| **phoX (KO K07093)** | | | | | |
| Hacrobia | Cryptophyceae | *Guillardia theta* | gtt:GUITHDRAFT_110221 | PhoX | M |
| Chlorophyta | Chlorophyceae | *Volvox carteri* | vcn:VOLCADRAFT_74874 | PhoX | ^[38]^ |
| Chlorophyta | Chlorophyceae | *Chlamydomonas reinhardtii* | cre:CHLRE_04g216700v5 | PhoX | ^[39]^ |
| Chlorophyta | Mamiellophyceae | *Micromonas commoda* | mis:MICPUN_64401 | PhoX | ^[40]^ |
| Chlorophyta | Trebouxiophyceae | *Coccomyxa subellipsoidea* | csl:COCSUDRAFT_42448 | PhoX | M |
| Chlorophyta | Chlorophyceae | *Monoraphidium neglectum* | mng:MNEG_4949 | PhoX | M |
| Chlorophyta | Chlorophyceae | *Monoraphidium neglectum* | mng:MNEG_4948 | PhoX | M |
| Chlorophyta | Chlorophyceae | *Monoraphidium neglectum* | mng:MNEG_4947 | PhoX | M |
| Chlorophyta | Chlorophyceae | *Chlorella variabilis* | cvr:CHLNCDRAFT_135377 | PhoX | M |
| Chlorophyta | Chlorophyceae | *Chlorella variabilis* | cvr:CHLNCDRAFT_135376 | PhoX | M |
| **EHAP1** | | | | | |
| Hacrobia | Prymnesiophyceae | *Emiliania huxleyi* | ehx:EMIHUDRAFT_433041 | EH-AP1 | ^[41]^ |
| Hacrobia | Prymnesiophyceae | *Emiliania huxleyi* | ehx:EMIHUDRAFT_414308 |  | ^[41]^ |
| **AP_aty_** | | | | | |
| Stramenopiles | Bacillariophyceae | *Phaeodactylum tricornutum* | pti:PHATRDRAFT_47869 | aty. | ^[36,37,42]^ |
| Stramenopiles | Bacillariophyceae | *Phaeodactylum tricornutum* | pti:PHATRDRAFT_47612 | aty. | ^[36,37]^ |
| Stramenopiles | Bacillariophyceae | *Thalassiosira pseudonana* | tps:THAPSDRAFT_260835 | aty. | ^[34,35]^ |
| Stramenopiles | Bacillariophyceae | *Thalassiosira pseudonana* | tps:THAPSDRAFT_8004 | aty. | ^[34]^ |
| Stramenopiles | Bacillariophyceae | *Fragilariopsis cylindrus* | fcy:FRACYDRAFT_206408 | aty. | M |
| Stramenopiles | Dictyochophyceae | *Aureococcus anophagefferens* | aaf:AURANDRAFT_70668 | aty. | ^[43]^ |
| Hacrobia | Prymnesiophyceae | *Emiliania huxleyi* | ehx:EMIHUDRAFT_444279 | aty. | ^[42,44]^ |
| Hacrobia | Prymnesiophyceae | *Emiliania huxleyi* | ehx:EMIHUDRAFT_461703 | aty. | ^[44]^ |
| Hacrobia | Prymnesiophyceae | *Emiliania huxleyi* | ehx:EMIHUDRAFT_235275 | aty. | M |
| Hacrobia | Prymnesiophyceae | *Emiliania huxleyi* | ehx:EMIHUDRAFT_469213 | aty. | M |
| Chlorophyta | Chlorophyceae | *Chlamydomonas reinhardtii* | cre:CHLRE_08g359300v5 | aty. | ^[42]^ |
| Chlorophyta | Chlorophyceae | *Volvox carteri* | vcn:VOLCADRAFT_119898 | aty. | ^[42]^ |

**Table S4. Phosphodiesterase genes.** Sequences were combined from main KEGG orthologies, literature references as indicated in the table, and manually assigned genes (denoted ‘M’) based on sequence similarity to references via protein BLAST search (expectation value cutoff E 10^-25^), Pfam (protein family) or CDD (conserved domain database) classification, and multiple sequence alignments. Literature sequences without entries in the KEGG GENES database were omitted.

| **Clade** | **Class** | **Species** | **Gene** | **Ref.** |
| --- | --- | --- | --- | --- |
| **ugpQ (KO K01126)** | | | | |
| Stramenopiles | Bacillariophyceae | *Phaeodactylum tricornutum* | pti:PHATRDRAFT_32057 | ^[45]^ |
| Stramenopiles | Bacillariophyceae | *Phaeodactylum tricornutum* | pti:PHATRDRAFT_44900 | ^[45]^ |
| Stramenopiles | Bacillariophyceae | *Thalassiosira pseudonana* | tps:THAPSDRAFT_23858 | ^[34,35]^ |
| Stramenopiles | Bacillariophyceae | *Fragilariopsis cylindrus* | fcy:FRACYDRAFT_244085 | M |
| Stramenopiles | Bacillariophyceae | *Fragilariopsis cylindrus* | fcy:FRACYDRAFT_255557 | M |
| **PLC-PDE** | | | | |
| Stramenopiles | Bacillariophyceae | *Thalassiosira pseudonana* | tps:THAPSDRAFT_23896 | ^[35]^ |
| Stramenopiles | Bacillariophyceae | *Fragilariopsis cylindrus* | fcy:FRACYDRAFT_179257 | M |
| Stramenopiles | Bacillariophyceae | *Thalassiosira pseudonana* | tps:THAPSDRAFT_21757 | ^[34]^ |
| **PDE_nc_** | | | | |
| Stramenopiles | Bacillariophyceae | *Thalassiosira pseudonana* | tps:THAPSDRAFT_907 | ^[34]^ |
| Stramenopiles | Bacillariophyceae | *Phaeodactylum tricornutum* | pti:PHATR_43978 | M |
| Stramenopiles | Bacillariophyceae | *Phaeodactylum tricornutum* | pti:PHATRDRAFT_48680 | M |
| Stramenopiles | Bacillariophyceae | *Fragilariopsis cylindrus* | fcy:FRACYDRAFT_276829 | M |
| Hacrobia | Prymnesiophyceae | *Emiliania huxleyi* | ehx:EMIHUDRAFT_458132 | M |
| Hacrobia | Prymnesiophyceae | *Emiliania huxleyi* | ehx:EMIHUDRAFT_456620 | M |
| Hacrobia | Prymnesiophyceae | *Emiliania huxleyi* | ehx:EMIHUDRAFT_449531 | M |
| Hacrobia | Prymnesiophyceae | *Emiliania huxleyi* | ehx:EMIHUDRAFT_247620 | M |
| Hacrobia | Cryptophyceae | *Guillardia theta* | gtt:GUITHDRAFT_92084 | M |
| Chlorophyta | Mamiellophyceae | *Micromonas pusilla* | mpp:MICPUCDRAFT_70850 | M |
| Chlorophyta | Mamiellophyceae | *Micromonas commoda* | mis:MICPUN_113551 | M |
| Chlorophyta | Chlorophyceae | *Monoraphidium neglectum* | mng:MNEG_3286 | M |
| Chlorophyta | Chlorophyceae | *Chlamydomonas reinhardtii* | cre:CHLRE_03g152900v5 | M |
| Chlorophyta | Chlorophyceae | *Chlamydomonas reinhardtii* | cre:CHLRE_13g589870v5 | M |
| Chlorophyta | Chlorophyceae | *Volvox carteri* | vcn:VOLCADRAFT_104564 | [38] |
| Chlorophyta | Chlorophyceae | *Chlorella variabilis* | cvr:CHLNCDRAFT_144467 | [39] |
| Chlorophyta | Trebouxiophyceae | *Coccomyxa subellipsoidea* | csl:COCSUDRAFT_44541 | M |

**Table S5. Psip1 genes.** Sequences were assigned based on sequence similarity to literature references[46] via protein BLAST search (expectation value cutoff E 10^-25^).

| **Clade** | **Class** | **Species** | **Gene** |
| --- | --- | --- | --- |
| Stramenopiles | Bacillariophyceae | *Phaeodactylum tricornutum* | pti:PHATR_33251 |
| Hacrobia | Prymnesiophyceae | *Emiliania huxleyi* | ehx:EMIHUDRAFT_458652 |
| Chlorophyta | Mamiellophyceae | *Bathycoccus prasinos* | bpg:Bathy01g02860 |
| Chlorophyta | Mamiellophyceae | *Ostreococcus tauri* | ota:OT_ostta19g00190 |
| Chlorophyta | Mamiellophyceae | *Ostreococcus tauri* | ota:OT_ostta08g00030 |
| Chlorophyta | Mamiellophyceae | *Ostreococcus tauri* | ota:OT_ostta19g00670 |

**Table S6.** Taxonomic information of all eukaryotic organisms transcribing *phoA* genes.

| **Taxonomy (Eukaryota)** |
| --- |
| Eukaryota;Alveolata;Ciliophora;Prostomatea;Prostomatea_X;Colepidae;Tiarina;Tiarina |
| Eukaryota;Alveolata;Dinophyta;Dinophyceae;Dinophyceae_X;Dinophyceae_XX;Gymnodinium;Gymnodinium |
| Eukaryota;Alveolata;Dinophyta;Dinophyceae;Dinophyceae_X;Dinophyceae_XX;Kryptoperidinium;Kryptoperidinium |
| Eukaryota;Alveolata;Dinophyta;Dinophyceae;Dinophyceae_X;Dinophyceae_XX;Lingulodinium;Lingulodinium |
| Eukaryota;Alveolata;Dinophyta;Dinophyceae;Dinophyceae_X;Dinophyceae_XX;Pyrodinium;Pyrodinium |
| Eukaryota;Alveolata;Dinophyta;Dinophyceae;Dinophyceae_X;Suessiales;Symbiodinium;Symbiodinium |
| Eukaryota;Amoebozoa;Conosa;Mycetozoa-Dictyostelea;Dictyosteliida;Dictyosteliida-Group-IV;Dictyostelium;Dictyostelium |
| Eukaryota;Archaeplastida;Chlorophyta;Chlorodendrophyceae;Chlorodendrales;Chlorodendrales_X;Tetraselmis;Tetraselmis |
| Eukaryota;Archaeplastida;Chlorophyta;Mamiellophyceae;Dolichomastigales;Crustomastigaceae;Crustomastix;Crustomastix |
| Eukaryota;Archaeplastida;Chlorophyta;Mamiellophyceae;Mamiellales;Mamiellaceae;Micromonas;Micromonas |
| Eukaryota;Archaeplastida;Chlorophyta;Prasinophyceae;unclassified |
| Eukaryota;Archaeplastida;Streptophyta;Embryophyceae;Embryophyceae_X;Embryophyceae_XX;Populus;Populus |
| Eukaryota;Hacrobia;Cryptophyta;Cryptophyceae;Cryptophyceae_X;Cryptomonadales;Geminigera;Geminigera |
| Eukaryota;Hacrobia;Haptophyta;Prymnesiophyceae;Coccolithales;Calcidiscaceae;Calcidiscus;Calcidiscus |
| Eukaryota;Hacrobia;Haptophyta;Prymnesiophyceae;Isochrysidales;Noelaerhabdaceae;Emiliania;Emiliania |
| Eukaryota;Hacrobia;Haptophyta;Prymnesiophyceae;Phaeocystales;Phaeocystaceae;Phaeocystis;Phaeocystis |
| Eukaryota;Hacrobia;Haptophyta;Prymnesiophyceae;Prymnesiales;Chrysochromulinaceae;Chrysochromulina;Chrysochromulina |
| Eukaryota;Hacrobia;Haptophyta;Prymnesiophyceae;Zygodiscales;Pontosphaeraceae;Scyphosphaera;Scyphosphaera |
| Eukaryota;Opisthokonta;Fungi;Ascomycota;Saccharomycotina;Saccharomycetales;Candida;Candida |
| Eukaryota;Opisthokonta;Metazoa;Annelida;Annelida_X;Annelida_XX;Helobdella;Helobdella |
| Eukaryota;Opisthokonta;Metazoa;Arthropoda;Chelicerata;Arachnida;Ixodes;Ixodes |
| Eukaryota;Opisthokonta;Metazoa;Arthropoda;Crustacea;Branchiopoda;Daphnia;Daphnia |
| Eukaryota;Opisthokonta;Metazoa;Arthropoda;Hexapoda;Insecta;Acyrthosiphon;Acyrthosiphon |
| Eukaryota;Opisthokonta;Metazoa;Arthropoda;Hexapoda;Insecta;Apis;Apis |
| Eukaryota;Opisthokonta;Metazoa;Arthropoda;Hexapoda;Insecta;Drosophila;Drosophila |
| Eukaryota;Opisthokonta;Metazoa;Arthropoda;Hexapoda;Insecta;Nasonia;Nasonia |
| Eukaryota;Opisthokonta;Metazoa;Arthropoda;Hexapoda;Insecta;Pediculus;Pediculus |
| Eukaryota;Opisthokonta;Metazoa;Arthropoda;Hexapoda;Insecta;Tribolium;Tribolium |
| Eukaryota;Opisthokonta;Metazoa;Cephalochordata;Cephalochordata_X;Cephalochordata_XX;Branchiostoma;Branchiostoma |
| Eukaryota;Opisthokonta;Metazoa;Cnidaria;Cnidaria_X;Anthozoa;Nematostella;Nematostella |
| Eukaryota;Opisthokonta;Metazoa;Craniata;Craniata_X;Amphibia;Xenopus;Xenopus |
| Eukaryota;Opisthokonta;Metazoa;Craniata;Craniata_X;Archosauria;Gallus;Gallus |
| Eukaryota;Opisthokonta;Metazoa;Craniata;Craniata_X;Archosauria;Meleagris;Meleagris |
| Eukaryota;Opisthokonta;Metazoa;Craniata;Craniata_X;Craniata_XX;Bos;Bos |
| Eukaryota;Opisthokonta;Metazoa;Craniata;Craniata_X;Craniata_XX;Cavia;Cavia |
| Eukaryota;Opisthokonta;Metazoa;Craniata;Craniata_X;Craniata_XX;Homo;Homo |
| Eukaryota;Opisthokonta;Metazoa;Craniata;Craniata_X;Craniata_XX;Rattus;Rattus |
| Eukaryota;Opisthokonta;Metazoa;Craniata;Craniata_X;Hyperoartia;Taeniopygia;Taeniopygia |
| Eukaryota;Opisthokonta;Metazoa;Craniata;Craniata_X;Mammalia;Monodelphis;Monodelphis |
| Eukaryota;Opisthokonta;Metazoa;Craniata;Craniata_X;Teleostei;Danio;Danio |
| Eukaryota;Opisthokonta;Metazoa;Craniata;Craniata_X;Teleostei;Takifugu;Takifugu |
| Eukaryota;Opisthokonta;Metazoa;Ctenophora;Ctenophora_X;Ctenophora_XX;Undescribed;Undescribed |
| Eukaryota;Opisthokonta;Metazoa;Echinodermata;Echinodermata_X;Echinodermata_XX;Strongylocentrotus;Strongylocentrotus |
| Eukaryota;Opisthokonta;Metazoa;Mollusca;Gastropoda;Patellogastropoda;Lottia;Lottia |
| Eukaryota;Opisthokonta;Metazoa;Placozoa;Placozoa_X;Placozoa_XX;Trichoplax;Trichoplax |
| Eukaryota;Opisthokonta;Metazoa;Urochordata;Urochordata_X;Ascidiacea;Ciona;Ciona |
| Eukaryota;Rhizaria;Foraminifera;Miliolida;Miliolida_X;Soritidae;Sorites;Sorites |
| Eukaryota;Rhizaria;Foraminifera;Rotaliida;Discorbacea;Rosalinidae;Rosalina;Rosalina |
| Eukaryota;Rhizaria;Foraminifera;Rotaliida;Rotaliida_X;Elphidiidae;Elphidium;Elphidium |
| Eukaryota;Rhizaria;Foraminifera;Rotaliida;Rotaliida_X;Rotaliida_XX;Ammonia;Ammonia |
| Eukaryota;Stramenopiles;Stramenopiles_X;Bacillariophyta;Bacillariophyta_X;Araphid-pennate;Staurosira;Staurosira |
| Eukaryota;Stramenopiles;Stramenopiles_X;Bacillariophyta;Bacillariophyta_X;Polar-centric-Mediophyceae;Chaetoceros;Chaetoceros |
| Eukaryota;Stramenopiles;Stramenopiles_X;Bacillariophyta;Bacillariophyta_X;Polar-centric-Mediophyceae;Thalassiosira;Thalassiosira |
| Eukaryota;Stramenopiles;Stramenopiles_X;Bacillariophyta;Bacillariophyta_X;Raphid-pennate;Fragilariopsis;Fragilariopsis |
| Eukaryota;Stramenopiles;Stramenopiles_X;Bacillariophyta;Bacillariophyta_X;Raphid-pennate;Nitzschia;Nitzschia |
| Eukaryota;Stramenopiles;Stramenopiles_X;Dictyochophyceae;Dictyochophyceae_X;Dictyochales;Dictyocha;Dictyocha |
| Eukaryota;Stramenopiles;Stramenopiles_X;Oomyceta;Oomyceta_X;Oomyceta_XX;Phytophthora;Phytophthora |

**Table S7. Differential expression of transcripts related to FeS biogenesis and photosynthesis in diatoms within incubation reactions at station 4 (Offshore).**

| **KEGG Ortho-logy** | **Gene** | **Description** | **+Fe vs. Ctrl** | | **+N vs. Ctrl** | | **+FeN vs. Ctrl** | | **Role** |
| --- | --- | --- | --- | --- | --- | --- | --- | --- | --- |
|  |  |  | **Fold-change** | ***p*-value** | **Fold-change** | ***p*-value** | **Fold-change** | ***p*-value** |  |
| K00858 | *ppnK* | NAD+ kinase | 1.327 | *0.0004* | -0.213 | 1.0000 | 0.126 | 1.0000 | **NADPH production** |
| K22074 | NFU1 | Fe-S cluster scaffold, mitochondrial | 0.815 | *0.0112* | 0.016 | 1.0000 | 0.199 | 0.8233 | **FeS biogenesis** |
| K09015 | *sufD* | Fe-S cluster assembly protein D | 0.857 | *0.0300* | -0.576 | 0.2877 | 0.888 | *0.0177* |  |
| K08910 | LHCA4 | light-harvesting complex I chlorophyll a/b binding protein 4 | 1.022 | *0.0058* | 0.043 | 1.0000 | 1.853 | >*0.0001* | **Photo-**  **synthesis** |
| K18010 | HCAR | 7-hydroxymethyl chlorophyll *a* reductase, [FeS] | 1.824 | *0.0300* | 0.025 | 1.0000 | 1.766 | *0.0290* |  |

**Table S8. Presence and regulation of phosphatase gene expression by major phytoplankton groups.** Presence of phosphatase gene transcription (**blue**) is indicated by Y = yes, N = no. Regulation of gene expression in response to natural gradients of ambient nutrient availability (**green**) and artificial enhancement of nutrient availability in incubations (**pink**) is indicated by + = present, - = absent. Regulation was deemed present when adjusted *p*-value < 0.05 (DESeq2 Wald test, Benjamini & Hochberg) between any stations or any treatments, respectively. Gene abbreviations are described in the main manuscript, **Figure 2**. For brevity, 5’-NT = 5’-NT (ecto) or 5’-NT (cyclic); NUDIX = *nudH* or *mutT*.

| **Group** | AP_aty_ | *phoA* | *phoD* | *phoX* | *psip1* | *ugpQ* | PLC-PDE | PDE_nc_ | 5’-NT | ppa | ppx | NUDIX | *phnY* | *phnZ* |
| --- | --- | --- | --- | --- | --- | --- | --- | --- | --- | --- | --- | --- | --- | --- |
| Total Community | **Y** | **Y** | **Y** | **Y** | **Y** | **Y** | **Y** | **Y** | **Y** | **Y** | **Y** | **Y** | **Y** | **Y** |
|  | **+** | **+** | **+** | **+** | **+** | **+** | **+** | **+** | **+** | **+** | - | **+** | **+** | - |
|  | **+** | **+** | **+** | **+** | **+** | **+** | - | **+** | **+** | **+** | **+** | **+** | - | - |
| *Dinophyta* | **Y** | **Y** | **Y** | **Y** | **Y** | **Y** | **Y** | **Y** | **Y** | **Y** | **Y** | **Y** | **Y** | **Y** |
|  | **+** | **+** | **+** | **+** | **+** | - | **+** | **+** | - | **+** | **+** | **+** | **+** | - |
|  | **+** | **+** | - | - | **+** | **+** | - | **+** | **+** | **+** | - | - | - | - |
| *Bacillario-phyta* | **Y** | **Y** | **Y** | **Y** | **Y** | **Y** | **Y** | **Y** | **Y** | **Y** | **Y** | **Y** | **Y** | **Y** |
|  | **+** | **+** | **+** | - | - | **+** | - | **+** | - | **+** | - | - | **+** | - |
|  | **+** | **+** | **+** | **+** | **+** | - | - | **+** | - | - | **+** | **+** | - | - |
| *Mamiello-phyceae* | N | **Y** | **Y** | N | **Y** | **Y** | **Y** | **Y** | **Y** | **Y** | **Y** | **Y** | N | N |
|  |  | **+** | **+** |  | **+** | **+** | **+** | **+** | - | **+** | **+** | **+** |  |  |
|  |  | - | **+** |  | - | **+** | **+** | **+** | **+** | - | - | **+** |  |  |
| *Prymnesio-phyceae* | **Y** | **Y** | **Y** | **Y** | **Y** | **Y** | **Y** | **Y** | **Y** | **Y** | **Y** | **Y** | **Y** | **Y** |
|  | **+** | - | **+** | - | - | **+** | **+** | **+** | - | **+** | **+** | **+** | **+** | - |
|  | - | - | - | - | **+** | **+** | - | **+** | - | **+** | - | **+** | - | - |
| *Pelago-monas* | **Y** | N | **Y** | N | N | **Y** | N | **Y** | **Y** | **Y** | **Y** | **Y** | **Y** |  |
|  | - |  | - |  |  | **+** |  | **+** | - | **+** | **+** | **+** | - | N |
|  | - |  | - |  |  | - |  | **+** | - | **+** | **+** | - | **+** |  |

**Table S9. Percentile ranks of selected phosphatase and iron stress marker genes among all transcribed community KOs.**

| **Gene** | **Station 2** | | | | **Station 4** | | | |
| --- | --- | --- | --- | --- | --- | --- | --- | --- |
|  | **Ctrl** | **+Fe** | **+N** | **+FeN** | **+Ctrl** | **+Fe** | **+N** | **+FeN** |
| **AP_aty_** | 88.0 | 90.5 | 96.7 | 92.7 | 59.8 | 62.2 | 59.5 | 61.2 |
| **PDE_nc_** | 95.5 | 96.5 | 94.5 | 96.9 | 98.8 | 98.6 | 98.8 | 99.0 |
| ***phoA*** | 79.9 | 74.7 | 85.3 | 80.6 | 51.7 | 45.9 | 39.6 | 48.7 |
| ***phoD*** | 86.2 | 86.0 | 84.7 | 86.6 | 73.6 | 72.9 | 74.4 | 73.6 |
| ***phoX*** | 68.2 | 69.3 | 62.8 | 77.9 | 58.7 | 61.2 | 59.6 | 56.9 |
| **ISIP1** | 98.5 | 87.6 | 98.4 | 91.9 | 61.3 | 55.0 | 67.8 | 55.4 |
| **ISIP2a** | 99.9 | 98.7 | 99.9 | 98.8 | 97.7 | 95.3 | 98.4 | 95.4 |
| **ISIP3** | 98.6 | 91.1 | 99.3 | 94.3 | 97.1 | 96.4 | 96.2 | 95.6 |

**References**

1. Strickland JDH, Parsons TR. *A Practical Handbook of Seawater Analysis*. 2 ed. vol 167. Canadian Bulletin of Fisheries and Aquatic Sciences. Fisheries Research Board of Canada; 1972:310.

2. Diaz JM *et al.* Dissolved organic phosphorus utilization by phytoplankton reveals preferential degradation of polyphosphates over phosphomonoesters. *Front Mar Sci*. 2018;5(380):1-17. doi:10.3389/fmars.2018.00380

3. Hansen H, Koroleff F. *Determination of Nutrients*. 3 ed. Methods of Seawater Analysis. WILEY-VCH; 1999:159-228.

4. Monaghan EJ, Ruttenberg KC. Dissolved organic phosphorus in the coastal ocean: Reassessment of available methods and seasonal phosphorus profiles from the Eel River Shelf. *Limnol Oceanogr*. 1999;44(7):1702-14. doi:10.4319/lo.1999.44.7.1702

5. Schlitzer R. Ocean Data View. 2025;doi:<https://odv.awi.de>

6. CalCOFI Methods Manual. California Cooperative Oceanic Fisheries Investigations. Accessed Mar 06, 2025. <https://calcofi.info/index.php/ccpublications/calcofi-methods>

7. Hogle SL *et al.* Pervasive iron limitation at subsurface chlorophyll maxima of the California Current. *Proc Natl Acad Sci*. 2018;115(52):13300-5. doi:10.1073/pnas.1813192115

8. Obata H, Karatani H, Nakayama E. Automated determination of iron in seawater by chelating resin concentration and chemiluminescence detection. *Anal Chem*. 1993;65(11):1524-8. doi:10.1021/ac00059a007

9. Lohan MC, Aguilar-Islas AM, Bruland KW. Direct determination of iron in acidified (pH 1.7) seawater samples by flow injection analysis with catalytic spectrophotometric detection: Application and intercomparison. *Limnol Oceanogr: Methods*. 2006;4(6):164-71. doi:10.4319/lom.2006.4.164

10. Mahaffey C *et al.* Alkaline phosphatase activity in the subtropical ocean: Insights from nutrient, dust and trace metal addition experiments. *Front Mar Sci*. 2014;1(73):1-13. doi:10.3389/fmars.2014.00073

11. Lampe RH *et al.* Short-term acidification promotes diverse iron acquisition and conservation mechanisms in upwelling-associated phytoplankton. *Nat Commun*. 2023;14(1):7215. doi:10.1038/s41467-023-42949-1

12. Rabines A, Lampe RH, Allen AE. Sterivex RNA extraction. 2020:Accessed Mar 06 2025. doi:10.17504/protocols.io.bd9ti96n

13. Bolger AM, Lohse M, Usadel B. Trimmomatic: a flexible trimmer for Illumina sequence data. *Bioinformatics*. 2014;30(15):2114-20. doi:10.1093/bioinformatics/btu170

14. Ewels P *et al.* MultiQC: summarize analysis results for multiple tools and samples in a single report. *Bioinformatics*. 2016;32(19):3047-8. doi:10.1093/bioinformatics/btw354

15. Bushmanova E *et al.* rnaSPAdes: a de novo transcriptome assembler and its application to RNA-Seq data. *GigaScience*. 2019;8(9):giz100. doi:10.1093/gigascience/giz100

16. Li D *et al.* MEGAHIT: an ultra-fast single-node solution for large and complex metagenomics assembly via succinct de Bruijn graph. *Bioinformatics*. 2015;31(10):1674-6. doi:10.1093/bioinformatics/btv033

17. Tang S, Lomsadze A, Borodovsky M. Identification of protein coding regions in RNA transcripts. *Nucleic Acids Res*. 2015;43(12):e78. doi:10.1093/nar/gkv227

18. Langmead B, Salzberg SL. Fast gapped-read alignment with Bowtie 2. *Nat Methods*. 2012;9(4):357-9. doi:10.1038/nmeth.1923

19. Bertrand EM *et al.* Phytoplankton–bacterial interactions mediate micronutrient colimitation at the coastal Antarctic sea ice edge. *Proc Natl Acad Sci*. 2015;112(32):9938-43. doi:10.1073/pnas.1501615112

20. Buchfink B, Xie C, Huson DH. Fast and sensitive protein alignment using DIAMOND. *Nat Methods*. 2015;12(1):59-60. doi:10.1038/nmeth.3176

21. Podell S, Gaasterland T. DarkHorse: a method for genome-wide prediction of horizontal gene transfer. *Genome Biol*. 2007;8(2):R16. doi:10.1186/gb-2007-8-2-r16

22. Kanehisa M *et al.* KEGG: new perspectives on genomes, pathways, diseases and drugs. *Nucleic Acids Res*. 2016;45(D1):D353-D61. doi:10.1093/nar/gkw1092

23. Aramaki T *et al.* KofamKOALA: KEGG Ortholog assignment based on profile HMM and adaptive score threshold. *Bioinformatics*. 2019;36(7):2251-2. doi:10.1093/bioinformatics/btz859

24. Granzow BN *et al.* A sensitive fluorescent assay for measuring carbon-phosphorus lyase activity in aquatic systems. *Limnol Oceanogr: Methods*. 2021;19(4):235-44. doi:10.1002/lom3.10418

25. Acker M *et al.* Phosphonate production by marine microbes: Exploring new sources and potential function. *Proc Natl Acad Sci*. 2022;119(11):e2113386119. doi:10.1073/pnas.2113386119

26. Sosa OA *et al.* Phosphate-limited ocean regions select for bacterial populations enriched in the carbon–phosphorus lyase pathway for phosphonate degradation. *Environ Microbiol*. 2019;21(7):2402-14. doi:10.1111/1462-2920.14628

27. Waggoner EM *et al.* Dissolved organic phosphorus bond-class utilization by *Synechococcus*. *FEMS Microbiol Ecol*. 2024;100(9):fiae099. doi:10.1093/femsec/fiae099

28. Davis CE, Mahaffey C. Elevated alkaline phosphatase activity in a phosphate-replete environment: Influence of sinking particles. *Limnol Oceanogr*. 2017;62(6):2389-403. doi:10.1002/lno.10572

29. Lidbury IDEA *et al.* A widely distributed phosphate-insensitive phosphatase presents a route for rapid organophosphorus remineralization in the biosphere. *Proc Natl Acad Sci*. 2022;119(5):e2118122119. doi:10.1073/pnas.2118122119

30. Thomson B *et al.* Resolving the paradox: Continuous cell-free alkaline phosphatase activity despite high phosphate concentrations. *Mar Chem*. 2019;214:103671. doi:10.1016/j.marchem.2019.103671

31. Davis C *et al.* Diurnal variability in alkaline phosphatase activity and the potential role of zooplankton. *Limnol Oceanogr Lett*. 2019;4(3):71-8. doi:10.1002/lol2.10104

32. Behnke J, LaRoche J. Iron uptake proteins in algae and the role of Iron Starvation-Induced Proteins (ISIPs). *Eur J Phycol*. 2020;55(3):339-60. doi:10.1080/09670262.2020.1744039

33. Kellogg RM *et al.* Adaptive responses of marine diatoms to zinc scarcity and ecological implications. *Nat Commun*. 2022;13(1):1995. doi:10.1038/s41467-022-29603-y

34. Dyhrman ST *et al.* The transcriptome and proteome of the diatom *Thalassiosira pseudonana* reveal a diverse phosphorus stress response. *PLoS One*. 2012;7(3):e33768. doi:10.1371/journal.pone.0033768

35. Chen X-H *et al.* Quantitative proteomics reveals common and specific Responses of a marine diatom *Thalassiosira pseudonana* to different macronutrient deficiencies. *Front Microbiol*. 2018;9(2761):2761. doi:10.3389/fmicb.2018.02761

36. Shih C-Y, Kang L-K, Chang J. Transcriptional responses to phosphorus stress in the marine diatom, Chaetoceros affinis, reveal characteristic genes and expression patterns in phosphorus uptake and intracellular recycling. *J Exp Mar Biol Ecol*. 2015;470:43-54. doi:10.1016/j.jembe.2015.05.001

37. Dell’Aquila G *et al.* Mobilization and cellular distribution of phosphate in the diatom *Phaeodactylum tricornutum*. *Front Plant Sci*. 2020;11:579. doi:10.3389/fpls.2020.00579

38. Hallmann A. Enzymes in the extracellular matrix of *Volvox*: an inducible, calcium-dependent phosphatase with a modular composition. *J Biol Chem*. 1999;274(3):1691-7. doi:10.1074/jbc.274.3.1691

39. Quisel JD, Wykoff DD, Grossman AR. Biochemical characterization of the extracellular phosphatases produced by phosphorus-deprived *Chlamydomonas reinhardtii*. *Plant Physiol*. 1996;111(3):839-48. doi:10.1104/pp.111.3.839

40. Guo J *et al.* Specialized proteomic responses and an ancient photoprotection mechanism sustain marine green algal growth during phosphate limitation. *Nat Microbiol*. 2018;3(7):781-90. doi:10.1038/s41564-018-0178-7

41. Xu Y *et al.* A novel alkalie phosphatase in the coccolithophore *Emiliania huxleyi* (prymnesiophyceae) and its regulation by by phosphorus. *J Phycol*. 2006;42(4):835-44. doi:10.1111/j.1529-8817.2006.00243.x

42. Lin X *et al.* Rapidly diverging evolution of an atypical alkaline phosphatase (PhoAaty) in marine phytoplankton: insights from dinoflagellate alkaline phosphatases. *Front Microbiol*. 2015;6:868. doi:10.3389/fmicb.2015.00868

43. Wurch LL *et al.* Proteome changes driven by phosphorus deficiency and recovery in the brown tide-forming alga *Aureococcus anophagefferens*. *PLoS One*. 2011;6(12):e28949. doi:10.1371/journal.pone.0028949

44. Li T *et al.* Identification and expression analysis of an atypical alkaline phosphatase in *Emiliania huxleyi*. *Front Microbiol*. 2018;9:2156. doi:10.3389/fmicb.2018.02156

45. Alipanah L *et al.* Molecular adaptations to phosphorus deprivation and comparison with nitrogen deprivation responses in the diatom *Phaeodactylum tricornutum*. *PLoS One*. 2018;13(2):e0193335. doi:10.1371/journal.pone.0193335

46. Torcello-Requena A *et al.* A distinct, high-affinity, alkaline phosphatase facilitates occupation of P-depleted environments by marine picocyanobacteria. *Proc Natl Acad Sci*. 2024;121(20):e2312892121. doi:10.1073/pnas.2312892121
